# *MSP1* encodes an essential RNA-binding PPR factor required for *nad1* maturation and complex I biogenesis in *Arabidopsis* mitochondria

**DOI:** 10.1101/2022.11.12.516219

**Authors:** Corinne Best, Ron Mizrahi, Rana Edris, Hui Tang, Hagit Zer, Catherine Colas des Francs-Small, Omri M. Finkel, Hongliang Zhu, Ian D. Small, Oren Ostersetzer-Biran

## Abstract

**Summary:** Mitochondria are semi-autonomous organelles that serve as hubs for aerobic energy metabolism. The biogenesis of the respiratory (OXPHOS) system relies on nuclear-encoded factors, which regulate the transcription, processing and translation of mitochondrial (mt)RNAs. These include proteins of primordial origin, as well as eukaryotic-type RNA-binding families recruited from the host genomes to fu<u>nc</u>tion in mitogenome expression. Pentatricopeptide repeat (PPR) proteins constitute a major gene-family in angiosperms that is pivotal in many aspects of mtRNA metabolism, such as editing, splicing or stability. Here, we report the analysis of *MITOCHONDRIA STABILITY/PROCESSING PPR FACTOR1* (*MSP1*, At4g20090), a canonical mitochondria-localized PPR protein that is necessary for mitochondrial biogenesis and embryo-development. Functional complementation confirmed that the phenotypes result from a disruption of the *MSP1* gene. As a loss-of-function allele of *Arabidopsis MSP1* leads to seed abortion, we employed an embryo-rescue method for the molecular characterization of *msp1* mutants. Our data show that *msp1* embryo-development fails to proceed beyond the heart-torpedo transition stage as a consequence of a severe nad1 pre-RNA processing-defect, resulting in the loss of respiratory complex I (CI) activity. The maturation of *nad1* involves the processing of three RNA-fragments, *nad1*.*1, nad1*.*2* and *nad1*.*3*. Based on biochemical analyses and the mtRNA profiles in wild-type and *msp1* plants, we concluded that through its association with a specific site in *nad1*.*1*, MSP1 facilitates the generation of its 3’-terminus and stabilizes it -a prerequisite for *nad1* exons a-b splicing. Our data substantiate the importance of mtRNA metabolism for the biogenesis of the respiratory machinery during early-plant development.

## Introduction

Mitochondria serve as hubs for cellular metabolism. In non-photosynthetic cells, they generate most of the energy used in the biochemical reactions required to sustain growth and development. As descendants of a symbiotic bacterium, most mitochondria harbor an intrinsic genetic system (mtDNA, mitogenome) to make their own RNAs and proteins. Nevertheless, the majority of the organellar proteins and some tRNAs are encoded in the nucleus and imported into the organelle. The biogenesis of the OXPHOS system in plants is a multifaceted process, which relies on complex regulatory mechanisms to allow the stoichiometric accumulation of subunits encoded by the two separate genome machineries (Schwarzländer *et al*., 2012; Best, C. *et al*., 2020). Despite their universal role in eukaryotic cells, various important aspects distinguish the mitogenomes of land plants from those in animals. While the mtDNA in mammals, or humans, underwent extensive compaction (to ∼17 kb), their counterparts in angiosperms are notably larger in size (168 up to 11,000 kb) and much more complex in structure (Gualberto & Newton, 2017). Accumulating data suggest that mitochondrial (mt)RNA processing plays a pivotal role in the regulation of plants’ mitochondria gene-expression (Best, C. *et al*., 2020). For example, many essential genes in angiosperm mitochondria harbor introns, which must be excised post-transcriptionally from the pre-mRNAs to allow the integration of exons into functional reading-frames (Zmudjak & Ostersetzer-Biran, 2017). These are primarily found in NADH-dehydrogenase (CI) genes, and also interrupt the coding-sequences of cytochrome *c* oxidase (CIV), cytochrome *c* biogenesis (CCM) factors and ribosomal protein-coding genes. These are mainly group II introns (Bonen, 2008), an ancient class of catalytic RNAs that are evolutionarily related to spliceosomal-RNAs, non-LTR retrotransposons or telomerases (Lambowitz & Belfort, 2015). It is therefore speculated that the symbiont introns had a major role in shaping the genomic organization of terrestrial life-forms, and may have also been determining factors in the rise of the nuclear envelope (Martin & Koonin, 2006), in order to separate transcription from translation.

Group II introns are characterized by six domains (D1-D6) based on secondary structure, which adopt a complex tertiary structure mediated by long-range tertiary interactions (Zhao & Pyle, 2017). Canonical introns in this class are able to catalyze their own excision in-vitro, but their activities in-vivo rely on intron-encoded proteins (maturases, MATs), which assist in the folding of their cognate introns into the catalytically-active form (Zhao & Pyle, 2017; Mizrahi *et al*., 2022). The mitochondrial introns in angiosperms have diverged considerably in sequence and have usually also lost their cognate maturase-ORFs (Guo & Mower, 2013; Schmitz-Linneweber *et al*., 2015). Some are fragmented, requiring the intron parts to be transcribed independently and then to associate non-covalently allowing splicing in ‘*trans*’ (Bonen, 2008). During evolution, land plants have acquired many host factors to assist in the splicing of their highly derived organellar introns (Brown *et al*., 2014; Zmudjak & Ostersetzer-Biran, 2017). A few are related to maturases (Pfam-01348) (Mizrahi *et al*., 2022), but most belong to different protein families, such as RNA-helicases (pfam-00270), mTERFs (Pfam-02536), PORR (Pfam-11955), or PPR (Pfam-13812) proteins (see e.g., (Colas des Francs-Small & Small, 2014; Zmudjak & Ostersetzer-Biran, 2017). Currently there is very little mechanistic information about the molecular functions of these factors, specifically how they facilitate the processing of their RNA-ligands.

PPR proteins comprise one of the largest known protein families in angiosperms, with ∼450 members in *Arabidopsis* (Small & Peeters, 2000). These contain between 2 and 30 repeats of a 35-amino-acid domain, which adopts an ‘antiparallel’ helical form (Gully *et al*., 2015; Shen *et al*., 2015), and can be classified according to variations in their PPR motifs (Lurin *et al*., 2004). Most plant PPRs contain N-terminal signals that target them to the ‘energy-producing’ organelles, where they affect diverse RNA-processing steps, including stability, trimming, editing and splicing (Barkan & Small, 2014). These act as sequence-specific ssRNA-binding proteins, where the RNA winds around the protein, by a mechanism like that by which transcription activator-like effectors (TALEs) bind dsDNA (Barkan *et al*., 2012; Yagi *et al*., 2013). In this work, we elucidate the roles of a P-class PPR protein encoded by the *AT4G20090* gene. In accordance with its role in mtRNA-processing, we have renamed this gene *<u>M</u>ITOCHONDRIA <u>S</u>TABILITY/<u>P</u>ROCESSING PPR FACTOR1* (*MSP1*). By studying the physiology of previously unattainable *msp1* knockout-lines, we demonstrate the effects of MSP1 on mitochondria gene-expression and function.

## Materials and Methods

A complete version of the Materials and Methods is available in the supplemental information.

### Plant material and growth conditions

*Arabidopsis thaliana* was used in all experiments. Wild-type (Col-0) and *mps1* mutant SALK-lines were obtained from ABRC. The growth conditions used for the analysis of wild-type and mutant lines were identical to those described previously (Shevtsov-Tal *et al*., 2021).

### GFP localization assays

The N-terminal region (300 bps) of *MSP1* gene was fused in-frame to GFP and cloned into a modified pCAMBIA binary-vector (gift of Dr. Yoram Eyal, ARO). *In-vivo* protein localization of GFP-fusion proteins were performed via root transformation in tissue culture, similarly to the method described by Weigel and Glazebrook (2006).

### Embryo rescue and establishment of homozygous *MSP1* mutants

The development of *msp1* mutant-line embryos was performed essentially as described previously (Shevtsov-Tal *et al*., 2021), using white seeds obtained from green mature siliques of wild-type or heterozygous *msp1-1* plants.

### Plant transformation and functional complementation of the *MSP1-1* mutation

A 1,989 bp fragment containing the coding region of *MSP1* was generated by PCR, cloned under the control of the 35S promoter region within pCAMBIA-2300 vector, and used to transform *msp1-1* plants by a floral-dip method.

### Microscopic and macroscopic analyses of *Arabidopsis* wild-type and mutant plants

Analysis of whole plant morphology, roots, leaves, siliques and seeds of wild-type and mutant lines were examined under a stereoscopic (dissecting) and light microscopes.

### RNA extraction and analysis

RNA extraction and analysis were performed essentially as described previously (Sultan *et al*., 2016; Shevtsov-Tal *et al*., 2021). RNA-sequencing was carried out with HTS-Illumina platform, used for a paired-end 150-bp sequencing strategy.

### Preparation of recombinant MSP1 protein

A g-Block fragment (Synthezza, Jerusalem, Israel) containing the MSP1s’ PPRs 4-14 motifs was cloned into pGEX-4T1, in-frame to GST in its N-terminus and a C-terminal 6xHis tag. The recombinant protein (rMSP1) was expressed in *E. coli*, and purified by affinity chromatography.

### *In-vitro* RNA binding assays and PNPase-protection assays

Filter-binding assays were performed with a slot-blot manifold, essentially as described previously (Ostersetzer *et al*., 2005; Keren *et al*., 2008). Nuclease-protection assays were perform essentially as described by Prikryl *et al*. (2011), using PNPase, Benzonase or RNase A in the presence or absence of rMSP1.^32^P-labeled RNAs were denatured and snap cooled on ice, before was used in the reaction assays. The samples were denatured in 60% formamide sample-buffer and resolved on 10% AA/7M-urea gels, in 1×TBE buffer.

### Enriched mitochondrial membrane preparation and analysis

BN-PAGE was performed according to the method described previously (Pineau *et al*., 2008). Crude mitochondria were solubilized with 5% (w/v) digitonin and loaded onto a native 4-16% linear gradient gel. For immunoblotting, the gel was transferred to a PVDF membrane. CI activity assays were performed essentially as described previously (Eubel *et al*., 2005).

## Results

### *AT4G20090* encodes a canonical P-class PPR protein

Considering the importance of mtRNA-metabolism for embryo-development and plant physiology, we surveyed the *Arabidopsis* SeedGene/EMB databases (Meinke, 2020), which include ∼2,200 embryogenesis mutants. Among these, 35 are identified as PPRs, 19 of which are known or predicted to reside in the mitochondria (Table 1). These include APPR6 (Manavski *et al*., 2012), COD1 (Dahan *et al*., 2014), DEK36 (Wang *et al*., 2017) and OTP43 (Falcon de Longevialle *et al*., 2007) that function in mtRNA-processing. Here, we report the characterization of MSP1/EMB1025, a P-class PPR whose functions are essential during early embryogenesis.

**Table 1.** List of *Arabidopsis* PPR-related *emb* mutants with predicted or established roles in mitogenome expression or mtRNA-metabolism.

| Gene I.D. | Gene symbol | PPR group | Cellular location<sup>*1</sup> | Gene function | References |
| --- | --- | --- | --- | --- | --- |
| **At4g20090** | ***MSP1/EMB1025*** | **P** | **mit** | **Altered *nad1.1* processing** | **This study** |
| At1g05600 | *EMB3101* | P | (mit) | n.d.<sup>*2</sup> |  |
| At1g10270 | *GRP23* | P | nucl/mit | nuclear transcription (interacts with RNA pol II), mito's functions? | (Ding *et al.*, 2006; Narsai *et al.*, 2011; Senkler *et al.*, 2017) |
| At1g74900 | *OTP43* | P | mit | splicing of *nad1* i1 | (Falcon de Longevialle *et al.*, 2007) |
| At1g77360 | *APPR6* | P | mit | 5'-end processing, translational initiation of *rps3* RNA | (Manavski *et al.*, 2012) |
| At1g79490 | *EMB2217* | P | (mit) | n.d. | (Narsai *et al.*, 2011) |
| At2g01390 | *EMB3111* | P | (mit) | n.d. | (Narsai *et al.*, 2011) |
| At2g02150 | *EMB2794* | P | (mit/clp) | n.d. |  |
| At2g15690 | *DYW2* | DYW | (mit) | Provides the missing cytidine deaminase activity to the E+ PPR proteins | (Guillaumot *et al.*, 2017) |
| At2g32230 | *PRORP1* | P | mit/clp | RNase P-like (5' cleavage of pre-tRNAs) | (Gobert *et al.*, 2010) |
| At2g35030 | *COD1* | E+ | mit | mitochondrial RNA editing | (Dahan *et al.*, 2014) |
| At3g22670 | *EMP10* | P | (mit) | homolog of ZmEMP10 (putative splicing factor) | (Cai *et al.*, 2017) |
| At3g49240 | *NUWA* | P | mit | May bridge and stabilize the interaction between PPR-E+ and DYW2 proteins, or regulates mtRNA translation | (He *et al.*, 2017) |
| At4g21190 | *EMB1417* | P | (mit) | n.d. |  |
| At4g21300 | *DEK36* | E+ | mit | mitochondrial RNA editing | (Wang *et al.*, 2017) |
| At4g33990 | *EMB2758* | DYW | (mit) | n.d. |  |
| At5g39680 | *EMB2744* | DYW | (mit) | n.d. |  |
| At5g39710 | *EMB2745* | P | (mit) | n.d. (regulated by miR400?) | (Li *et al.*, 2014) |
| At1g13800 | *FAC19* | P | (mit) | possibly RNA editing | (Yu *et al.*, 2012) |

<sup>\*1</sup> - quotes indicate to predicted intracellular locations by the ExPASy portal (https://www.expasy.org) and the SUBAcon server (Hooper *et al.*, 2017) (mitochondrial, mit; chloroplast, clp; nucleus, nucl; cytosolic, cyto).
<sup>\*2</sup> - n.d., not determined.

Gene-expression databases (Hruz *et al*., 2008; Berardini *et al*., 2015) indicate that *MSP1* (Fig. 1A) is a lowly-expressed gene, showing most expression in embryonic and young-developing tissues (Fig. S1). Prediction of conserved motifs indicate that MSP1 (660 aa) harbors at least 11 PPR motifs (Fig. 1A), with a suggested topology of: NH_2_-76aa-(P)-(P)-(P)-P-P-P-P-P-P-P-P-P-P-P-(P)-50aa-COOH, with lower scores for the first three and last PPR motifs (Figs. 1A, S2A). The AlphaFold algorithm (Jumper *et al*., 2021) suggests a typical PPR-fold for MSP1 (Fig. S2B), with right-handed two-turn superhelical structure. Based on this model, the 11 well-supported PPR motifs of MSP1 are bounded by an undefined structure at the N’-terminus, which comprises of the 36/37 amino-acid organellar localization signal and the first three PPR-like motifs, and a low confidence region within the C-terminus that may contain an additional PPR motif (Fig. S2B).While the PPR domains confer sequence-specific RNA recognition, the low complexity regions may contribute to protein-protein or protein-ligand interactions, as also shown for the superhelix TPR or TALE repeat proteins (Grove *et al*., 2008; Deng *et al*., 2012). Like PPR10 (Yin *et al*., 2013; Gully *et al*., 2015), the internal core of MSP1 harbors a basic surface (Figs. S2B), which is postulated to represent an RNA-binding path. However, functional analyses were required to test the predicted RNA-binding characteristics of MSP1.

**Figure 1.**
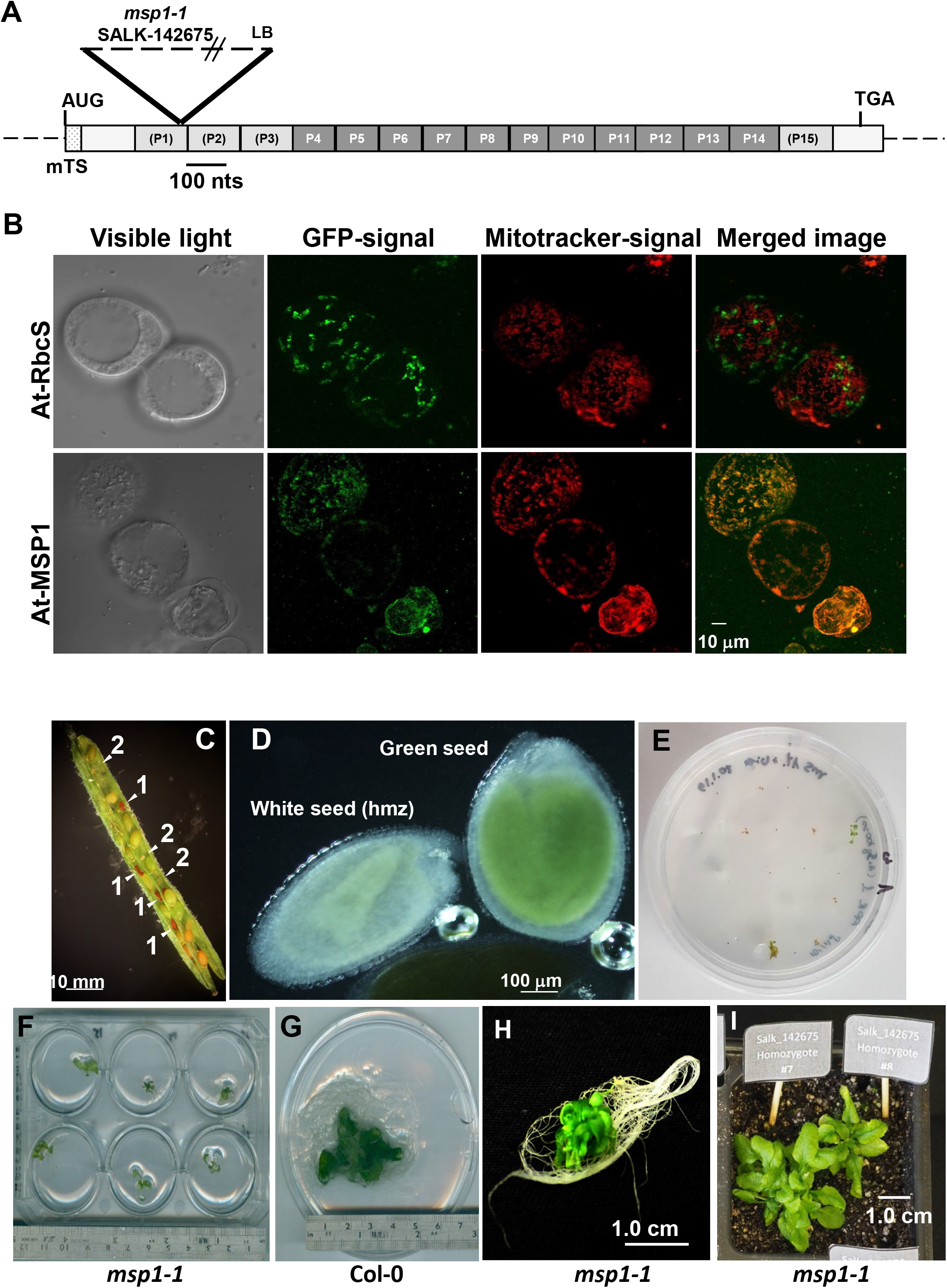
*AT4G20090* gene product encodes an essential mitochondria-localized PPR protein. (A) Scheme of *MSP1* (*AT4G20090*) gene structure. In silico analysis suggest the presence of at least 11 repeats (P4 to P14) of the conserved P-class PPR motif, and putative additional 3 PPRs (P1 to P3) at the N-terminus and the PPR domain P15 at the C-terminus. The positions of the predicted mitochondrial targeting signal (mTS) and the T-DNA insertion in SALK line 142675 (*msp1-1*) within the coding sequence of *MSP1* are outlined above the sequence. (B) GFP localization analyses. *Arabidopsis* root tissue culture cells were agroinfiltrated with pCAMBIA 2300 vectors containing GFP sequence fused in-frame to the plastidial Rubisco small subunit (*RBCS, AT1G67090*) and *MSP1* (*AT4G20090*) coding sequences, under the control of the *UBIQUTIN-10* (*AT4G05320*) promoter (Grefen *et al*., 2010). The figure shows bright-field images (grayscale) and confocal GFP signals (green), MitoTracker marker (red) and overlain GFP + MitoTracker images obtained with each construct. (C) Mature siliques obtained from heterozygous *msp1-1* (SALK-142675) lines. Arrows indicate homozygous mutant seeds. Numbers indicate seeds which contain embryos arrested at the late heart or torpedo stages (“1”), or aborted seeds with dead embryos (“2”). (D) White seeds (containing homozygous *msp1* embryos) and green seeds (containing heterozygous or wild-type embryos) obtained from the immature siliques of heterozygous *msp1-1* plants. Images taken with differential interference contrast (Nomarski) microscopy. (E) Seeds collected under sterile conditions from surface sterilized immature siliques of Col-0 and heterozygous *msp1-1* plants, were sown on petri dishes containing MS-Agar supplemented with 1% sucrose and vitamins (see Materials and methods) and grown in an *Arabidopsis* growth chamber under standard conditions. 6-week-old *msp1-1* (F) or Col-0 (G) plantlets grown the MS-plates (L6 growth stage), were transferred to liquid MS-media supplemented with 1% sucrose and vitamins (see Materials and Methods). These often develop short leaves with limited stems resulting with a bushy-like phenotype (H) that was previously noted by Dahan *et al*. (2014). (I) Five-month-old rosettes of homozygous *MSP1-1* plants grown on soil at 22^º^C under short-day conditions.

### MSP1 is located within the mitochondria, *in-vivo*

The N-terminal region of MSP1 shares homology with the consensus sequences of mitochondrial targeting signals (Fig. S2A, Table 1). This was also supported by global confocal microscopy-based GFP-fusions of *Arabidopsis* proteins (Lurin *et al*., 2004). To further confirm the intracellular location of MSP1, constructs encoding the plastidial Rubisco small subunit (RbcS) (Keren *et al*., 2009) and At.MSP1 were fused in-frame to GFP, introduced into *Arabidopsis* cells, via root transformation in tissue culture, and analyzed by confocal microscopy (Fig. 1B). As anticipated, the RbcS-GFP showed a typical root plastidial-like fluorescent patterns, whereas the signals of MSP1-GFP were detected as round or rod-shaped particles of about 1 to 2 μm in size, which overlapped with that of the MitoTracker marker, further demonstrating mitochondrial localization of At.MSP1.

### MSP1 roles are essential during early embryo-development in *Arabidopsis*

We sought to define the molecular functions of MSP1 by analyzing loss-of-function mutations within the *MSP1* locus. T-DNA insertional-lines inside *MSP1*’s gene-locus available at ‘The *Arabidopsis* Information Resource’ (TAIR), included SALK-018927 (T-DNA reported to be inserted in the 5’ UTR, 192 nucleotides before the start codon), SALK-142675 (*msp1-1*) (T-DNA insertion 312 nucleotides after the start codon), and SALK-070654 (T-DNA reported to be inserted in the 3’ UTR, 232 nucleotides after the termination codon) (Figs. 1A, S3A). Sequencing of genomic PCR products spanning the T-DNA insertional junctions confirmed the integrity of the *msp1-1* mutant line (Fig. S3B), but could not support insertional events within the UTRs in the SALK-018927 or SALK -070654 lines.

Similarly to various *emb* mutants (Meinke, 2020), the selfed progeny of the *msp1-1* showed a heterozygous to wild-type segregation ratio of ∼2:1 (*i*.*e*., 119 out of 175 seedlings). No homozygous seedlings for the *msp1-1* mutant-allele could be recovered among the seeds obtained from the heterozygous plants. Microscopic analyses showed that a quarter of the seeds analyzed in mature-green siliques of *msp1-1* (*i*.*e*., 25.1 ± 4.3%, n = 278) were small and deformed, and either collapsed during their development (Fig. 1C-1), or degenerated into shrunken seeds (Fig. 1C-2). A similar seed-morphology, known as *wrinkled1*, was previously associated with altered endosperm metabolism (Focks & Benning, 1998), but is also apparent in various mutants affected in mitochondrial biogenesis (Colas des Francs-Small & Small, 2014). Likewise, the siliques (12∼14 days post-fertilization, dpf) of heterozygous *msp1-1* contained ∼25% translucent seeds, which contained embryos arrested at the heart-to-torpedo transition stage, whereas their green seeds contained mature embryos (Fig. 1D). As the heterozygous *msp1-1* seedlings were phenotypically indistinguishable from the wild-type plants, we concluded that the insertional event in *MSP1* results in a recessive embryo-arrested phenotype, and that a single copy of the gene is sufficient to support ‘normal’ embryogenesis and seedling development.

### A modified method for embryo-rescue of *Arabidopsis msp1-1* mutant plants

As loss-of-function of *msp1-1* resulted in seed-abortion, we employed a modified embryo-rescue method that was previously used for the recovery of viable homozygous *nmat3* seedlings (Shevtsov-Tal *et al*., 2021). White seeds collected from surface-sterilized siliques (12∼14 dpf) of *msp1-1*, were germinated on MS-agar plates, supplemented with 1% sucrose and vitamins (Fig. 1E). Under these conditions we were able to obtain a few homozygous *msp1-1* plantlets that slowly developed (4-to-5 months) into a six-true-leaf (L6) growth-stage (Boyes *et al*., 2001). As the *nmat3* mutants, the homozygous *msp1-1* plantlets were able to develop beyond the L6-phase upon their transplant to a liquid growth medium (Fig. 1F). Wild-type embryos collected at the heart-to-torpedo stage (Col-0-heart) achieved a more robust growth than the *msp1-1* mutants (Fig. 1G). We also noticed a phenotypic variation, where some of the *msp1-1* mutants showed a bushy leaf morphology (Fig. 1H), which was also reported for other rescued seedlings, such as *emb* (Franzmann *et al*., 1989), *ndufv1* (Kuhn *et al*., 2015), *cod1* (Dahan *et al*., 2014) or *nmat3* mutants. A small number of rescued *msp1-1* plantlets, which were able to survive when transferred into a potting mix (Fig. 1I), displayed a curled-leaf phenotype (Katz *et al*., 2004) that was also noted in various *Arabidopsis* mitochondrial mutants (Colas des Francs-Small & Small, 2014). None of the rescued *msp1-1* seedlings was able to flower or to produce seeds. Thus, for the molecular characterization of *msp1*, we used 4∼5-month-old embryo-rescued MS-grown plantlets obtained from white seeds of heterozygous *msp1-1*plants. The seedlings were verified to be homozygous by sequencing of genomic PCR products.

### Rescuing *msp1-1* phenotypes by a constitutive expression of MSP1

As T-DNA lines often show multiple insertions (O’Malley *et al*., 2015; Pucker *et al*., 2021), and since we had only a single mutant line, it was important to confirm that the phenotypes associated with *msp1-1* indeed relate to the loss of MSP1. To facilitate a functional complementation, we cloned the *At*.*MSP1* gene under the control of 35S-promoter, transformed the construct into heterozygous *msp1-1* mutants and screened for homozygous *msp1-1* plants that harbor the *35S-MSP1* construct. Several plant-lines recovered among the progeny of the T2 plants that contained the *35S:MSP1* transgene, were also homozygous plants for the *msp1-1* insertion (Fig. S3C, *msp1*:*35S-MSP1*). One of the lines, plant-18 (Fig. S3C), was included as a control in the subsequent analyses.

### *msp1* mutants show altered *nad1* RNA processing in *Arabidopsis* mitochondria

The mitogenome of *Arabidopsis* harbors 57 genes encoding for different rRNAs, tRNAs and organellar proteins (Unseld *et al*., 1997; Sloan *et al*., 2018). We anticipated that MSP1 would function in mtRNA-metabolism. At first, the mtRNA profiles of embryo-rescued *msp1-1*, Col-0-heart, heterozygous *msp1-1* complemented *msp1*:*35S-MSP1* and 3-week-old MS-grown Col-0 plants (used as control) were assessed by RT-qPCR with oligonucleotides designed to the different exon-exon and exon-intron regions of each organellar transcript (Table S1). A strong reduction (∼10,000-fold) in the steady-state level of transcripts corresponding to a spliced *nad1* exons a-b product was noted in *msp1-1*, as compared to the control 3-week-old Col-0 plants (Fig. 2).

**Figure 2.**
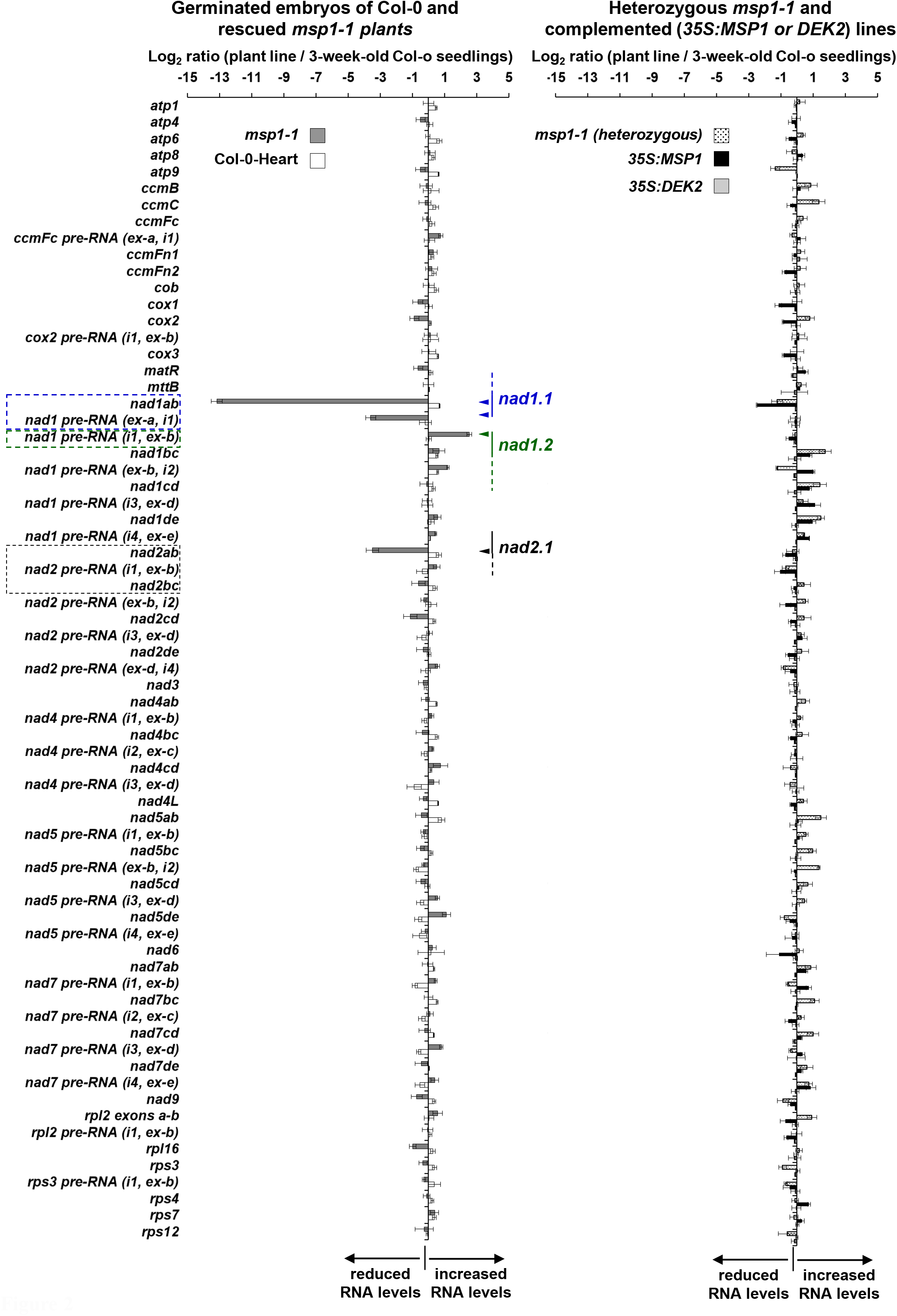
Transcript abundance in *msp1* mutants. RT-qPCR analyses of mitochondrial protein-coding genes was performed essentially as described previously (Zmudjak *et al*., 2013; Cohen *et al*., 2014; Sultan *et al*., 2016; Shevtsov-Tal *et al*., 2021). RNA extracted from 3-week-old seedlings of wild-type plants, 5-month-old rescued *msp1-1* mutant plantlets and plantlets obtained from the immature seeds of Col-0 plants containing embryos at the heart stage (Fig. 1D) was reverse-transcribed and the relative steady-state levels of cDNAs corresponding to the different organellar transcripts were evaluated by qPCR with primers which specifically amplified mRNAs (Table S1). Histograms showing the relative mRNAs levels (log2 ratios) in (1) rescued homozygous *msp1-1* and germinated embryos of wild-type (Col-0-heart) plantlets, or (2) heterozygous *msp1-1* (*htz*) plants and *msp1:35S-MSP1* (line #18, Fig. S3) complemented plants, versus those of 3-week-old MS-grown wild-type (Col-0) plants. Transcripts analyzed in these assays include the NADH dehydrogenase (CI) subunits *nad1* exons a-b, b-c, c-d, d-e, *nad2* exons a-b, b-c, c-d, d-e, *nad4* exons a-b, b-c, c-d, *nad5* exons a-b, b-c, c-d, d-e, *nad7* exons a-b, b-c, c-d, d-e, the cytochrome oxidase (CIV) *cox2* exons a-b, *ccmFc* exons a-b, and the ribosomal subunits *rpl2* exons a-b, *rps3* exons a-b, as well as the intronless transcripts of *nad3, nad4L, nad6, nad9* (lacking introns), the CIII *cob* subunit, CIV *cox1* and *cox3* subunits, the ATP synthase (CV) subunits *atp1, atp4, atp6, atp8* and *atp9*, genes encoding the cytochrome c maturation (*ccm*) factors *ccmB, ccmC, ccmFn1, ccmFn2*, the ribosomal factors *rps4, rps7, rps12, rpl16, rrn26*, and the *mttB* gene. Arrows indicate genes that are interrupted by group II intron sequences in *Arabidopsis* mitochondria (Sloan *et al*., 2018), while asterisks indicate transcripts where the mRNA levels were reduced in *msp1-1* plants, but not of the rescued Col-0 seedlings. The values are means of 15 reactions corresponding to five biological replicates (error bars indicate one standard deviation).

The mitochondrial *nad1, nad2, nad4, nad5, nad7, cox2, ccmFc, rpl2* and *rps3* genes are interrupted by group II introns that are removed post-transcriptionally to generate functional/translatable mRNAs. The processing of the *nad1, nad2* and *nad5* pre-mRNAs further involves the joining of several exons found in individually expressed gene-fragments, which are physically separated in the mitogenome by *trans*-splicing (Bonen, 2008). The ‘breaks’ between these gene-fragments are all within the 4^th^ domain (D4) of the *trans*-spliced introns. The maturation of *nad1* involves the processing of three RNA-fragments, *nad1*.*1, nad1*.*2* and *nad1*.*3* (Fig. S4A). Two of the *nad1* introns, including the 2^nd^ intron in *nad1*.*2* (*nad1i477*) and the 4^th^ intron (*nad1i728*) in *nad1*.*3* are postulated to undergo *cis*-splicing prior to the three RNAs being joined together via the *trans*-splicing of introns 1 (*nad1i394*) and 3 (*nad1i669*) (Bonen, 2008; Brown *et al*., 2014). Likewise, *nad2* maturation involves the joining of two pre-mRNA fragments separated by intron 2, while *nad5* is fragmented into three gene-pieces within its 2^nd^ and 3^rd^ introns (Bonen, 2008).

Altered splicing is assumed to be present in cases where the accumulation of a specific pre-RNA transcript is correlated with reduced levels of its corresponding mRNA. We considered that the large reduction we observed in transcripts corresponding to spliced *nad1* exons a-b relate to a defect in the excision of the first intron. Yet, the RT-qPCR data indicated that the *nad1*.*1* pre-RNA (amplified by primers designed to exon ‘a’ and *nad1* 11) was also notably reduced (about 750-fold) in *msp1-1* mutant. These data indicate to reduced stability of *nad1*.*1* rather than to altered *nad1* i1 splicing. The moderately increased level (*i*.*e*., 5.6 ± 1.2-fold) of *nad1*.*2* pre-RNA in *msp1-1* (amplified by primers designed to the intron’s D5 and exon-b) (Fig. 2) is thus anticipated to result as a consequence of the absence of the corresponding *nad1*.*1* RNA-fragment.

A milder effect was also apparent for *nad2* exons a-b splicing in *msp1-1* (*nad2ab*, 11.2 ± 1.4-fold reduction), while the relative level of the *nad2* i1 pre-RNA was slightly increased (1.5 ± 1.4-fold) in the mutant (Fig. 2). However, we noticed that the splicing efficiencies of *nad2* i1 are also reduced in other *Arabidopsis* mutants affected in the mtRNA-metabolism, some of which showing altered *nad1* processing (Table S2). Although various P-class PPRs have been previously shown to affect the splicing of organellar introns (Table S2), our data could not support a direct role for MSP1 in the excision of either *nad1* i1 or *nad2* i1. Other than the effects seen in the processing of *nad1* or *nad2*, the expression of different mtRNAs were seemingly not affected by the loss of MSP1. These included the intron-containing transcripts of *ccmFc, cox2, nad4, nad5, nad7, rpl2* and *rps3*, as well as the ‘intronless’ transcripts of *cox1* and *cox3* of CIV, ATP-synthase (CV) subunits, CCM-factors and ribosomal-related genes (Fig. 2). The accumulation of mtRNAs in heterozygous *msp1-1* were generally equivalent to those of the Col-0 plants, while the steady-state levels of many mtRNAs in germinated Col-0-heart embryos were slightly increased (2∼4-fold) in comparison to the wild-type plants (Fig. 2). Based on these data, we concluded that the *nad1* processing defects were due to the *msp1* mutation and not the embryo-rescue procedure. Furthermore, the RNA defects seen in *msp1-1* were mostly restored in the ‘complemented’ line, which express an intact *MSP1* gene (Fig. 2).

### *msp1-1* show reduced stability of the *nad1*.*1* pre-RNA fragment

We expect that reduced levels of the pre-*nad1*.*1* transcript and its corresponding spliced *nad1ab* product, rely on altered processing of *nad1*.*1* pre-RNA, which was not directly assayed in the RT-PCR experiment. This purpose, the mtRNA landscapes in Col-0 and *msp1-1* plants were further examined by RNA-seq analyses, using total seedling RNAs. The transcriptomic data showed that the relative levels of reads corresponding to the *nad1*.*1* gene-fragment (nucleotides 81,785– 83,966), were reduced to residual levels in *msp1-1* (Figs. 3, S4). Analysis of the relative read coverage of each individual exon, between wild-type (Col-0) and *msp1-1* plants, indicated a large reduction in *nad1* exon-a (Fig. 3A). Likewise, the ratio of read coverage between Col-0 and *msp1-1* across the whole *Arabidopsis* mitogenome indicated a single major peak corresponding to the *nad1* exon-a sequence (Fig. 3B). In agreement with the RT-qPCR data, no significant variations in the maturation of *nad1* exons b-c, c-d, or d-e transcripts were evident in the RNA-seq data (Figs. 3, S4). A minor increase in nucleotide abundances in *msp1-1* was apparent for *nad2* i1 (Data S1), whereas the levels and read-depth profiles of *nad2-a, nad2-b*, or other mtRNA transcripts in *msp1-1* were mostly equivalent to those of the wild-type plants (Figs. 3, S4, Data S1). Taken together, these data showed that MSP1 has a key role in the stabilization of *nad1*.*1* pre-mRNA, and that the loss of MSP1 reduces the excision efficiency of *nad2* i1, an effect which may relate to altered mitochondria biogenesis and/or gene-expression (Table S2).

**Figure 3.**
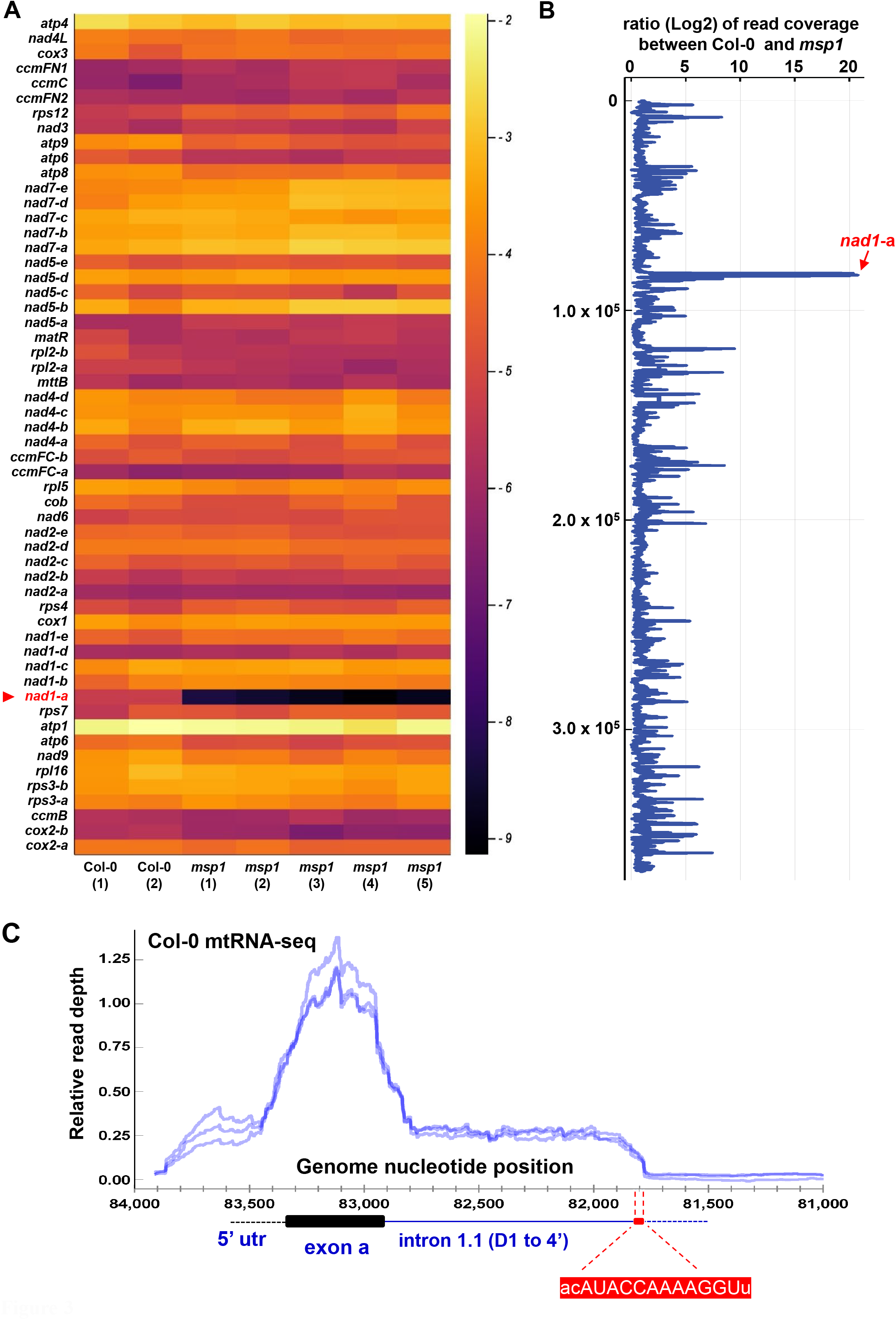
Transcription profiles of *msp1-1* mutant-line by RNA-seq analyses. Total RNA extracted from Col-0 and *msp1-1* plantlets was used to generate Illumina-sequencing (NovaSeq 6000, 150 bp paired-end) libraries (BioProjects PRJNA704631 and PRJNA768306). (A) Heatmap illustrating the relative expression levels between two wild-type (Col-0) plants and five mutant (*msp1*) lines of each individual exon. Darker colors indicate to reduced exon levels in *msp1 vs* the Col-0 plants. (B) Ratio of read coverage between Col-0 and *msp1* across the whole genome. (C) RNA-seq data of *Arabidopsis* mitochondrial *nad1*.*1* transcript (BioProject PRJNA768306, corresponding to 3 individual Col-0 plants), showing the read depth along the *nad1*.*1* sequence (containing exon ‘a’ and domains 1-4 of *nad1* intron 1.1). The predicted MSP1 RNA-binding site is indicated below the plot.

### MSP1 is expected to be associated with an RNA motif found in its defined intron target

A combinatorial code for RNA-recognition by PPR proteins has been worked out, where amino acids at positions 5 and 35 of each domain act as specificity-determining residues (Barkan *et al*., 2012; Cheng *et al*., 2016). Several PPRs were previously shown to affect the stability of their RNA-ligands by protecting them from excessive exonuclease activities (Barkan & Small, 2014). *In-silico* data (see Fig. S5A) indicate the preferred sequence 5’-UACCAAAAGGU for MSP1’s PPR 4-11 motifs, while PPRmatcher (Royan *et al*., 2021) was used to score potential alignments to every possible position (735,616x) in the mtDNA (Fig. S5B). Comparative analyses with the updated *Arabidopsis* mitogenome (Sloan *et al*., 2018), revealed three mtDNA loci exactly matching this sequence. One of these is found within the *nad1* i1, 1,140 bp downstream to exon-a (nucleotides 81,785-81,796) (Figs. 3A, S4A, S5), whereas the two other sites are mapped to noncoding/unexpressed loci. The RNA-seq data of Col-0 plants showed that the relative read-depth in the *nad1*.*1* region is noticeably reduced immediately following the predicted MSP1 binding-site (Fig. 3C). These data strongly support the association of MSP1 with a specific region in D4 of *nad1* i1, which may be required for the correct 3’-end processing of *nad1*.*1* transcript.

### MSP1 binds to *nad1*.*1* by associating with nucleotides at the predicted binding site and protects *nad1* i1 from excessive 3’-to-5’ exonucleolytic degradation, *in-vitro*

The binding activity and specificity of MSP1 to *nad1*.*1* RNA was analyzed by biochemical analyses, in-vitro. As attempts to express and purify an intact MSP1 were unsuccessful, we analyzed the binding properties of a recombinant product, which harbors the PPR motifs 4-14 of MSP1. The codon optimized rMSP1 construct was cloned in-frame to GST in its N-terminus and a C-terminal 6xHis tag, expressed in *E. coli*, purified by affinity chromatography (Fig. S6), and studied for its binding characteristics by filter-binding assays (Keren *et al*., 2008). The rMSP1 protein was found to binds to *nad1* i1 (nucleotides 80,646-81,796) with a micromolar affinity (calculated *K*_*d*_ = 4.01 ± 0.31 μM) and binding stoichiometry (n) of 1.0 ± 0.05 (Fig. 4A). We further analyzed the association between rMSP1 and a *nad1* i1 RNA-ligand (i.e., *nad1*.*1*-ΔMBS) that is lacking the postulated MSP1 binding site (MBS, UACCAAAAGGU). The binding activity of rMSP1 to *nad1*.*1*-ΔMBS was found to be notably lower (*K*_*d*_ = 37.06 ± 7.93 μM) than that of the ‘intact’ *nad1* i1. For binding competition experiments, a synthetic UACCAAAAGGU RNA-oligo was used in the binding reactions in a 100:1 molar ratio to *nad1* i1. In support of the specific association of MSP1 with this region, the addition of the competitor RNA reduced the maximal amount of bound *nad1* i1 by about 3 orders of magnitude (*K*_*d*_ ≈ 13 mM), whereas the binding of MSP1 to *nad1* i1 was seemingly unaffected by a synthetic poly-C RNA-oligo (Fig. 4B, *K*_*d*_ = 3.77 ± 0.39 μM). No binding activity was indicated for GST alone, which was used as a control for the integrity of the binding assays.

**Figure 4.**
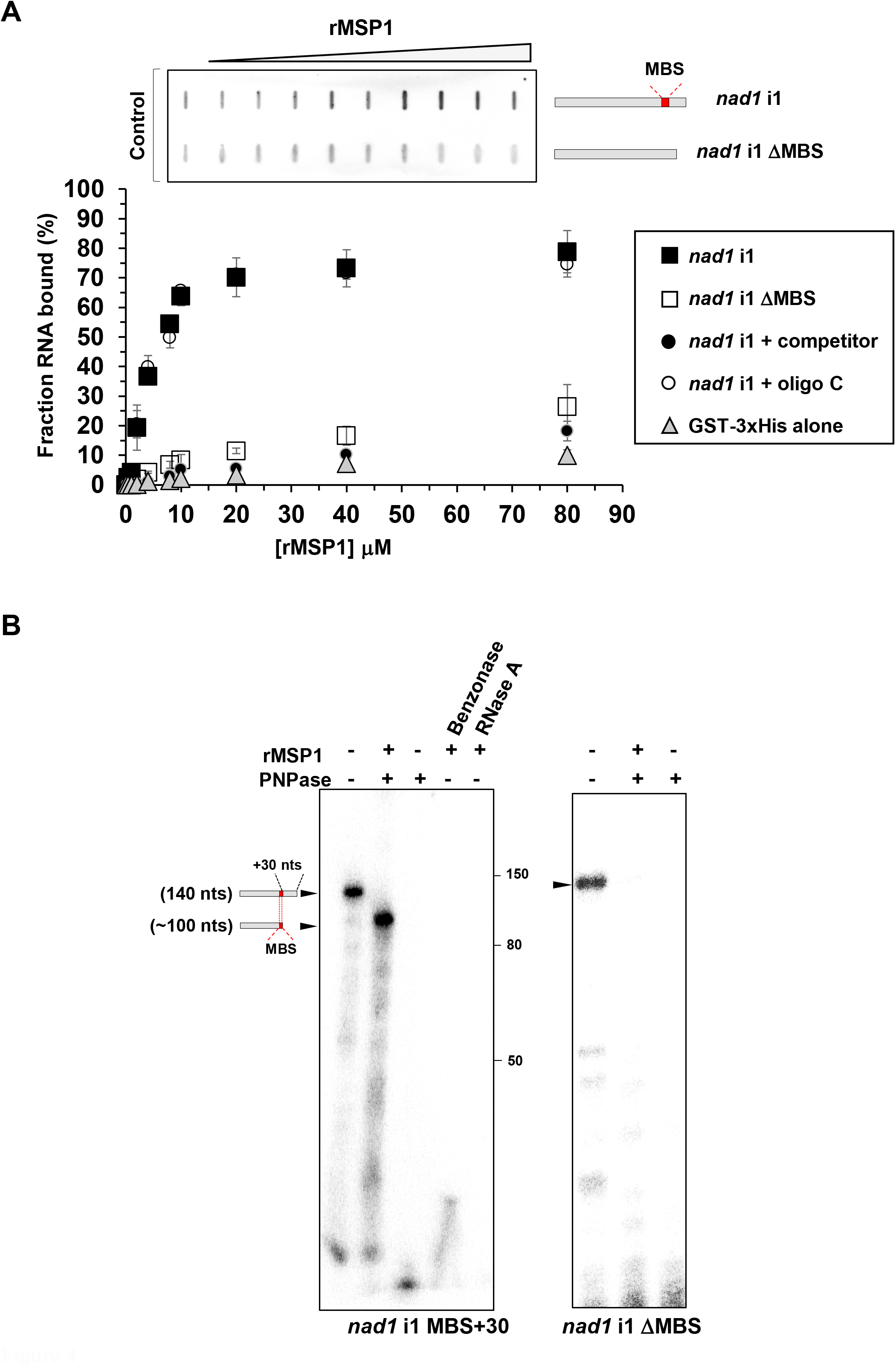
MSP1 is associated with a specific intronic region in *nad1* i1 and eliminates 3′-end exoribonucleolytic degradation by PNPase, in-vitro. (A) The association between the recombinant GST-rMSP1-6xHis (rMSP1, Fig. S6) protein and *nad1* pre-RNA transcripts was analyzed by incubating increasing amounts of the protein (0 – 80 μM) with in-vitro transcribed labeled RNAs. Binding was analyzed by filtration of the RNA-protein complex filter binding assays (Ostersetzer *et al*., 2005; Barkan *et al*., 2007; Keren *et al*., 2008). A raw slot-blot data is shown from representative experiments on top of the graph. The graph shows the binding activity of MSP1 to *nad1* i1in the presence or absence of competitor RNA-oligos, or to *nad1* i1 ΔMBS. GST-6xHis was used as a control for the *nad1* i1 binding assays. The values are means of four independent measurements (error bars indicate one standard deviation). (B) MSP1-mediated protection of *nad1* i1 from 3′→5′ exonuclease activity. Polynucleotide phosphorylase (PNPase)-protection assays (Prikryl *et al*., 2011) were performed with 5′-end ^32^P-labeled RNAs corresponding to *nad1* i1 or *nad1* i1 ΔMBS. The RNA templates were denatured (i.e., 2 min at 95 °C, and snap cooled on ice) and then incubated in the presence or absence with rMSP1 prior to their use in the PNPase-reaction assays. The *nad1* i1 stabilization effects of rMSP1 were further analyzed in the presence of the exonucleases Benzonase (cleaves all forms of DNA or RNA into 3 to 5 bases in length) or RNase A (cleaves the phosphodiester bond between the 5’-ribose of a nucleotide and the phosphate group attached to the 3’-ribose of an adjacent pyrimidine nucleotide). The reactions were stopped by heating the samples in RNA sample buffer containing 60% formamide, and the RNAs were then resolved on 10% AA gels containing 7M urea in 1×TBE buffer. The position of protected *nad1* i1 terminus was determined from the size of the corresponding end-labeled RNA.

Here, we further aimed to determine whether the binding of MSP1 to the UACCAAAAGG site may relate to the altered *nad1*.*1* stability seen in *msp1-1* mutant. PNPase-protection assays with 5’-labeled *nad1* i1 RNA-ligands, further indicated a role for MSP1 as a mitochondrial *nad1*.*1* stability factor. The addition of rMSP1 to the reaction assays led to the accumulation of a specific RNA product of ∼100 nts, generated by the 3’-to-5’ PNPase exonuclease (Figs. 3A, S4A), which is expected to corresponds to the mature *nad1*.*1* transcript seen in the RNA-seq data. In the absence of rMSP1, the *nad1*.*1* RNA ligand was fully digested by the PNPase (Fig. 4B). Likewise, the rMSP1 was not able to protect the *nad1*.*1*-ΔMBS transcript, which is lacking the UACCAAAAGGU site, from exonucleolytic processing by PNPase (Fig. 4B). rMSP1 also failed to stabilizes *nad1* i1 from endonucleolytic cleavage by the Benzonase enzyme. The canonical RNAse A enzyme worked like Benzonase, and also led to the complete degradation of *nad1* i1 in the presence or absence of rMSP1 (Fig. 4B).

### MSP1 functions are required for the biogenesis of respiratory CI in *Arabidopsis* plants

Nad1 is essential for CI biogenesis and function (Braun *et al*., 2014; Guerrero-Castillo *et al*., 2017; Ligas *et al*., 2019). The RNA maturation defects we see (Figs. 2-3, S4-S5) suggest that Nad1 exists in extremely low levels (or absent) in the mutant-line. Altered *nad2* maturation is also anticipated to affect the biogenesis of the respiratory system, as Nad2 is also postulated to be incorporated during initial stages in CI assembly in Arabidopsis plants (Ligas *et al*., 2019). The relative accumulation of different organellar proteins in *msp1-1* versus Col-0 plants was estimated by immunoblotting (Table S3). These analyses indicated that Nad1 (∼36 kDa) (Shevtsov-Tal *et al*., 2021) is reduced to trace levels in *msp1-1*, but is clearly visible in the heterozygous, Col-0-heart or ‘complemented’ lines (Fig. 5A, Table S4A). The appearance of various bands in the Nad1-blot of *msp1-1* may correspond to aberrant Nad1 products that can be translated by alternative AUG sites present in semi-processed *nad1* transcripts (Figs. 2-3), which are recognized by the polyclonal anti-Nad1 antibodies. The steady-state levels of CA2 (30 kDa) and Nad9 (23 kDa) subunits of the ‘membranous’ and ‘soluble’ domains, respectively, of CI were found to be slightly higher in *msp1-1* than in Col-0 plants (*i*.*e*., 3.7 ± 0.5 and 2.1 ± 0.3, respectively). The level of Atp2 (CV) in *msp1-1* was equivalent to that of the Col-0 plants, whereas the signals of the nuclearly-encoded Atp1 (CV), RISP (CIII), VDAC, or the organellar-encoded COX2 (CIV), were found to be higher in the mutant or Col-0-heart seedlings (Fig. S7A, Table S4A). The accumulation of different organellar proteins in heterozygous *msp1-1* was generally equivalent to Col-0 plants.

**Figure 5.**
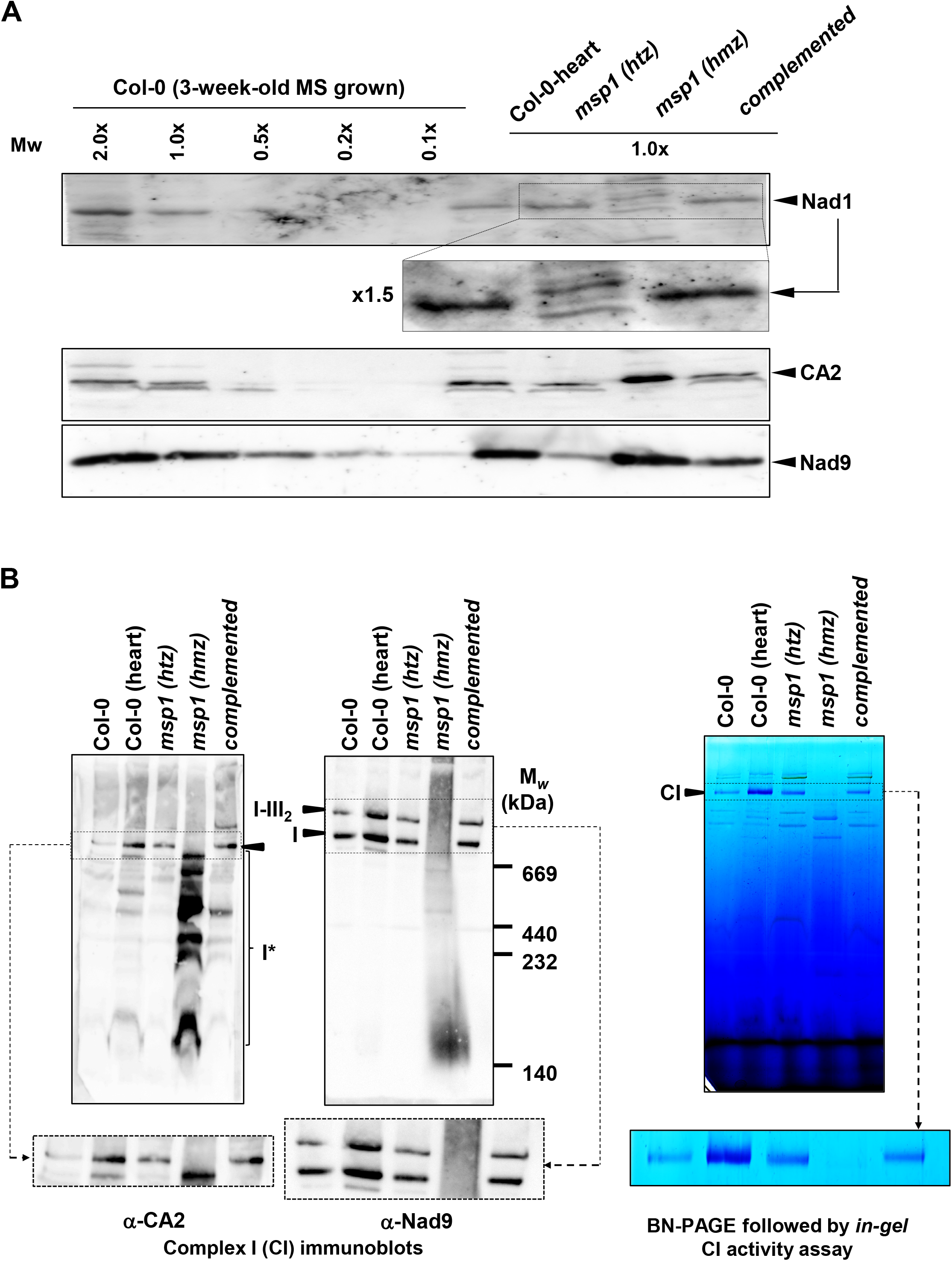
Analyses of the organellar protein profiles of wild-type and *msp1-1* plants. (A) Immunoblots with proteins (about 50 μg) of mitochondrial enriched membranous fractions (Pineau *et al*., 2008; Sultan *et al*., 2016), isolated from MS-grown Col-0 and rescued *msp1-1* plants. The blots were probed with polyclonal antibodies (Table S4) raised against the NADH-oxidoreductase subunit 1 (Nad1), γ-carbonic anhydrase-like subunit 2 (CA2) and Nad9 proteins. Detection was carried out by chemiluminescence assays after incubation with HRP-conjugated secondary antibody (note that the dashed box in the Nad1 blot corresponds to a 1.5x magnified image copied below the blot). (B) Blue native (BN) PAGE analyses of crude organellar membranous fractions. An aliquot equivalent to 40 mg of crude *Arabidopsis* mitochondria extracts, obtained from wild-type plants, *msp1* mutant-lines and complemented *msp1:35S-MSP1*, was solubilized with 5% (w/v) digitonin, and the organellar complexes were resolved by BN-PAGE. For immunodetection, the proteins were transferred from the native gels onto a PVDF membrane. The membranes probed with antibodies raised to complex I (CI) NADH-oxidoreductase subunit 9 (Nad9) and γ-carbonic anhydrase-like subunit 2 (CA2) proteins. In-gel CI activity assays were performed essentially as described previously (Eubel *et al*., 2005). Arrows indicate to the native supercomplex I+III2 (∼1,500 kDa) or holo-CI (∼1,000 kDa), while I* indicates the presence of CI assembly intermediates.

The respiratory system in plants also possesses type-II NAD(P)H dehydrogenases (NDs) and alternative oxidases (AOXs) (Popov *et al*., 2020; Møller *et al*., 2021). In *Arabidopsis*, mutants affected in their respiration typically show notable increases in the expression or accumulation of AOXs and NDs. Likewise, the immunoblots indicated that AOX1/2 signal was higher in the *msp1-1* mutant (Fig. S7A, Table S4A). An upregulation in the accumulation of several AOXs and NDs was also evident by RT-qPCR, where transcripts corresponding to AOX2, NDA2, AOX1A or NDB2 were increased in the mutant by 3.2-, 3.4-, 16- and 10-fold, respectively (Fig. S7B).

Considering the effects of MSP1 on Nad1 expression, we further analyzed the relative abundances of native respiratory complexes I, III, IV and V in Col-0 and mutant plants by Blue-native (BN) PAGE analyses (Figs. 5B, S7C). Arrows indicate native complexes I (∼1,000 kDa), CIII dimer (III_2_, ∼500 kDa), supercomplex I+III_2_ (∼1,500 kDa) (Braun *et al*., 2014; Senkler *et al*., 2017), CIV (∼220 kDa), and CV (∼700 kDa). The BN-PAGE/immunoblot analyses showed that CI is below detectable levels in *msp1-1*, but is correctly assembled in the complemented line (Fig. 5B, Table S4B). A significant reduction in CI was also evident by ‘*in-gel*’ CI-activity assays, which further indicated that none of the sub-CI particles in *msp1-1* were active (Fig. 5B). No significant changes in CI accumulation were seen in the heterozygous line. The immunoblots of CA2 also indicated the presence of CI assembly intermediates (Ligas *et al*., 2019) of about 200-to-700 kDa in size in *msp1-1* (Fig. 5B-I*). BN-PAGE analyses of *nmat1*, which is impaired in the splicing of *nad1* i1 (Keren *et al*., 2012), also indicated the presence of several sub-CI particles, with a dominant band of ∼700 kDa (Fig. S8). Accumulation of partially-assembled CI intermediates was also evident in the germinated embryos of Col-0 (Col-0-heart), but these plants rather exhibited increased CI level and activity (Fig. 5B, Table S4B). Due to the cross-reactivity of the anti-Nad1 antibodies with various organellar proteins (Fig. 5B) (Shevtsov-Tal *et al*., 2021), it was impossible to determine whether the CI intermediates harbor the Nad1 protein.

The BN-PAGEs further showed that CIII, CIV and CV are more abundant in *msp1-1* (possibly to accommodate changes in electron transport and/or H^+^-pumping rates following the loss of holo-CI), while their levels in the complemented line were generally equivalent to those of the Col-0 seedlings (Fig. S7C, Table S4B). The loss of CI in the *msp1-1* mutant-line is further correlated with the upregulation of mRNAs and proteins associated with the alternative pathway of the electron transport (*i*.*e*., NDs and AOXs) (Figs. S7A, S7B).

## Discussion

### *MSP1* encodes a PPR factor that is pivotal for *nad1* pre-RNAs maturation

Embryogenesis and seed-germination are complex developmental processes that rely on increased cellular metabolism to support their high-energy demands, see e.g., (Best, C. *et al*., 2020). These correlate with experimental data that show that the expression/processing of mtRNAs in plants is both developmentally and environmentally responsive (Li-Pook-Than *et al*., 2004; Law *et al*., 2012; Dalby & Bonen, 2013). Likewise, the expression of different factors that function in mtRNA-metabolism, including MSP1 (Table 1, Fig. S1), is tightly regulated during early plant development or under stress conditions (Law *et al*., 2012; Chen *et al*., 2018). To gain more insights into MSP1-mediated roles, we analyzed an embryo-rescued (Shevtsov-Tal *et al*., 2021) *msp1-1* knockout-line and a complemented plant expressing the *MSP1* gene in the genetic background of homozygous *msp1-1* mutants (Figs. 1, S3, Table S5). Our data show that *msp1* mutants are strongly affected in the maturation of *nad1* exon-a (Figs. 2-3), whose processing involves the splicing of two mtRNA fragments, *nad1*.*1* and *nad1*.*2*, separated by intron 1 (Fig. S4A). The analysis of the RNA profiles of Col-0 and *msp1* plants indicated that MSP1 has a key role in *nad1*.*1* transcript maturation, by a molecular mechanism that is not directly related with the splicing of *nad1* i1 (i.e., splicing defects are typically associated with the accumulation of pre-RNAs and the reduced levels of their corresponding mRNAs).

Various PPRs have been implicated in the post-transcriptional generation of 5’ or 3’ termini of organellar transcripts in plants (Barkan & Small, 2014). In *Arabidopsis* mitochondria, these involve at least MTSF1, MTSF2, PPR19, RPF1, RPF2, RPF3, MTSF3 (Colas des Francs-Small & Small, 2014; Zmudjak & Ostersetzer-Biran, 2017; Wang *et al*., 2022). These may drive the accumulation of short RNA footprints (cosRNAs), often corresponding to the precise termini of organellar transcripts (Pfalz *et al*., 2009; Ruwe & Schmitz-Linneweber, 2012). Likewise, the MSP1 binding site is located at the very end of the mature *nad1*.*1* transcript (Figs. 3C, S4, S5), although was not previously identified among the mitochondrial cosRNAs (these are seemingly less obvious than the plastidial ones) (Ruwe *et al*., 2016). The RNA-seq data (Figs. 4, S4, S5) further showed that *nad1*.*1* is reduced to residual levels in the *msp1* mutant, indicating that MSP1 is pivotal for its accumulation. Based on these data we speculated that through its association with a specific site in *nad1*.*1* (Fig. 4, S4A, S5), MSP1 facilitates the generation of its 3’-terminus and stabilizes it -a prerequisite for the *trans*-splicing of *nad1* i1. In-vitro biochemical assays provided with further evidence that the association of MSP1 with *nad1*.*1* rely on the UACCAAAAGGU site (Fig. 4A), and that this interaction is sufficient to block 3′→5′ degradation by the PNPase beyond this site, but not to protect the RNA from endonucleolytic digestion (Fig. 4B).

### The excision of *nad2* i1 in angiosperms: an epistatic regulation associated with mtRNA-metabolism or the biogenesis status of the OXPHOS system?

A specific role for MSP1 in the excision of *nad2* i1 remains ambiguous. A region with a limited homology to the predicted MSP1 binding site can be found in *nad2* i1 (CAAAAGGU, nts 162,555-162,562). It is possible, therefore, that MSP1 may enhance the excision efficiency of *nad2* exons a-b, *e*.*g*., by assisting in the folding of this highly-derived intron-RNA (Bonen, 2008). However, reduced *nad2* i1 splicing is also evident in many other mutants affected in mtRNA-metabolism (Table S2). It’s possible that the processing of various mtRNAs (e.g., *nad1, nad5*, or *nad7*) is linked to the splicing of *nad2* i1 through an as yet unknown mechanism. Alternatively, The processing of *nad2* may involve an epistatic effect (e.g., the translation of some mRNAs in algae chloroplasts is orchestrated with the *de-novo* assembly of their related complexes) (Choquet & Wollman, 2009). However, the excision of *nad2* i1 was not affected in *ca2* (Fig. S8) (Cordoba *et al*., 2016), nor in a mutant lacking another CI subunit (Meyer *et al*., 2009), which are both strongly affected in CI assembly. Based on these data, it is unlikely that CI-biogenesis is directly related to *nad2* pre-RNA processing. Other than the processing defects seen in *nad1* and *nad2*, the expression of other organellar transcripts remained mostly intact in *msp1-1* (Figs. 2-3, Data S1).

### The evolution of a *nad1ab trans*-splicing event in the mitochondria of spermatophytes

The regions corresponding to the first two exons in *nad1* in the mitogenomes of algae or basal land plants, such as liverworts (as their bacterial *NuoH-*related genes), are found as uninterrupted ‘intronless’ reading-frames. The *nad1* i1 (*nad1i394*) sequence is also not present in the mtDNAs of mosses, or hornworts, which harbor another intron (*nad1i287*) in a different mtDNA-locus (Fig. 6). Within angiosperms (besides *Viscum* sp. that lacks the *nad* genes entirely, (Petersen *et al*., 2015; Skippington *et al*., 2015), contain a *trans*-spliced *nad1* i1 where the putative MSP1 binding site seems perfectly conserved (Figs. 6, S9). Comparative analyses of genomic databases indicated that genes related to *MSP1* are present in both monocot and dicot species (Figs. 6, S10A). Particularly, the homologous gene in maize (*DEK2*) (Fig. S10B) was shown to affect the processing of *nad1* (Qi *et al*., 2017). Based on RT-PCRs, the authors considered a role for DEK2 in the splicing of *nad1* i1 (Qi *et al*., 2017). Yet, the accumulation of *nad1*.*1* was not assessed, and thus it is possible, even likely, that DEK2 has a similar role to that of MSP1, *i*.*e*., the stabilization of *nad1*.*1* transcript. The predicted RNA-binding site for DEK2 is identical to that of MSP1 (Fig. S9), and is present in the equivalent position at the end of the *nad1*.*1* (nts. 120612-120603) in the mtDNA of maize (NC_007982). We further noted that the transformation of the *Zm*.*DEK2* gene into Arabidopsis, complements the *msp1* mutation and restores the maturation of *nad1ab* (Fig. 2), demonstrating that *DEK2* encodes an orthologue of Arabidopsis.

**Figure 6.**
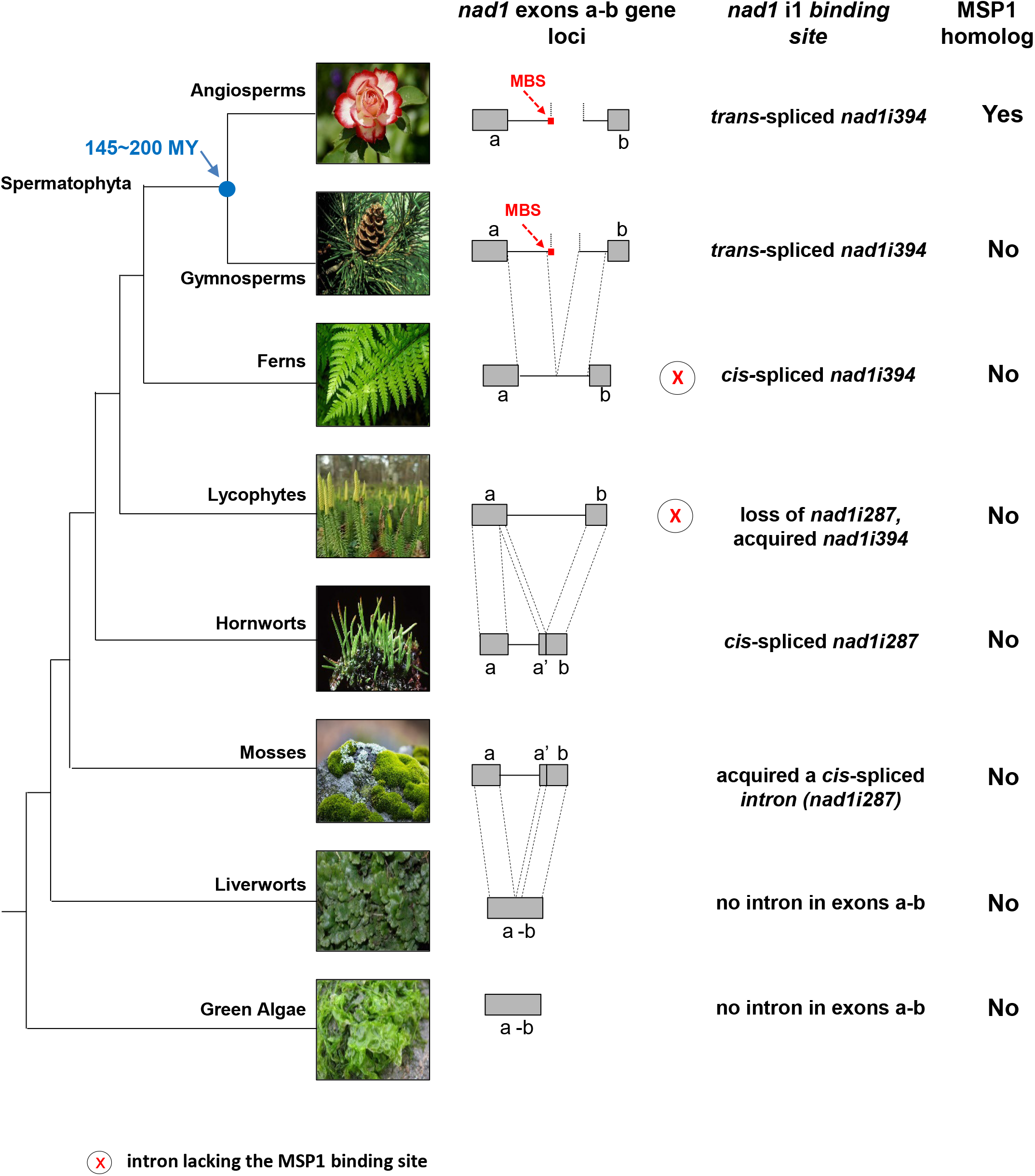
phylogenetic distribution of MSP1 gene and its genetically-defined *nad1* intron 1 RNA target. Genomic and organellar sequences available at the NCBI were used to search for *MSP1*-related PPR genes and *nad1* sequences in different plant clades, using The Basic Local Alignment Search Tool (BLAST). The results of the BLAST searches were integrated into the phylogenetic relationships among photosynthetic organisms, as indicated for the terrestrial flora (Kenrick *et al*., 2012; Delwiche & Cooper, 2015). Branch lengths as illustrated do not represent phylogenetic distances and are intended to indicate instead relationships among different clades. Mitochondrial *nad1* intron 1 (*nad1* i1) and the nuclear *MSP1* genes are shown as circles and boxes, respectively. Black boxes (MSP1) and black circles (*nad1* i1 in a *trans-* or *cis-*configurations) show that corresponding sequences are present, whereas open boxes indicate that the gene is absent in the organisms belonging to a specific plant clade. Pattern fill indicates the presence of homologous sequences, whereas red crosses indicate the absence of the proposed MSP1 binding site.

We anticipated that gymnosperms also encode homologs of the MSP1/DEK2 proteins. Yet, no potential orthologs for these factors could be supported in gymnosperms, although the intronic binding site is perfectly conserved in these plants (Fig. S9). It is therefore expected that this site has already been under selection in early spermatophytes, apparently as a target for a different (PPR) factor. Still, an orthologues MSP1 factor might be identified in some gymnosperms, once full genome assemblies are established for different cycads, ginkgo, yews or conifers (Liu *et al*., 2021). Based on these observations, we speculate that MSP1 was recruited 150∼200 MYA (Kenrick *et al*., 2012; Delwiche & Cooper, 2015) to facilitate the generation of 3’-ends in fragmented *nad1* i1 sequences already present in the mtDNAs of early spermatophytes. The ability of different intron-RNA segments to splice in *trans* could be followed by genome fragmentation, *i*.*e*., recombination and insertion of individual transcriptional initiation and termination sites during evolution. In fact, *trans*-splicing can be facilitated in model group II introns that are artificially fragmented within D4 (Belhocine *et al*., 2008). This remarkable molecular mechanism seems related to the evolution of the spliceosomal system (Schmitz-Linneweber *et al*., 2015; Zimmerly & Semper, 2015).

Besides to its role in stabilizing/determining *nad1*.*1* 3’-end, MSP1 may function in other activities, such as the recruitment of splicing cofactors, or in the association of *nad1*.*1* with *nad1*.*2* to form a catalytically active intron-RNP complex. Yet, EMB2654 and PPR4, are obligatory required for the *trans*-splicing of *rps12* (Lee *et al*., 2019), and not for transcript accumulation, even though they bind near the ends of their respective transcripts. These data indicate that RNA stabilization/trimming and splicing are not necessarily linked to one-another, although such effects cannot be ruled out yet for MSP1 in Arabidopsis mitochondria.

### Impaired CI biogenesis in *msp1* mutants and its effect on embryogenesis, seed-germination and plant development

The respiratory machinery is made of four major electron transport carriers, denoted as CI-to-CIV, and the ATP synthase enzyme (CV). The loss of MSP1 is detrimental during early embryogenesis, most likely as the Nad1 is essential for the biogenesis of an active holo-CI (Braun *et al*., 2014; Ostersetzer-Biran, 2016) (Fig. 5B, S7C). Likewise, the germination of *msp1-1* plantlets was enabled only under in-vitro conditions (Shevtsov-Tal *et al*., 2021) (Fig. 1C), which were able to continue their development beyond the L6-growth stage only when transferred to a liquid MS-medium. BN-PAGE analyses indicated that a holo-CI cannot be assembled in *msp1*, which seems to lack Nad1 (Fig. 5). The levels of the respiratory complexes CIII, CIV and CV were increased in the mutant (Fig. S7, Table S4), possibly to increase ATP-production, as well as for the reduced-NADH homeostasis whose accumulation is toxic to cells (Yang *et al*., 2020). The defects in CI are also associated with the upregulation of different NDs or AOXs (Fig. S4B), effects which were observed in many other plants affected in mitochondrial functions (Best, C. *et al*., 2020).

Nad1 is likely incorporated early during CI biogenesis, and was previously identified (mainly) in a ∼450 kDa sub-CI particle (Ligas *et al*., 2019). BN-PAGE indicated the presence of partially assembled CI intermediates in *msp1* (≥200∼700 kDa), with a major band of ∼700 kDa in *nmat1* (Figs. 5B, S7). The differences seen in patterns of sub-CI particles in different mutant-lines could correspond to altered levels of *nad1* mRNA between, e.g., *nmat1* (strongly reduced) (Keren *et al*., 2012) and at levels below a critical threshold to promote CI functions in *msp1* or *nmat3* mutants (Figs. 2-4, S4). Accordingly, the phenotypic variation between different Arabidopsis mutants could relate to residual differences in CI levels (Kuhn *et al*., 2015), or to developmental effects mediated by retrograde signaling (Woodson & Chory, 2008; Schwarzländer *et al*., 2012).

## Supporting information

Dataset S1 - RNA-seq data

## Acknowledgments

We thank Dr. Etienne Meyer for providing the anti-NAD1 antibodies, Prof. Hans-Peter Braun for providing us laboratory procedures for BN-PAGE analysis, Prof. Ariel Chipman for his help with Nomarski microscopy analyses, and Dr. Hakim Mireau for his help with MSP1 binding predictions and useful discussions regarding the role of MSP1 in mtRNA-metabolism. A special thank goes to Prof. Gadi Schuster (Technion) for providing the PNPase enzyme and useful discussions regarding the MS, and Dr. Michal Zmudjak, a former graduate student of O.O.B, for her inspiring role in establishing the modified embryo-rescue method. This work was supported by grants to O.O.B from the ‘Israeli Science Foundation’ ISF grants no. 1834/20 and 3254/20, and by grants to H.Z. from the National Natural Science Foundation of China (32061143022).

## Author contributions

<u>Dr. Corinne Best</u>: Plant growth and analysis, establishment of embryo rescued mutant lines, RNA and protein analysis. <u>Ron Mizrahi</u>: Expression and analysis of recombinant MSP1 protein and its RNA binding characteristics, help with GFP localization analysis. <u>Rana Edris</u>: Plant growth and analysis of *ca2* mutant lines. <u>Hui Tang</u>: Phylogenetic analysis of the MSP1 protein family. <u>Dr. Hagit Zer</u>: GFP localization analysis, cloning, assisted in the expression of recombinant proteins. <u>Dr. Catherine Colas des Francs-Small:</u>analysis of the RNA landscapes of wild type and mutant lines. <u>Dr. Omri M. Finkel</u>: assisted with Bioinformatic analyses of the RNA landscapes of wild-type plants and mutant lines. <u>Prof. Hongliang Zhu</u>: Bioinformatic analyses, phylogenetic map of the MSP1 protein family in plants. <u>Prof. Ian Small</u>: analysis of the RNA landscapes of wild type and mutant lines, prediction of MSP1 RNA binding site, and manuscript preparation. <u>Prof. Oren Ostersetzer-Biran</u>: PI, manuscript preparation and corresponding author.

## Data availability

The Arabidopsis Information Resource (TAIR) maintains a database of genetic and molecular biology data for the model higher plant *Arabidopsis thaliana*: http://www.arabidopsis.org. High-throughput RNA sequences are available as BioProjects PRJNA704631 and PRJNA768306. Genome Maps is an opensource collaborative initiative available in the GitHub repository (https://github.com/compbio-bigdata-viz/genome-maps). The ‘AlphaFold Protein Structure Database’ provides open access to protein structure predictions (https://alphafold.ebi.ac.uk).. Accession numbers: Arabidopsis mitogenome BK010421.1. MITOCHONDRIAL SPLICING/STABILITY PPR FACTOR 1 (MSP1) accession number: AT4G20090. RNA-seq data: BioProjects PRJNA704631 and PRJNA768306. A complete version of the Materials and Methods is available in the supplemental information.

## Supporting Information

**Figure S1**. *MSP1* gene expression patterns at different tissues and during various developmental stages.

**Figure S2**. MSP1 protein structure and its predicated RNA biding site.

**Figure S3**. PCR screening of *msp1* plants.

**Figure S4**. The mitochondrial transcription profiles of *msp1-1*.

**Figure S5**. Analysis of MSP1 RNA binding site.

**Figure S6**. Expression and purification of a recombinant rMSP1 protein.

**Figure S7**. The relative accumulation of organellar proteins and respiratory complexes in wild-type and *msp1-1* plants.

**Figure S8**. Relative accumulation and activity of respiratory CI in wild-type and *nmat1* plants.

**Figure S9**. Splicing efficiencies in *ca2* mutants.

**Figure S10**. The predicted MSP1 binding site is highly conserved among different spermatophytes.

**Figure S11**. Phylogenetic analysis of homologous MSP1 proteins in plants.

**Table S1**. Lists of oligonucleotides used for the analysis of the splicing profiles of wild-type and mutant plants by RT-qPCR experiments.

**Table S2**. List of splicing factors in *Arabidopsis* and their mtRNA profiles.

**Table S3**. List of antibodies used for the analysis of Col-0 and *msp1* mutants.

**Table S4**. Relative accumulation of mitochondrial proteins in wild-type and *msp1* plants.

**Table S5**. Oligonucleotides used in screening of individual T-DNA insertion lines in *Arabidopsis* and cloning of the *MSP1*-*GFP* gene-fusion construct.

**Data S1**. mtRNA landscapes of *msp1* mutants.

