## Supplementary material for "*MSP1* encodes an essential RNA-binding PPR factor required for *nad1* maturation and complex I biogenesis in *Arabidopsis* mitochondria": Dataset S1 - RNA-seq data

**Supplementary Data S1. mtRNA landscapes of *msp1* mutants.**

Total mtRNA extracted from Col-0 and embryo-rescued *msp1-1* plantlets was used to generate Illumina sequencing libraries (BioProjects PRJNA704631 and PRJNA768306). The alignments were visualized using the IGV genome browser (Thorvaldsdottir et al., 2013). Data are shown for the regions corresponding to different organellar transcripts in *Arabidopsis* mitochondria (Best et al., 2020). Green, blue and red lines point to variation in sequence between the mtRNA (RNA-seq data) and the updated mitogenome sequence of *Arabidopsis* (Sloan et al., 2018).

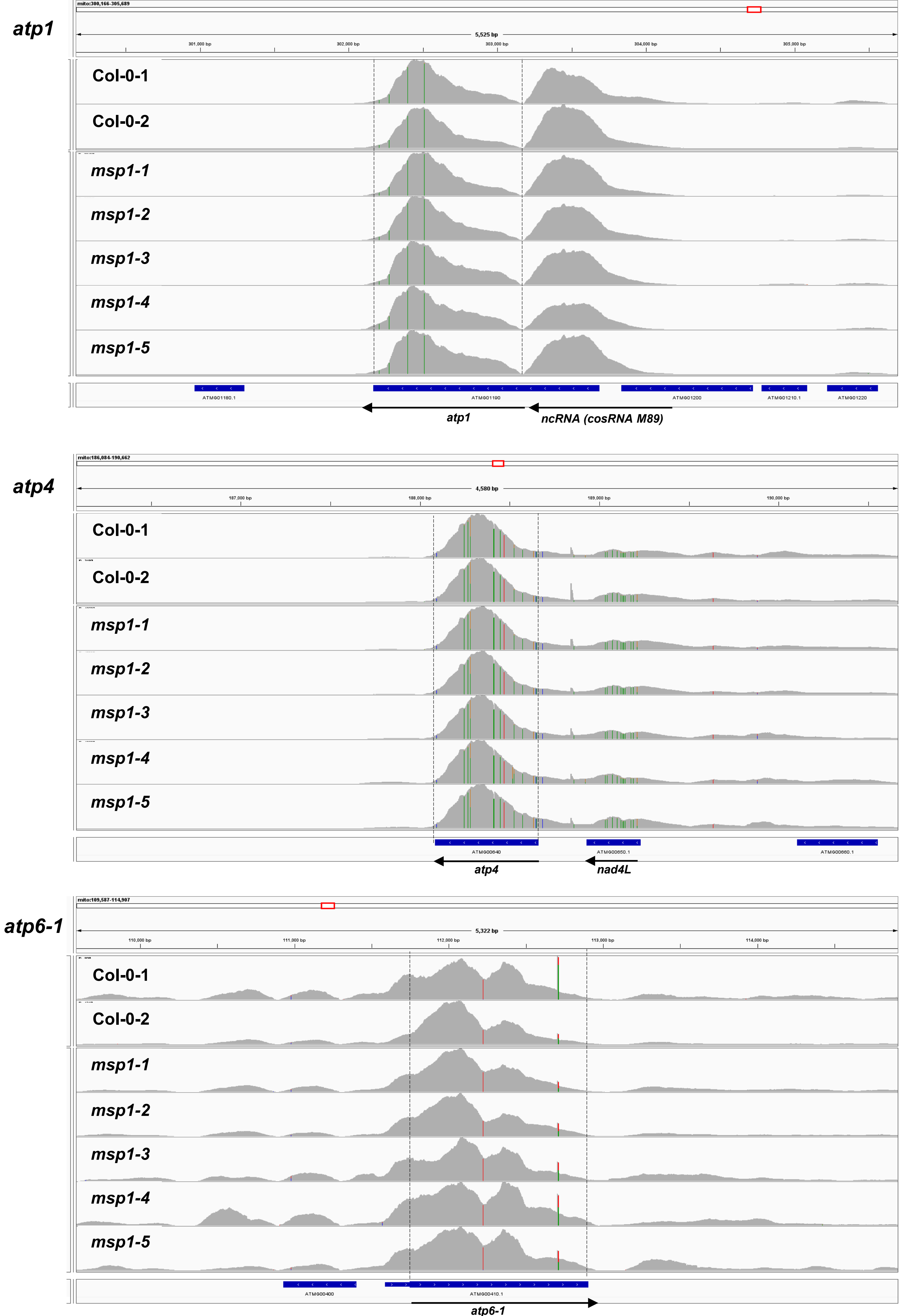

atp1
mito:300,166-305,689
5,525 bp
301,000 bp
302,000 bp
303,000 bp
304,000 bp
305,000 bp
Col-0-1
Col-0-2
msp1-1
msp1-2
msp1-3
msp1-4
msp1-5
ATMG01180.1
ATMG01190
ATMG01200
ATMG01210.1
ATMG01220
atp1
ncRNA (cosRNA M89)
atp4
mito:186,084-190,662
4,580 bp
187,000 bp
188,000 bp
189,000 bp
190,000 bp
Col-0-1
Col-0-2
msp1-1
msp1-2
msp1-3
msp1-4
msp1-5
ATMG00640
ATMG00650.1
ATMG00660.1
atp4
nad4L
atp6-1
mito:109,587-114,907
5,322 bp
110,000 bp
111,000 bp
112,000 bp
113,000 bp
114,000 bp
Col-0-1
Col-0-2
msp1-1
msp1-2
msp1-3
msp1-4
msp1-5
ATMG00400
ATMG00410.1
atp6-1

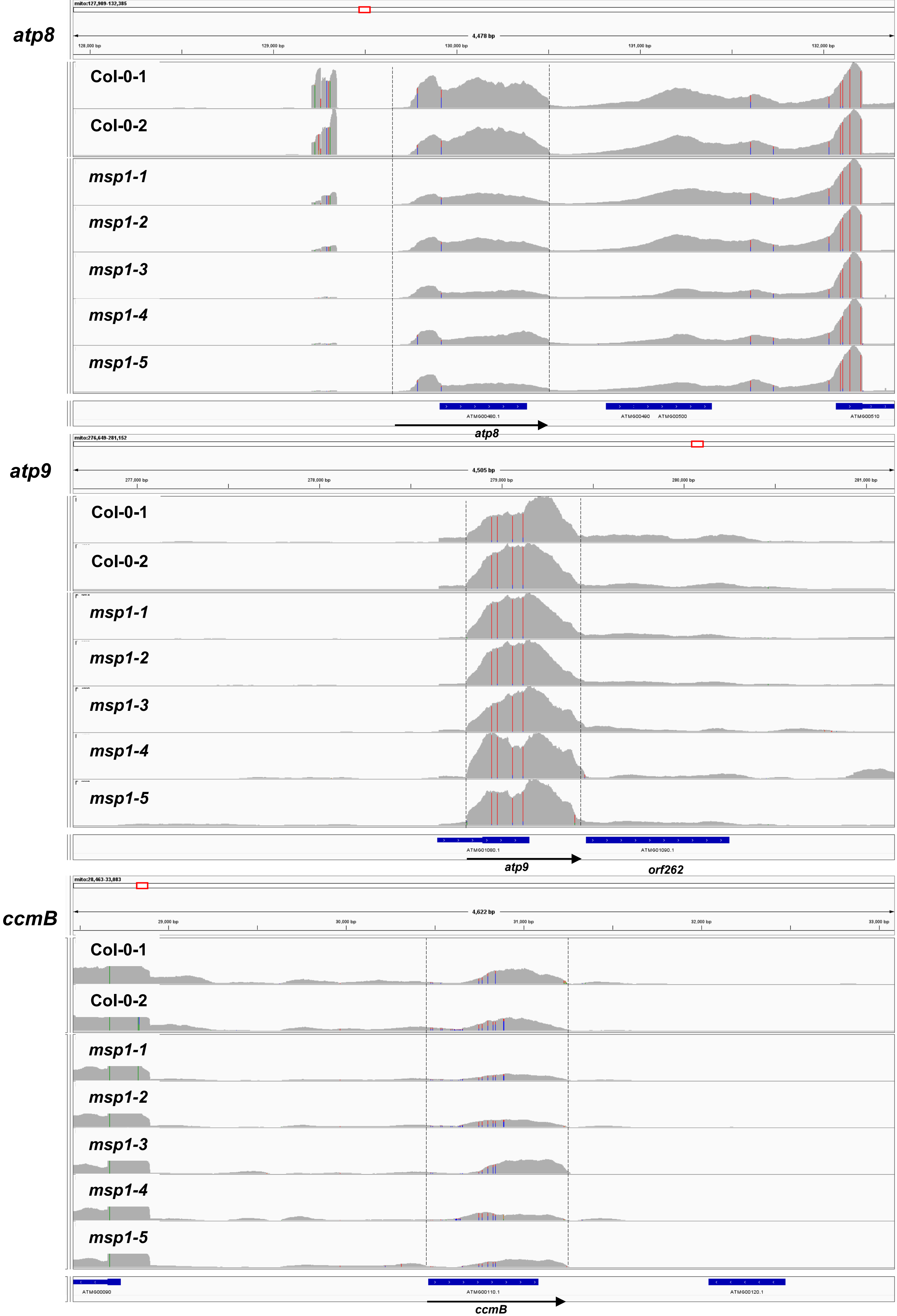

atp8
mito:127,909-132,385
4,478 bp
128,000 bp
129,000 bp
130,000 bp
131,000 bp
132,000 bp
Col-0-1
Col-0-2
msp1-1
msp1-2
msp1-3
msp1-4
msp1-5
ATMG00480.1
ATMG00490
ATMG00500
ATMG00510
atp8
atp9
mito:276,649-281,152
4,505 bp
277,000 bp
278,000 bp
279,000 bp
280,000 bp
281,000 bp
Col-0-1
Col-0-2
msp1-1
msp1-2
msp1-3
msp1-4
msp1-5
ATMG01080.1
ATMG01090.1
atp9
orf262
ccmB
mito:28,463-33,083
4,622 bp
29,000 bp
30,000 bp
31,000 bp
32,000 bp
33,000 bp
Col-0-1
Col-0-2
msp1-1
msp1-2
msp1-3
msp1-4
msp1-5
ATMG00090
ATMG00110.1
ATMG00120.1
ccmB

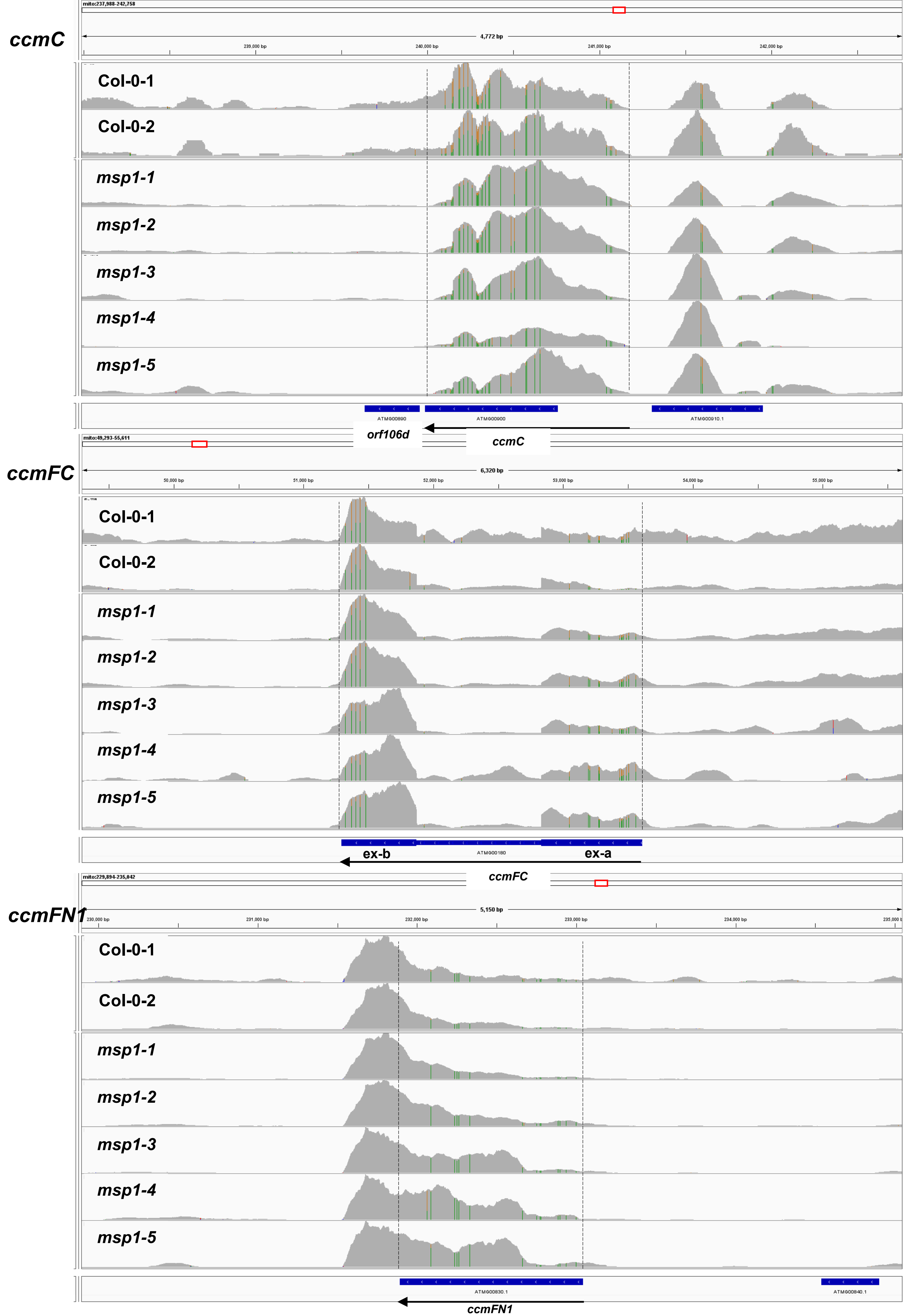
ccmC
mito:237,988-242,758
4,772 bp
239,000 bp
240,000 bp
241,000 bp
242,000 bp
Col-0-1
Col-0-2
msp1-1
msp1-2
msp1-3
msp1-4
msp1-5
ATMG00890
ATMG00900
ATMG00910.1
orf106d
ccmC
ccmFC
mito:49,293-55,611
6,320 bp
50,000 bp
51,000 bp
52,000 bp
53,000 bp
54,000 bp
55,000 bp
Col-0-1
Col-0-2
msp1-1
msp1-2
msp1-3
msp1-4
msp1-5
ex-b
ATMG00180
ex-a
ccmFC
ccmFN1
mito:229,894-235,042
5,150 bp
230,000 bp
231,000 bp
232,000 bp
233,000 bp
234,000 bp
Col-0-1
Col-0-2
msp1-1
msp1-2
msp1-3
msp1-4
msp1-5
ATMG00830.1
ATMG00840.1
ccmFN1

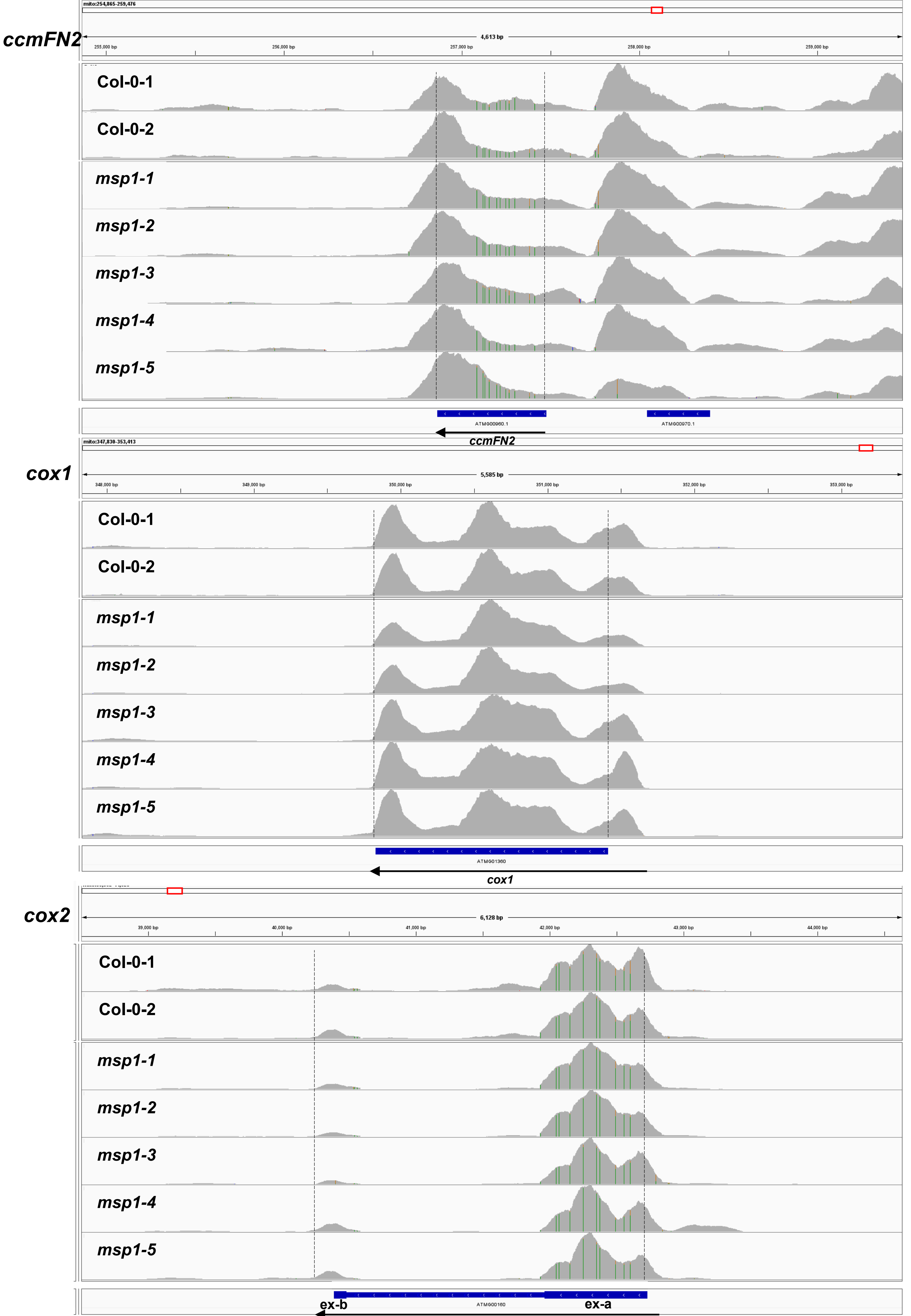

ccmFN2
mito:254,865-259,476
4,613 bp
255,000 bp
256,000 bp
257,000 bp
258,000 bp
259,000 bp
Col-0-1
Col-0-2
msp1-1
msp1-2
msp1-3
msp1-4
msp1-5
ATMG00960.1
ATMG00970.1
ccmFN2
cox1
mito:347,830-353,413
5,585 bp
348,000 bp
349,000 bp
350,000 bp
351,000 bp
352,000 bp
353,000 bp
Col-0-1
Col-0-2
msp1-1
msp1-2
msp1-3
msp1-4
msp1-5
ATMG01360
cox1
cox2
6,128 bp
39,000 bp
40,000 bp
41,000 bp
42,000 bp
43,000 bp
44,000 bp
Col-0-1
Col-0-2
msp1-1
msp1-2
msp1-3
msp1-4
msp1-5
ex-b
ATMG00160
ex-a

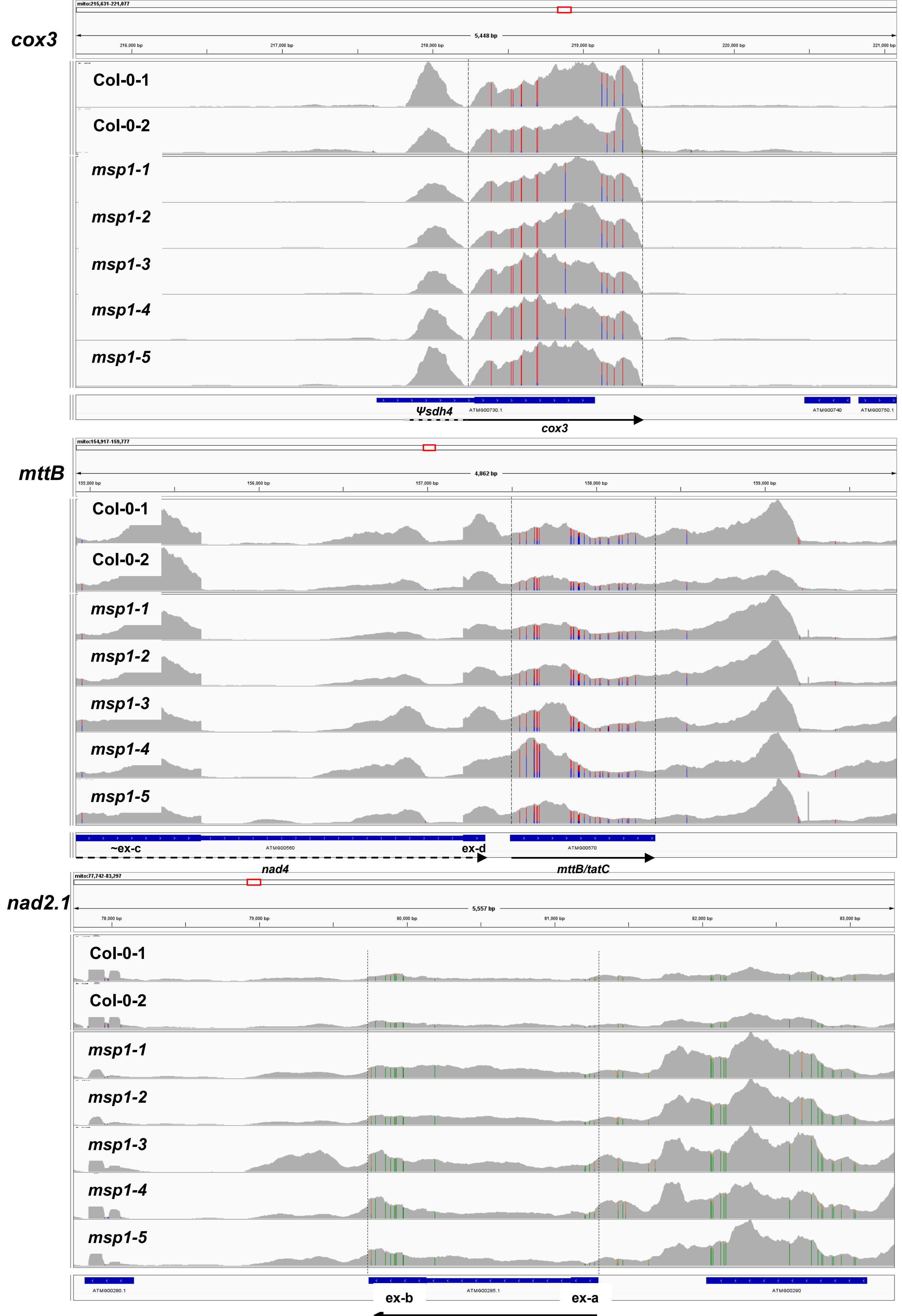
cox3
mito:215,631-221,077
5,448 bp
216,000 bp
217,000 bp
218,000 bp
219,000 bp
220,000 bp
221,000 bp
Col-0-1
Col-0-2
msp1-1
msp1-2
msp1-3
msp1-4
msp1-5
Ψsdh4
ATMG00730.1
ATMG00740
ATMG00750.1
cox3
mttB
mito:154,917-159,777
4,862 bp
155,000 bp
156,000 bp
157,000 bp
158,000 bp
159,000 bp
Col-0-1
Col-0-2
msp1-1
msp1-2
msp1-3
msp1-4
msp1-5
~ex-c
ATMG00560
ex-d
ATMG00570
nad4
mttB/tatC
nad2.1
mito:77,742-83,297
5,557 bp
78,000 bp
79,000 bp
80,000 bp
81,000 bp
82,000 bp
83,000 bp
Col-0-1
Col-0-2
msp1-1
msp1-2
msp1-3
msp1-4
msp1-5
ATMG00280.1
ATMG00285.1
ATMG00290
ex-b
ex-a

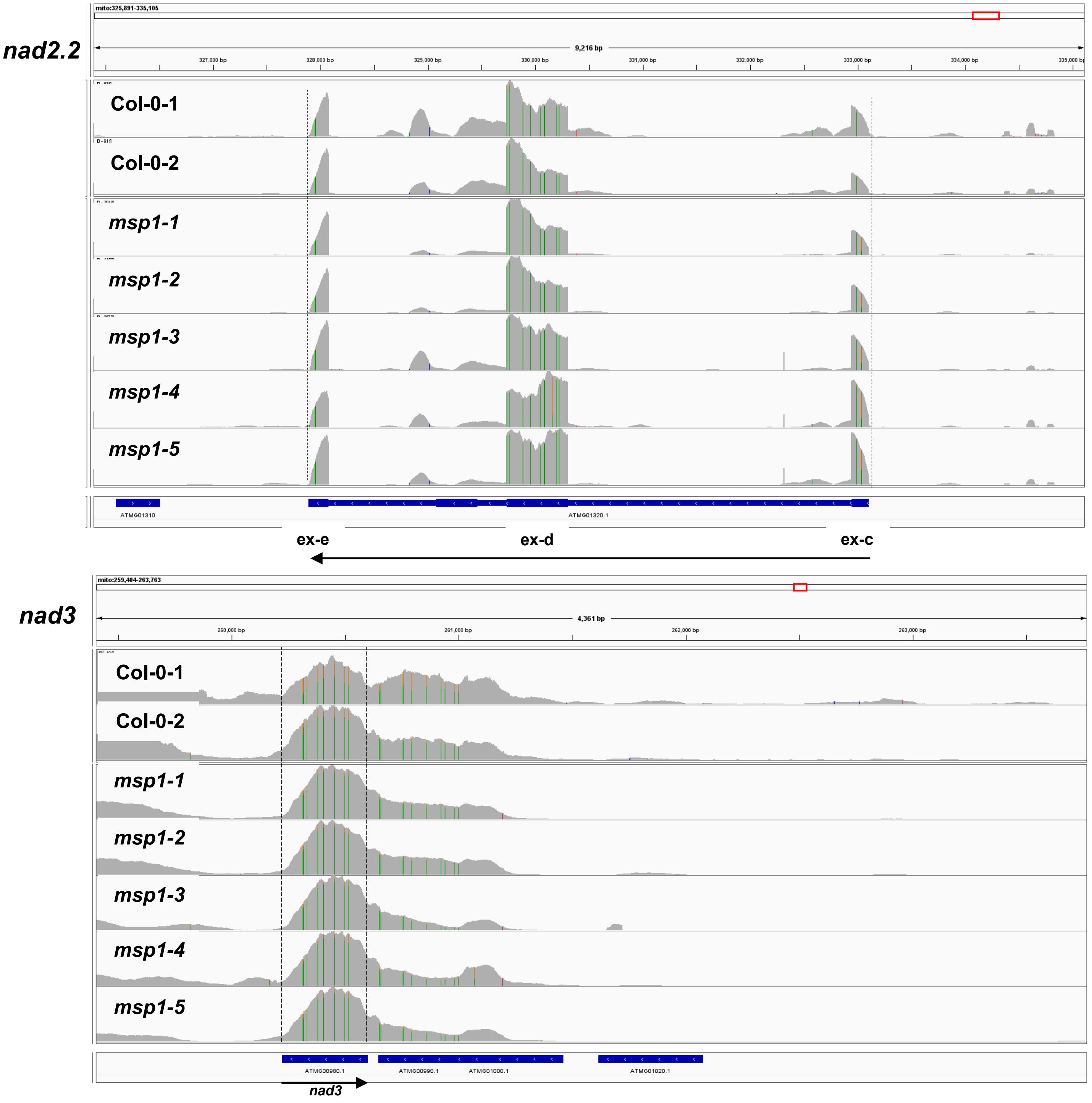
nad2.2
mito:325,891-335,105
9,216 bp
327,000 bp
328,000 bp
329,000 bp
330,000 bp
331,000 bp
332,000 bp
333,000 bp
334,000 bp
Col-0-1
Col-0-2
msp1-1
msp1-2
msp1-3
msp1-4
msp1-5
ATMG01310
ATMG01320.1
ex-e
ex-d
ex-c
nad3
mito:259,404-263,763
4,361 bp
260,000 bp
261,000 bp
262,000 bp
263,000 bp
Col-0-1
Col-0-2
msp1-1
msp1-2
msp1-3
msp1-4
msp1-5
ATMG00980.1
ATMG00990.1
ATMG01000.1
ATMG01020.1
nad3

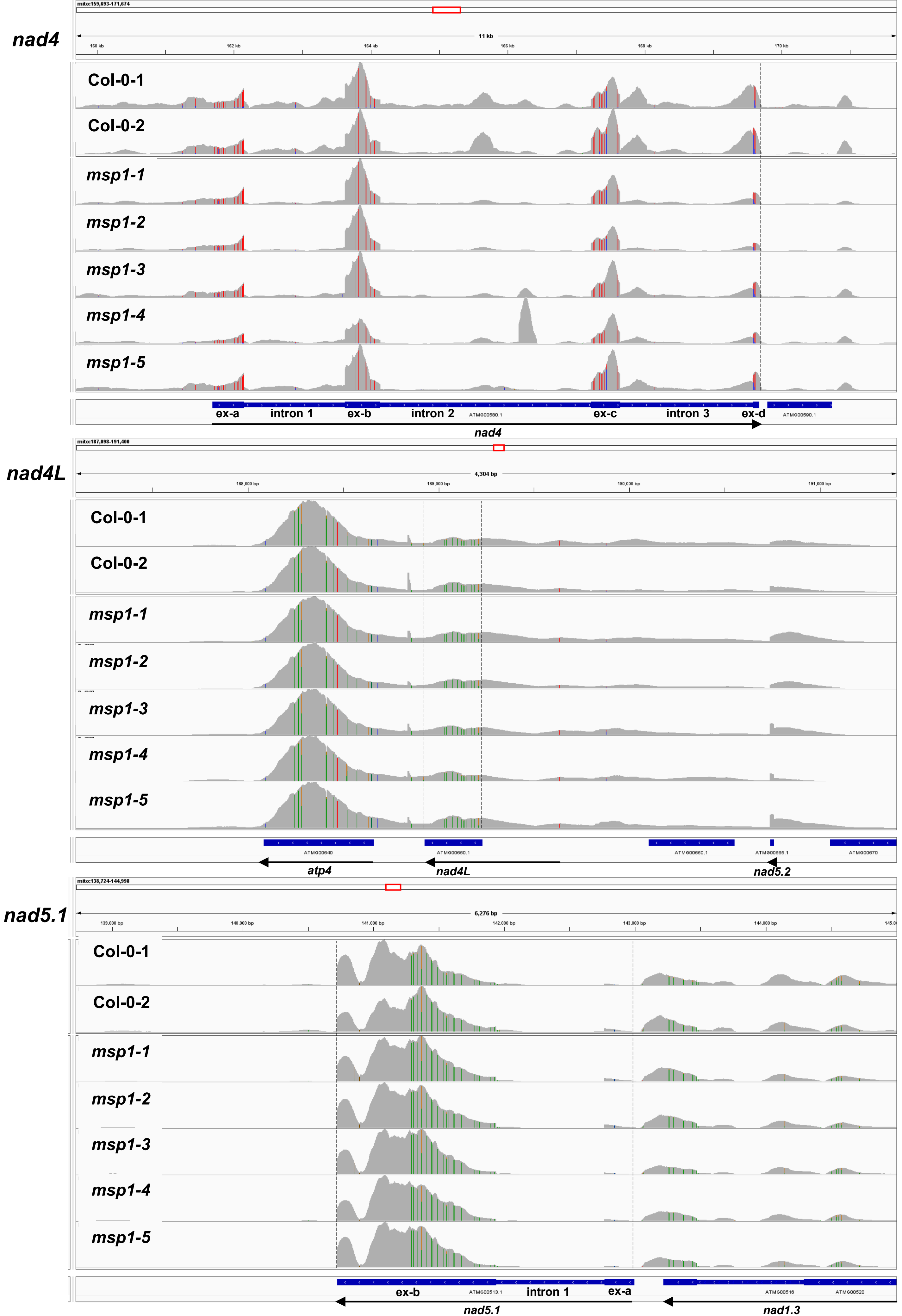
nad4
mito:159,693-171,674
11 kb
160 kb
162 kb
164 kb
166 kb
168 kb
170 kb
Col-0-1
Col-0-2
msp1-1
msp1-2
msp1-3
msp1-4
msp1-5
ex-a
intron 1
ex-b
intron 2
ATMG00580.1
ex-c
intron 3
ex-d
ATMG00590.1
nad4
nad4L
mito:187,098-191,400
4,304 bp
188,000 bp
189,000 bp
190,000 bp
191,000 bp
Col-0-1
Col-0-2
msp1-1
msp1-2
msp1-3
msp1-4
msp1-5
ATMG00640
ATMG00650.1
ATMG00660.1
ATMG00665.1
ATMG00670
atp4
nad4L
nad5.2
nad5.1
mito:138,724-144,998
6,276 bp
139,000 bp
140,000 bp
141,000 bp
142,000 bp
143,000 bp
144,000 bp
Col-0-1
Col-0-2
msp1-1
msp1-2
msp1-3
msp1-4
msp1-5
ex-b
ATMG00513.1
intron 1
ex-a
ATMG00516
ATMG00520
nad5.1
nad1.3

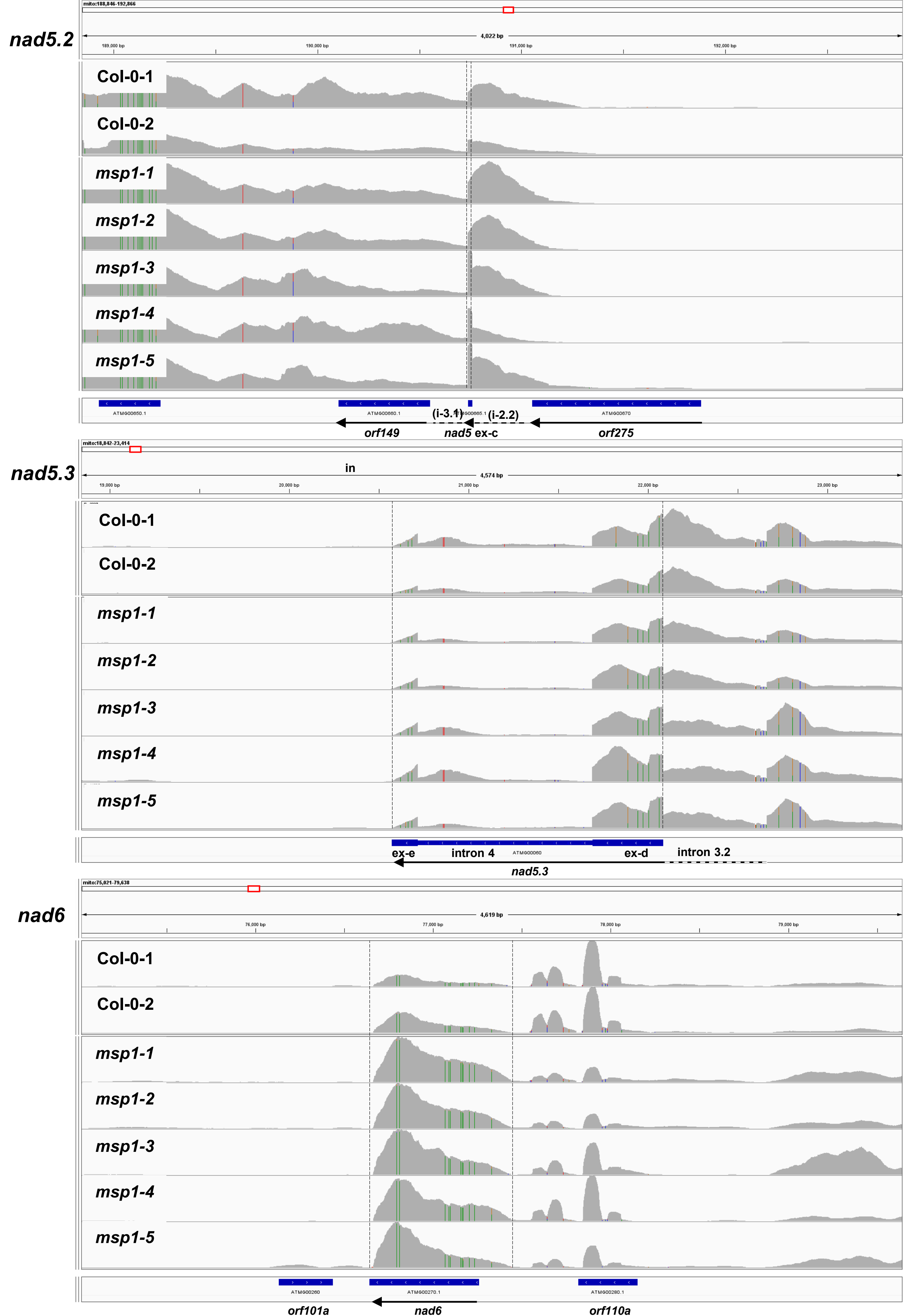

nad5.2
mito:188,846-192,866
4,022 bp
189,000 bp
190,000 bp
191,000 bp
192,000 bp
Col-0-1
Col-0-2
msp1-1
msp1-2
msp1-3
msp1-4
msp1-5
ATMG00650.1
ATMG00660.1
ATMG00665.1
ATMG00670
(i-3.1)
(i-2.2)
orf149
nad5 ex-c
orf275
nad5.3
mito:18,842-23,414
4,574 bp
19,000 bp
20,000 bp
21,000 bp
22,000 bp
23,000 bp
in
Col-0-1
Col-0-2
msp1-1
msp1-2
msp1-3
msp1-4
msp1-5
ex-e
intron 4
ATMG00060
ex-d
intron 3.2
nad5.3
nad6
mito:75,021-79,638
4,619 bp
76,000 bp
77,000 bp
78,000 bp
79,000 bp
Col-0-1
Col-0-2
msp1-1
msp1-2
msp1-3
msp1-4
msp1-5
ATMG00260
ATMG00270.1
ATMG00280.1
orf101a
nad6
orf110a

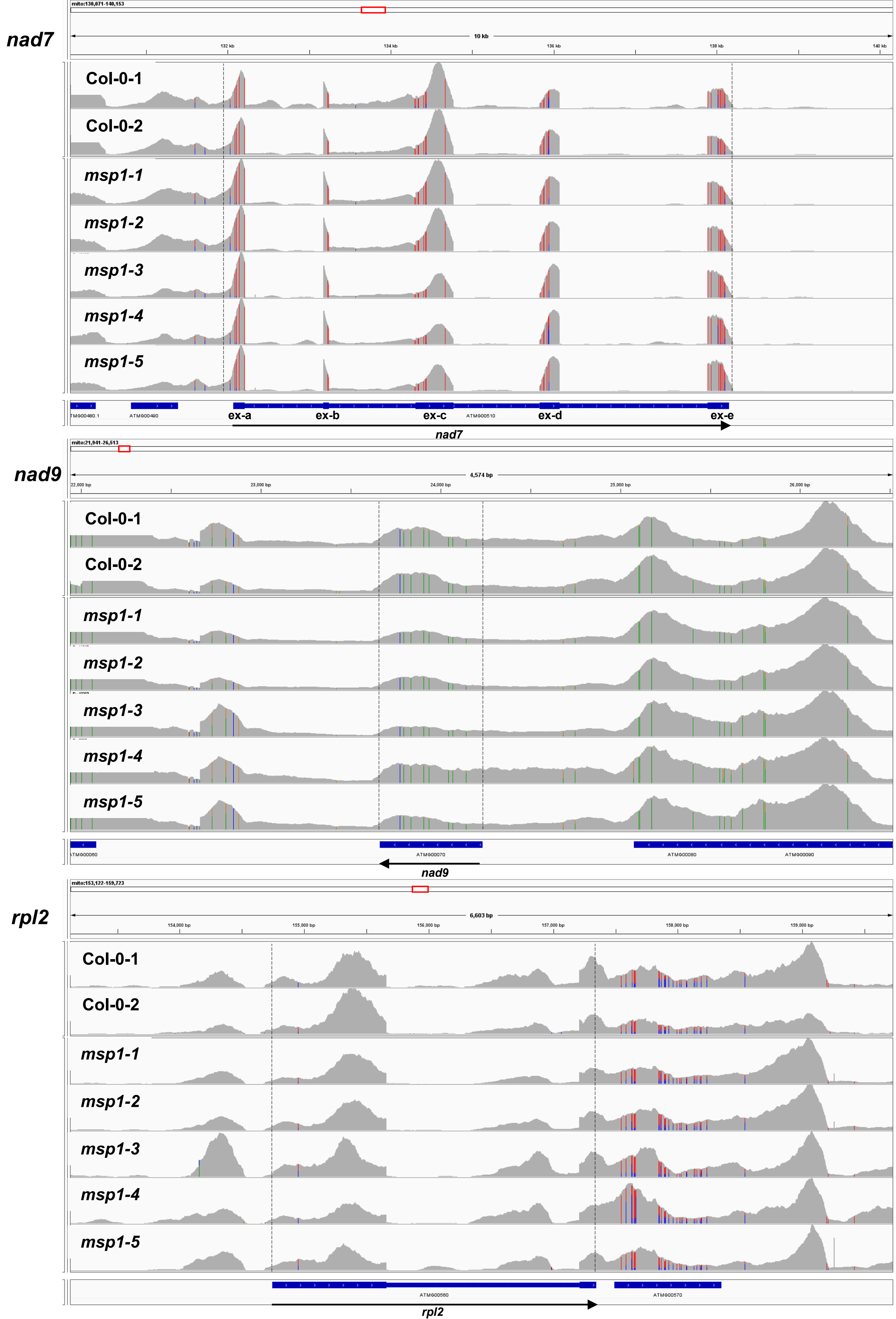
nad7
mito:130,071-140,153
10 kb
132 kb
134 kb
136 kb
138 kb
140 kb
Col-0-1
Col-0-2
msp1-1
msp1-2
msp1-3
msp1-4
msp1-5
TMG00480.1
ATMG00490
ex-a
ex-b
ex-c
ATMG00510
ex-d
ex-e
nad7
nad9
mito:21,941-26,513
4,574 bp
22,000 bp
23,000 bp
24,000 bp
25,000 bp
26,000 bp
Col-0-1
Col-0-2
msp1-1
msp1-2
msp1-3
msp1-4
msp1-5
TMG00060
ATMG00070
ATMG00080
ATMG00090
nad9
rpl2
mito:153,122-159,723
6,603 bp
154,000 bp
155,000 bp
156,000 bp
157,000 bp
158,000 bp
159,000 bp
Col-0-1
Col-0-2
msp1-1
msp1-2
msp1-3
msp1-4
msp1-5
ATMG00560
ATMG00570
rpl2

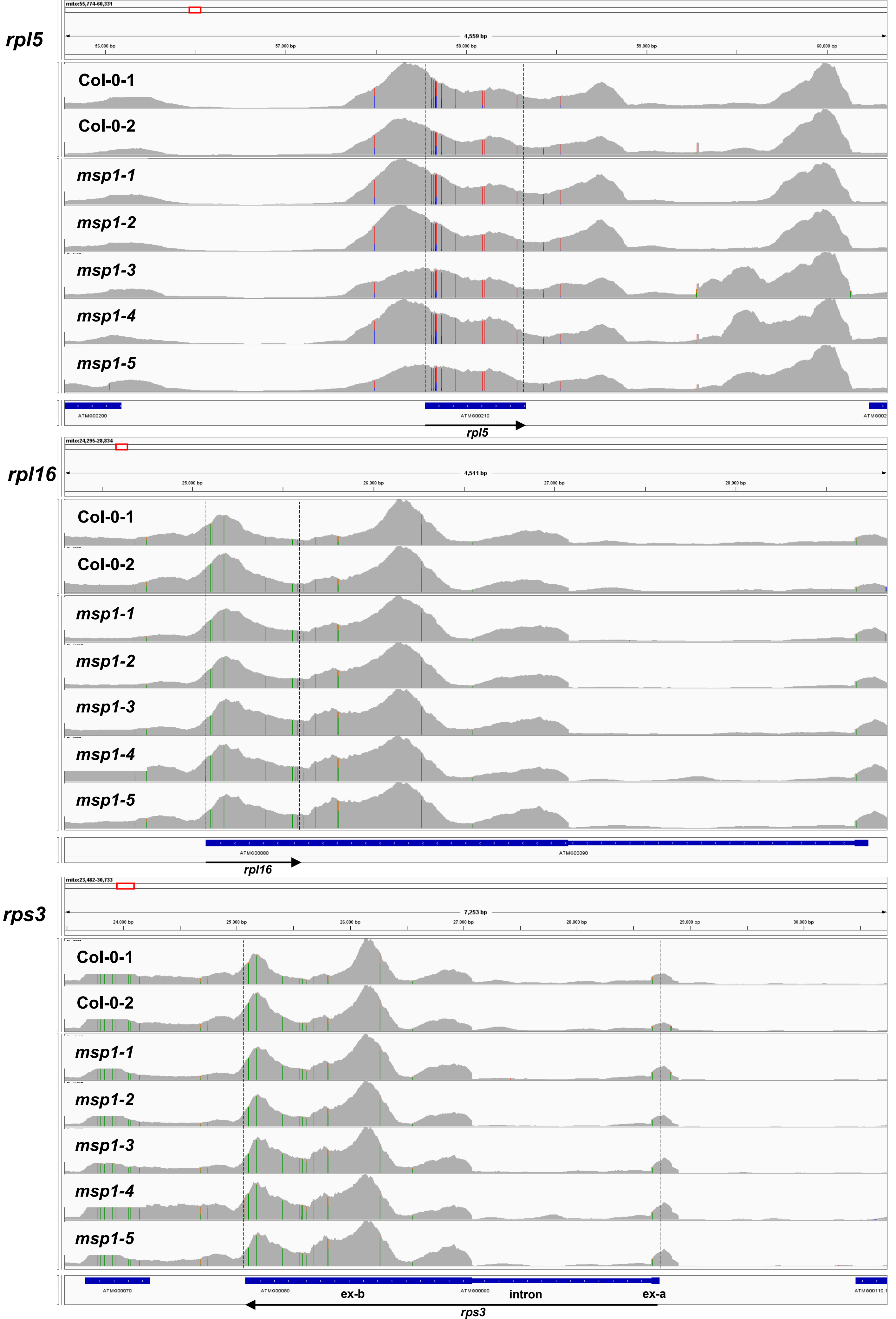
rpl5
mito:55,774-60,331
4,559 bp
56,000 bp
57,000 bp
58,000 bp
59,000 bp
60,000 bp
Col-0-1
Col-0-2
msp1-1
msp1-2
msp1-3
msp1-4
msp1-5
ATMG00200
ATMG00210
rpl5
rpl16
mito:24,295-28,834
4,541 bp
25,000 bp
26,000 bp
27,000 bp
28,000 bp
Col-0-1
Col-0-2
msp1-1
msp1-2
msp1-3
msp1-4
msp1-5
ATMG00080
ATMG00090
rpl16
rps3
mito:23,482-30,733
7,253 bp
24,000 bp
25,000 bp
26,000 bp
27,000 bp
28,000 bp
29,000 bp
30,000 bp
Col-0-1
Col-0-2
msp1-1
msp1-2
msp1-3
msp1-4
msp1-5
ATMG00070
ATMG00080
ex-b
ATMG00090
intron
ex-a
rps3

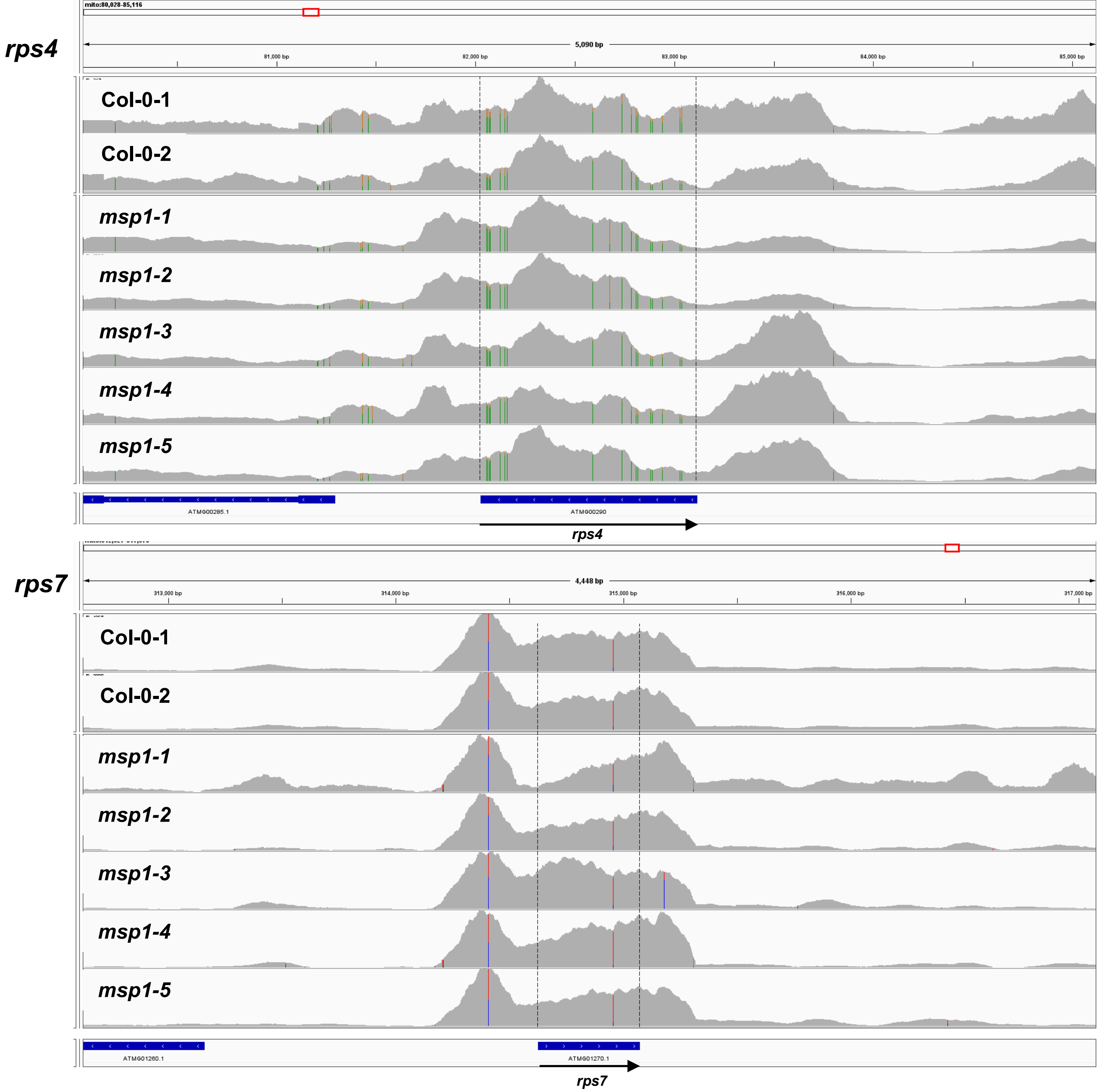
rps4
mito:80,028-85,116
5,090 bp
81,000 bp
82,000 bp
83,000 bp
84,000 bp
85,000 bp
Col-0-1
Col-0-2
msp1-1
msp1-2
msp1-3
msp1-4
msp1-5
ATMG00285.1
ATMG00290
rps4
rps7
4,448 bp
313,000 bp
314,000 bp
315,000 bp
316,000 bp
317,000 bp
Col-0-1
Col-0-2
msp1-1
msp1-2
msp1-3
msp1-4
msp1-5
ATMG01260.1
ATMG01270.1
rps7
