## Supplemental Figures and Tables for "*MSP1* encodes an essential RNA-binding PPR factor required for *nad1* maturation and complex I biogenesis in *Arabidopsis* mitochondria"

#### Supporting Information

**Figure S1. *MSP1* gene expression patterns at different tissues and during various developmental stages.**

The expression patterns of *MSP1* were analyzed by publicly available microarray and high throughput sequencing databases, including (A) Genevestigator analysis toolbox (Zimmermann *et al.*, 2004; Hruz *et al.*, 2008) and (B) The *Arabidopsis* Information Resource, TAIR; http://www.arabidopsis.org).

**Figure S2. MSP1 protein structure and its predicated RNA biding site.**

(A) The deduced amino acid sequence of MSP1. The predicted mitochondrial targeting sequence (36~37 amino acid long) is highlighted in magenta, with the cleavage sites indicated by box. Regions of low confidence are colored in grey. Predicted PPR motifs are indicated (domains of low scoring are highlighted in cyan and coded blue for regions of high confident). (B) Homology modeling of *Arabidopsis thaliana* (Col-0) MSP1. The 3D model of the MSP1 protein structure was obtained from AlphaFold server (Jumper *et al.*, 2021), using a per-residue confidence score (pLDDT) (indicated below the figure). The hypothetical 3D surface of MSP1 was visualized with PyMol. The "I" and "IV" panels represent ribbon structures at different angles, whereas "II" and "III" indicate the hypothetical surface of MSP1. The color code is red for negative values, white for near zero values and blue for positive values.

**Figure S3. PCR screening of *msp1* plants.**

(A) The nucleotide sequence of *MSP1* encoded by *AT4G20090*. Lower case indicates the 5' and 3' untranslated regions (UTRs), while uppercase letters represent the open reading frame of *MSP1* (as indicated by the TAIR database). The reported positions of T-DNA insertions in *msp1-1* (SALK-142675), SALK-018927 and SALK-070654 lines are indicated in the sequence of *MSP1*. PCR followed by sequencing suggest a double insertion event for the *msp1-1* line within the coding region of *MSP1*. Analyses of genomic PCR products spanning the reported T-DNA insertion junctions failed to support T-DNA insertions in the *MSP1* in SALK-lines 018927 and 070654. (B) PCR screening of germinated plantlets obtained from mature seeds of heterozygous *msp1-1* mutants, with primers designed to the *MSP1* gene and SALK T-DNA left border primer (LBb1.3). Arrows indicate the 465 bp and 541 bp products that correspond to the gene fragment and the *T-*

*DNA-MSP1* insertional product, respectively. The nucleotide sequence of the precise insertion position of *msp1-1* is presented in Fig. S3. (C) PCR screening of homozygous *msp1-1* mutants carrying the 35S:*MSP1* transgene (*i.e.*, complemented lines). Specific primers used for the screening of complemented *msp1-1* lines are listed in Table S5.

**Figure S4. The mitochondrial transcription profiles of *msp1-1*.**

(A) Transcriptomic data are shown for mtDNA loci corresponding to the *nad1.1* (nucleotides 81,785 - 83,966, indicated in blue color), *nad1.2* (nucleotides 58,315 - 60,868, indicated in green) and *nad1.3* (nucleotides 230,304 - 237,530, indicated in red) gene fragments. The *nad1.1* and *nad1.2* fragments are separated by domain 4 of intron 1, while the *nad1.2* and *nad1.3* transcription units are fragmented by domain 4 of intron 3. The structure of *nad1* i1 intron was modeled based on previous structural models of group II intron RNAs (Dai *et al.*, 2003; Candales *et al.*, 2011), structural analyses of bacterial MATs bound to their cognate group II intron RNAs (Piccirilli & Staley, 2016; Qu *et al.*, 2016; Zhao & Pyle, 2016), and the secondary structure prediction algorithms MFOLD (Zuker, 2003) and Alifold (Hofacker *et al.*, 2002). The figure includes 16 nucleotides of the 5' exon and 1,151 nucleotides of the intron from *nad1.1* transcript (in blue color) plus 108 nucleotides of the 3' part of the intron and 11 nucleotides of the 3' exon from *nad1.2* transcript (in green color). Residue numbering begins at the first intron nucleotide (*i.e.*, 5'-part). Nucleotides predicted to be involved in the canonical group II intron tertiary interactions, including the IBS1-EBS1, IBS2-EBS2, α-α', β-β', γ-γ', δ-δ', ε-ε', ζ-ζ', κ-κ', λ-λ', θ-θ' and the η-η' interactions are indicated, while the predicted RNA binding site of MSP1 (5'-acaUACCAAAAGGUu, Fig. S5) is highlighted. The predicted bulged A cleavage site (Li-Pook-Than & Bonen, 2006) is indicated in bold red color in domain VI. (B) For the RNA-seq analyses, total mtRNA was extracted from individual Col-0 and embryo-rescued *msp1-1* plantlets (seven runs corresponding to 3 biological repeats), which was used to generate Illumina-sequencing libraries (BioProjects PRJNA704631 and PRJNA768306). The alignments between Col-0 or *msp1-1* plants(154) and the established *Arabidopsis* mtDNA (Sloan *et al.*, 2018) were visualized using the IGV genome browser (Thorvaldsdottir *et al.*, 2013). Green and red (-strand) or blue and red (+strand) lines point to different mtRNA-editing (C-to-U conversions) events established for exon regions in *Arabidopsis* mitochondria (Sloan *et al.*, 2018). Arrows show the direction of transcription. Boxes represent different exons or ORF regions, while lines indicate to group II intron regions. Dashed lines represent UTR regions (Best, Corinne *et al.*, 2020). The lower panel indicates read coverage in the *nad1* exon-a region in Col-0 (blue) and *msp1* (orange) plants.

**Figure S5. Analysis of MSP1 RNA binding site.**

(A) A combinatorial code for RNA recognition by the predicted MSP1 PPR motifs. Amino acids at positions 5 and the end of each domain act as specificity-determining residues (Barkan *et al.*, 2012; Takenaka *et al.*, 2013; Cheng *et al.*, 2016). *In-silico* data (Cheng *et al.*, 2016; Harrison *et al.*, 2016; Shen *et al.*, 2019) indicate the preferred sequence 5'-UACCAAAAGGU for MSP1's PPR 4-11 motifs. (B) The PPRmatcher algorithm (Royan *et al.*, 2021), which aligns the predicated PPR motifs to every possible position in the mitogenome of Arabidopsis (i.e., ~720K positions) and scores each alignment based on the data in (Yan *et al.*, 2019).

**Figure S6. Expression and purification of a recombinant rMSP1 protein.**

(A) The codon optimized region of the PPR domains 3 to 14 of MSP1 was cloned into pGEX-4T1, in-frame to GST in its N-terminus and a C-terminal 6xHis tag. The recombinant protein (rMSP1, 71,365 kDa) was expressed in *E. coli* (strain BL21 cells), and induced with 1.0 mM IPTG for 2 h at 37°C. The cells were pelleted, resuspended with in a 6M guanidine hydrochloride buffer, and lysed by a high-pressure chamber manifold. Non-soluble materials were removed by centrifugation. Immunoblots with anti-GST antibodies and protein MS-analysis confirmed the integrity of the over-expressed recombinant protein. (B) The rMSP1 was purified by Ni-NTA column under denaturing conditions, using the BioRad Profinia (Petersen, 2007) purification system and column kit.

**Figure S7. The relative accumulation of organellar proteins and respiratory complexes in wild-type and *msp1-1* plants.**

(A) Immunoblots with proteins (about 50 μg) of mitochondrial enriched membranous fractions (Pineau *et al.*, 2008; Sultan *et al.*, 2016), isolated from MS-grown Col-0 and rescued *msp1-1* plants. The blots were probed with polyclonal antibodies (Table S4) raised against Rieske iron-sulfur protein (RISP) of complex III (CIII), the cytochrome oxidase subunit 2 (Cox2) of complex IV (CIV), mitochondrial Atp1 and ATP2 subunits of ATP-synthase (CV) and the mitochondrial voltage-dependent anion channel (VDAC, or porin). Detection was carried out by chemiluminescence assays after incubation with HRP-conjugated secondary antibody (note that the box in the RISP blot is a cropped image of the lanes that correspond to *msp1*, so that the heterozygous and homozygous lines are found at the same loading order as in the other blots; no other changes have been made to the original blots). (B) Analyses of the expression levels of various alternative oxidases (AOXs) and rotenone-insensitive NAD(P)H dehydrogenases (NDs) in *Arabidopsis* plants was performed by RT-qPCR, as described previously (Zmudjak *et al.*, 2013; Cohen *et al.*, 2014; Sultan *et al.*, 2016). RNA extracted from wild-type (Col-0) and embryo-rescued *msp1-1* mutant was reverse-transcribed, and the relative steady-state levels of cDNAs corresponding to the different *AOX* and *ND* transcripts were evaluated by qPCR with primers which specifically amplified their mRNAs (Table S1). The histogram shows the relative mRNAs levels in *msp1-1* and rescued Col-0 plantlets versus 3-week-old MS-grown Col-0 plants. (C) Blue native (BN) PAGE analyses of crude organellar membranous fractions. An aliquot equivalent to 40 mg of crude *Arabidopsis* mitochondria extracts, obtained from wild-type plants, *msp1* mutant-lines and complemented *msp1:35S-MSP1*, was solubilized with 5% (w/v) digitonin, and the organellar complexes were resolved by BN-PAGE. For immunodetection, the proteins were transferred from the native gels onto a PVDF membrane. The membranes probed with antibodies raised to Rieske iron-sulfur protein (RISP) of complex III (CIII), the cytochrome oxidase subunit 2 (Cox2) of complex IV (CIV), mitochondrial ATP2 subunit of ATP-synthase (CV) (Table S4), as indicated below each blot. Arrows indicate to the native supercomplex I+III2 (~1,500 kDa), $III_2$ (~500 kDa), CIV (~220 kDa) and CV (~600 kDa).

**Figure S8. Relative accumulation and activity of respiratory CI in wild-type and *nmat1* plants.**

BN-PAGE of crude organellar membranous fractions was performed generally as described previously (Pineau *et al.*, 2008). An aliquot equivalent to 40 mg of crude *Arabidopsis* mitochondria extracts, obtained from wild-type (Col-0) plants and *nmat1* mutants (Nakagawa & Sakurai, 2006; Keren *et al.*, 2012), was solubilized with 5% (w/v) digitonin and the organellar complexes were resolved by BN-PAGE. For immunodetection, the proteins were transferred from the native gels onto a PVDF membrane. The membranes were destained with ethanol before probing with specific antibodies (Table S4), as indicated below each blot. *In-gel* complex I activity assays were performed essentially as described previously (Meyer*, et al.* 2009). Arrows indicate the native supercomplex I+III2 (~1,500 kDa) and holo-complex I (~1,000 kDa). The asterisks in the CA2 panel indicate the presence of partial CI assembly intermediates (I*), with calculated masses of about 800, 750, 700, 650, 600 and 350 kDa. Note that the four lanes in each of the immunoblots were loaded and separated on the same gels. We cropped the images to remove lanes corresponding to another mutant line. No other modifications were done to the original images.

**Figure S9. Splicing efficiencies in *ca2* mutants.**

The relative accumulation of mRNA and pre-RNA transcripts in wild-type and *ca2* plants (Soto *et al.*, 2015; Cordoba *et al.*, 2016; Fromm *et al.*, 2016a; Fromm *et al.*, 2016b; Ci Rdoba *et al.*, 2019), corresponding to the 23 group II intron sequences in *Arabidopsis*, was evaluated by RT-qPCR, essentially as described previously (Cohen *et al.*, 2014; Sultan *et al.*, 2016). RNA extracted from Col-0 and mutant plants was reverse-transcribed and the relative steady-state levels of cDNAs corresponding to the different organellar transcripts were evaluated by qPCR with primers which specifically amplified pre-RNAs and mRNAs. The histogram shows the splicing efficiencies as indicated by the log2 ratios of pre-RNA to mRNA transcript abundance in *ca2* mutants, compared with those of 3-week-old MS-grown wild-type plants. The values are means of six replicates (error bars indicate one standard deviation).

**Figure S10. The predicted MSP1 binding site is highly conserved among different spermatophytes.**

Alignment of homologous mitochondrial *nad1.1* sequences from *Arabidopsis thaliana (A.t.)*, *Ceratozamia hildae (C.h.)*, *Cycas taitungensis (C.t.)*, *Ginkgo biloba (G.b.)*, *Glycine max (G.ma.)*, *Gnetum montanum (G.mo)*, *Nicotiana tabacum (N.t.), Oryza sativa (O.s.)*, *Solanum lycopersicum (S.h.)*, *Triticum aestivum (T.a.), Zamia integrifolia (Z.i.)* and *Zea mays (Z.m.)*, was conducted with T-Coffee multiple sequence alignment server and displayed using GeneDoc (http://nrbsc.org/gfx/genedoc), with the conserved residue shading mode.

**Figure S11. Phylogenetic analysis of homologou**s **MSP1 proteins in plants.**

Homologues sequences to the MSP1 protein were obtained by the Basic Local Alignment Search Tool (BLAST), against the plant genome sequence databases. A phylogeny tree (A) was constructed, using the MAFFT multiple sequence alignment server (Katoh *et al.*, 2017), using the genome sequences of different plant species (bootstrap values for 1,000 bootstrap replicates). These include the *A. thaliana* MSP1 protein (NP_001320003.1, labeled in white), and its putative homologs in different *Brassicaceae*, as well as in *Amygdaloideae*, *Anacardiaceae*, *Betulaceae*, *Citrus*, *Cleomacea*, *Dipterocarpaceae*, *Euphorbiaceae*, *Fabaceae*, *Fagaceae*, *Malvaceae*, *Poaceae*, *Rosoideae*, *Salicaceae*, *Sapindaceae*, *Solanaceae*, *Theaceae* and *Vitaceae*. The graphics at the end of the branches indicate to the specific plant family. (B) Alignment of homologous MSP1 and DEK2 related proteins from *A. thaliana*, *B. rapa*, *B. distachyon*, *H. vulgare*, *J. microsperma*, *N. tabacum*, *O. sativa*, *P. sitchensis*, *P. alba*, *S. lycopersicum*, *T. aestivum* and *Z. mays*, was conducted with T-Coffee multiple sequence alignment server, and displayed using GeneDoc (http://nrbsc.org/gfx/genedoc), with the conserved residue shading mode.

**Table S1.** Lists of oligonucleotides used for the analysis of the splicing profiles of wild-type and mutant plants by RT-qPCR experiments.

**Table S2.** List of splicing factors in *Arabidopsis* and their mtRNA profiles.

**Table S3.** List of antibodies used for the analysis of Col-0 and *msp1* mutants.

**Table S4.** Relative accumulation of mitochondrial proteins in wild-type and *msp1* plants.

**Table S5.** Oligonucleotides used in screening of individual T-DNA insertion lines in *Arabidopsis* and cloning of the *MSP1-GFP* gene-fusion construct.

A

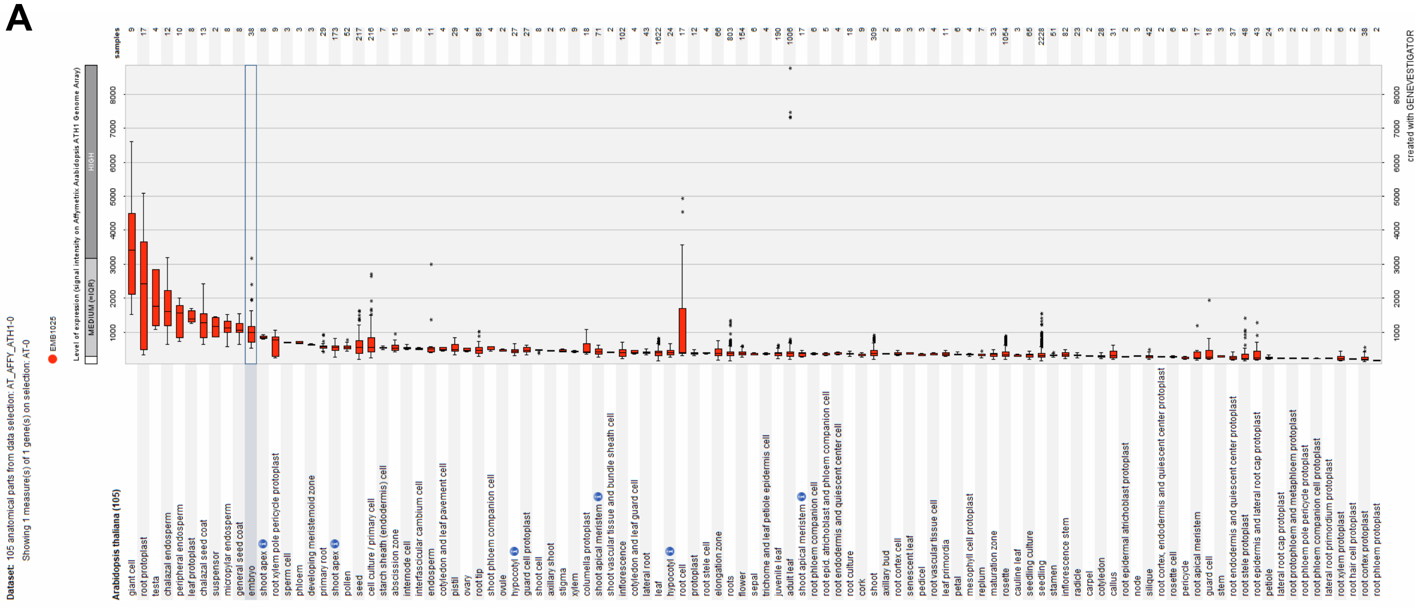

B

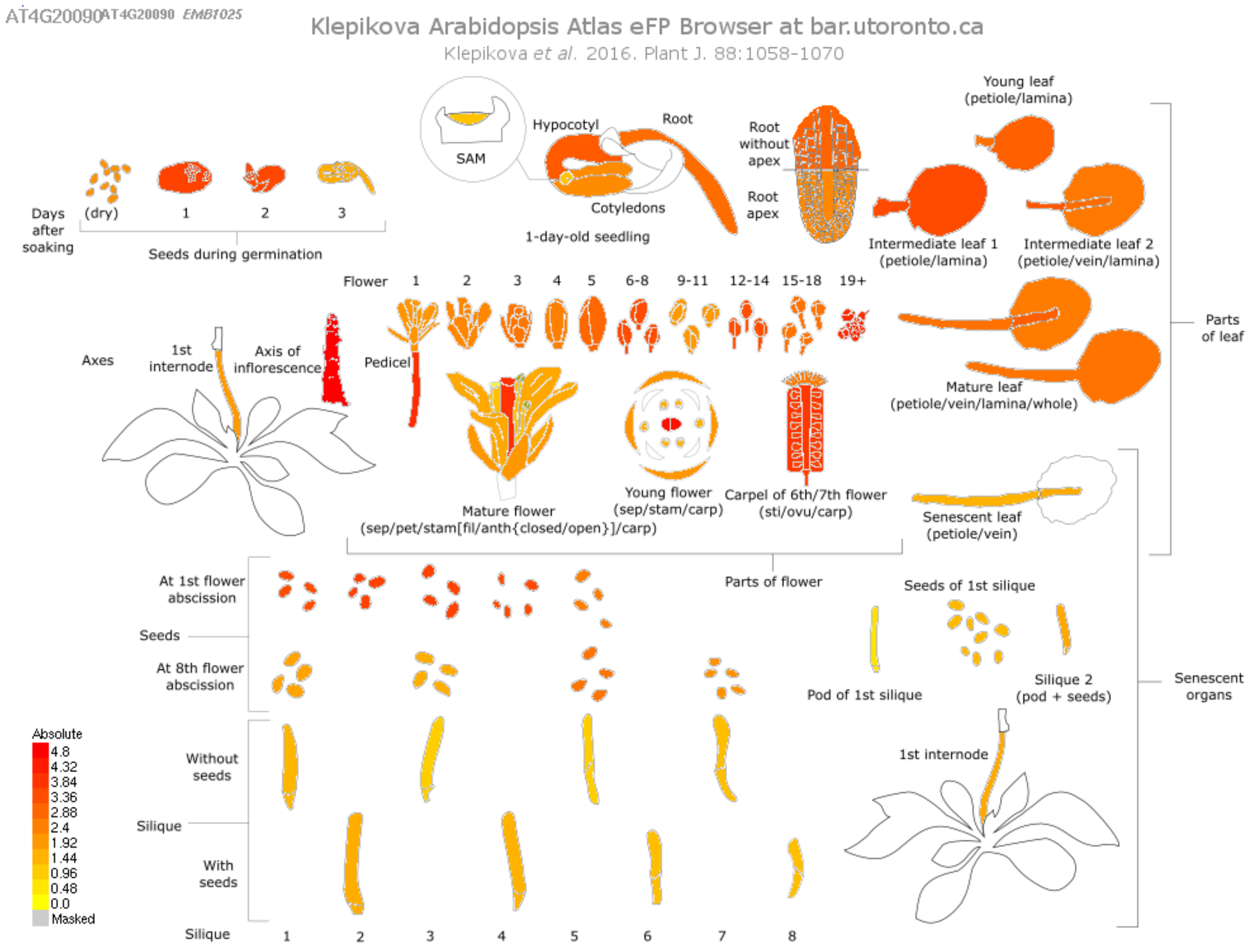
AT4G20090 AT4G20090 EMB1025
Klepikova Arabidopsis Atlas eFP Browser at bar.utoronto.ca
Klepikova et al. 2016. Plant J. 88:1058-1070
SAM
Hypocotyl
Root
Cotyledons
1-day-old seedling
Root without apex
Root apex
Young leaf (petiole/lamina)
Intermediate leaf 1 (petiole/lamina)
Intermediate leaf 2 (petiole/vein/lamina)
Days after soaking
(dry)
1
2
3
Seeds during germination
Flower
1
2
3
4
5
6-8
9-11
12-14
15-18
19+
Axes
1st internode
Axis of inflorescence
Pedicel
Mature flower (sep/pet/stam[fil/anth{closed/open}]/carp)
Young flower (sep/stam/carp)
Carpel of 6th/7th flower (sti/ovu/carp)
Mature leaf (petiole/vein/lamina/whole)
Parts of leaf
Senescent leaf (petiole/vein)
Parts of flower
At 1st flower abscission
Seeds
At 8th flower abscission
Seeds of 1st silique
Pod of 1st silique
Silique 2 (pod + seeds)
Senescent organs
1st internode
Absolute
4.8
4.32
3.84
3.36
2.88
2.4
1.92
1.44
0.96
0.48
0.0
Masked
Without seeds
Silique
With seeds
Silique
1
2
3
4
5
6
7
8

Figure S1

**A**

predicted MPP/ICP55 cleavage sites

MPKCPIPIRISFFSYFLKESRILSSNPVNFSIHLRFSSSVSVSPNPSMEVVENPLEAPISEKMFKSAPKMGSFKLG|DSTLSSMIESYANSGDFDSVE
KLLSRIRLENRVII|ERSFIVVFRAYGKAHLPDKAVDLFHRMVDEFRCKR|SVKSFNSVLNVIINEGLYHRGLEFYDYVVNSNMNMNISPN|GLSFNL
VIKALCKLRFVDRAIEVFRGMPERKCLPD|GYTYCTLMDGLCKEERIDEAVLLLDEMQSEGCSPS|PVIYNVLIDGLCKKGDLTRVTKLVDNMFLKGC
VPN|EVTYNTLIHGLCLKGKLDKAVSLLERMVSSKCIPN|DVTYGTLINGLVKQRRATDAVRLLSSMEERGYHLN|QHIYSVLISGLFKEGKAEEAMS
LWRKMAEKGCKPN|IVVYSVLVDGLCREGKPNEAKEILNRMIASGCLPN|AYTYSSLMKGFFKTGLCEEAVQVWKEMDKTGCSRN|KFCYSVLIDGLC
GVGRVKEAMMVWSKMLTIGIKPD|TVAYSSIIKGLCGIGSMDAALKLYHEMLCQEEPKSQPD|VVTYNILLDGLCMQKDISRAVDLLNSMLDRGCDPD
|VITCNTFLNTLSEKSNSCDKGRSFLEELVVRLLKQ|RVSGACTIVEVMLGKYLAPKTSTWAMIVREICKPKKINAAIDKCWRNLCT

**B**

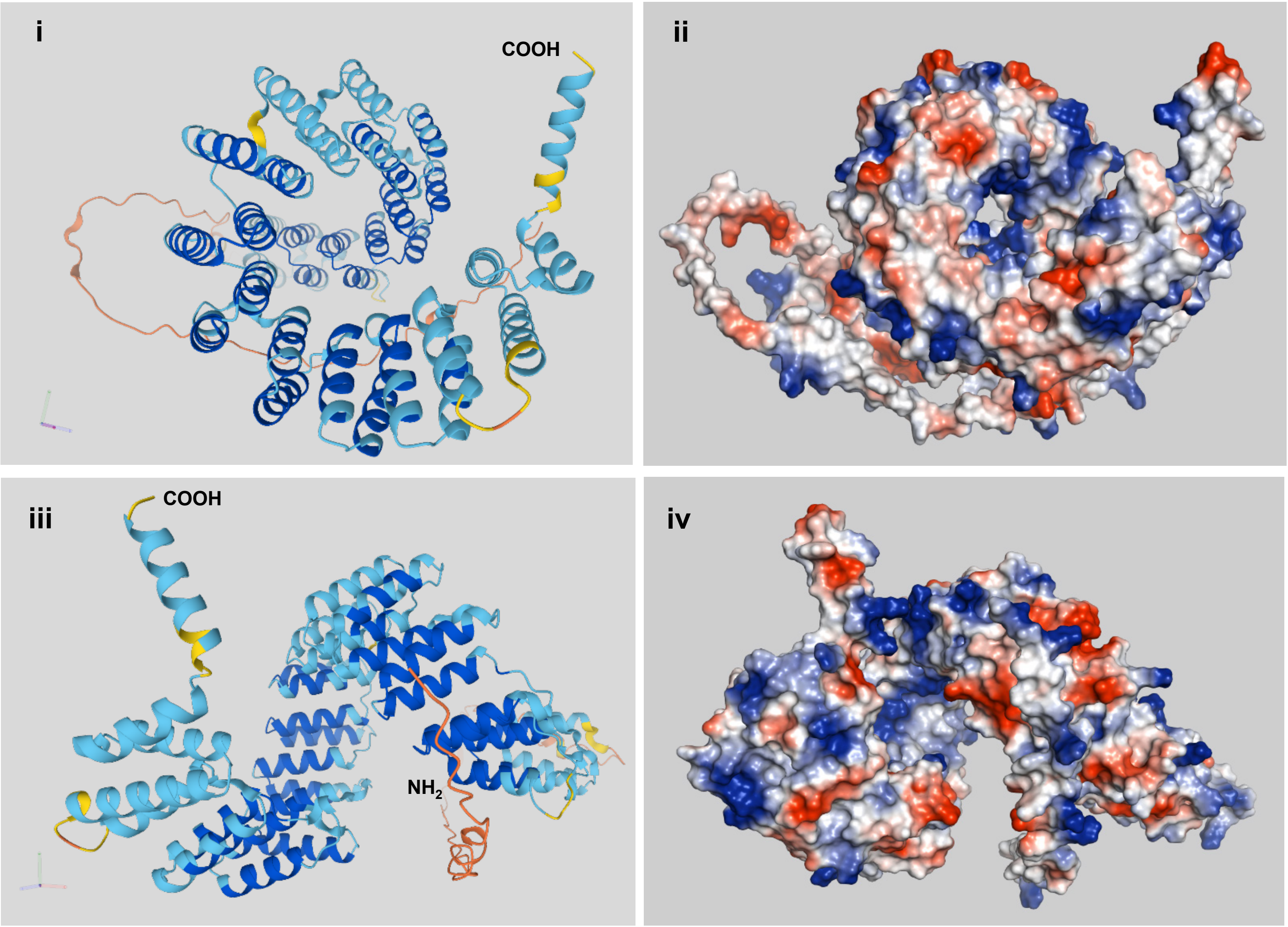


**AlphaFold Model Confidence :**

- Very high (pLDDT > 90)
- Confident (90 > pLDDT > 70)
- Low (70 > pLDDT > 50)
- Very low (pLDDT < 50)

**Figure S2**

**A.** **Arabidopsis *MSP1* gene**

5′…tatatagacatataaattagtatatataaatttgtttacttttaagattttctattttttagagaataaaaagttaaattt
tgaatgacacatgtaaacatctgatcgtttgatttgacacgtggcacaatcctatgagttagcaattttgaaaccatatattttat
ataataagatccccaaaattattatgtatatgagattatttaccatttggatttgattacataaattttataaattaaaaaattat
tggtaaatgaattagaaaatttcactttttaatagtaattaagaagtaattaatcaattaattgagaggaaacaaaaaaattgaaat
ttattgataactcttagtggcgaagaagagaagaagcagaaatcaaatgtcatcgtttatcaacaaacctctaaattgatcgaatat
cttcttcaatcttcatctctctttcaaatctgcaaatgagagagctgtaagagttgcatcatctctatctctttctctctctctctt
gcgaaaacaccaaactttgaaatccccaaataagggtttaagcataaaccctacgagttcATGCCCAAATGCCCAATCCCAATCCG
TATCAGCTTCTTCAGCTACTTTCTCAAAGAAAGTCGAATCCTTTCGAGTAACCCAGTTAACTTCTCTATCCATCTTCGCTTCTCTTC
TTCTGTTTCGGTTTCTCCTAACCCGTCAATGGAAGTGGTGGAGAATCCATTGGAAGCTCCAATCTCGGAGAAGATGTTTAAATCAGC
TCCCAAAATGGGTTCTTTCAAGTTGGGTGATTCTACTTTATCTTCAATGATTGAGAGTTACGCCAATTCGGGTGATTTTGATTCGGT
AGAGAAGCTTTTGAGTCGAATTAGA(SALK_142675)TTAGAGAACAGAGTGATTATAGAGCGTAGTTTCATTGTTGTCTTTAGAG
CTTATGGGAAAGCACATTTGCCTGATAAAGCTGTAGACTTGTTTCATAGAATGGTTGATGAGTTTCGATGTAAACGTAGTGTTAAGT
CGTTTAACTCTGTGTTGAATGTGATTATAAACGAGGGTCTTTATCATCGAGGATTGGAGTTTTATGATTATGTTGTGAATTCCAACA
TGAACATGAACATTTCTCCTAATGGGTTGAGTTTTAATTTGGTCATTAAGGCTCTTTGTAAGTTAAGATTTGTGGATAGAGCTATTG
AGGTCTTCAGAGGAATGCCTGAGAGAAAGTGTTTGCCTGATGGTTATACGTATTGTACATTGATGGATGGGTTGTGTAAGGAGGAAA
GGATTGATGAGGCGGTTTTACTGTTGGATGAGATGCAGAGCGAGGGGTGTTCACCGAGTCCGGTTATATATAATGTGTTGATTGATG
GATTGTGCAAGAAAGGGGATTTGACACGGGTGACAAAGCTTGTGGATAATATGTTTCTTAAGGGATGTGTTCCTAATGAGGTTACTT
ATAATACGCTTATTCATGGTCTATGTCTTAAGGGTAAGTTAGATAAAGCTGTTAGTCTCTTGGAGCGGATGGTGTCTAGCAAATGTA
TCCCAAATGATGTCACATATGGAACGCTGATTAATGGGTTGGTTAAGCAAAGACGAGCAACGGATGCGGTTAGGTTGTTGAGCTCTA
TGGAAGAAAGAGGGTATCATTTGAATCAGCATATTTATTCGGTCCTTATAAGTGGGTTGTTCAAAGAGGGAAAGGCCGAGGAGGCTA
TGTCCTTGTGGAGGAAAATGGCAGAGAAAGGATGCAAGCCCAATATTGTTGTGTACAGTGTTCTTGTAGATGGTTTATGCCGCGAAG
GGAAACCAAATGAAGCTAAAGAGATCCTTAATAGAATGATTGCTAGTGGCTGCTTGCCCAATGCGTATACGTATAGCTCATTGATGA
AAGGATTCTTCAAAACTGGTCTTTGTGAAGAAGCGGTTCAAGTGTGGAAAGAGATGGATAAAACCGGATGTTCTCGAAATAAGTTTT
GTTATAGTGTTCTAATCGATGGTCTTTGTGGGGTTGGGAGAGTTAAGGAGGCTATGATGGTGTGGTCAAAGATGCTCACTATCGGGA
TCAAACCTGATACAGTAGCTTATAGCTCAATTATAAAAGGTCTTTGTGGTATTGGCTCGATGGATGCGGCCTTAAAACTTTACCATG
AGATGTTATGCCAAGAAGAACCCAAGTCACAACCTGATGTTGTTACTTATAATATACTTCTTGATGGCTTATGTATGCAAAAAGATA
TCTCTAGAGCAGTAGATCTCTTGAACTCTATGTTAGATAGAGGTTGTGATCCCGATGTTATTACGTGTAACACCTTTTTGAATACTT
TGAGCGAGAAGTCAAATTCTTGTGACAAAGGGAGGAGTTTTCTAGAAGAGCTTGTTGTAAGGCTCTTAAAGCGTCAGAGAGTATCAG
GCGCGTGTACGATTGTGGAAGTAATGCTGGGTAAGTATCTGGCACCGAAGACCTCCACTTGGGCGATGATTGTTCGAGAGATCTGTA
AACCAAAGAAGATCAATGCCGCCATTGATAAATGCTGGAGGAACCTGTGTACTTGAacactacagagcattttgctcagatcccgt
ttccaccttaagagtctccatgcgtgggaactttttgcgcttctggagttttggaataccagaaatgcgcgcgaaaaacaggtagtc
agataaactatttttgaaattctaactttttaaacatgttacttgatttatgcgcgagtaagaccacattagctaaggtttataagg
atatgaaaatcccattttgttgctattgaatcctccattgttaagatatgttctgtttcagacatgtttgatttgactatttgagat
ctttgtt…3′

**B.** **PCR screening of *msp1-1* progeny**

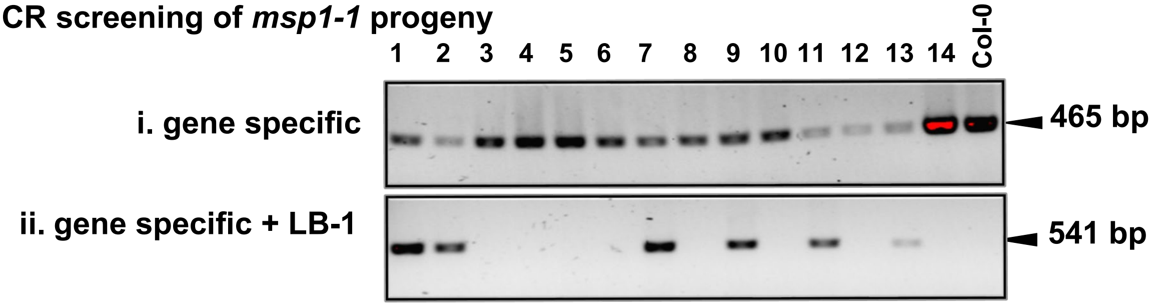


**C.** **Complementation of loss-of function *msp1* mutation**

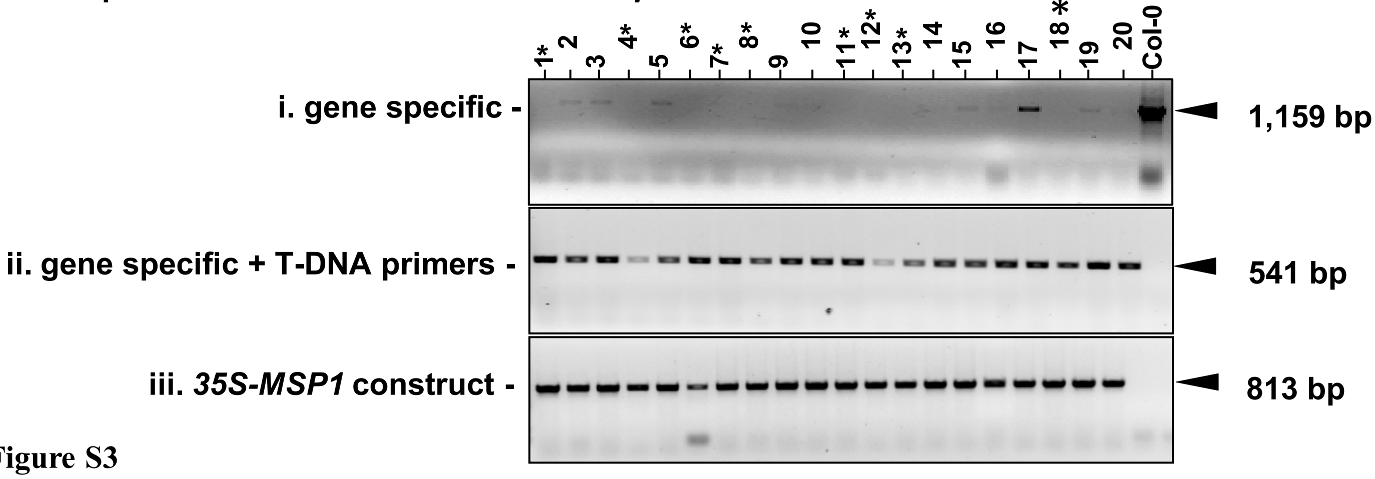


Figure S3

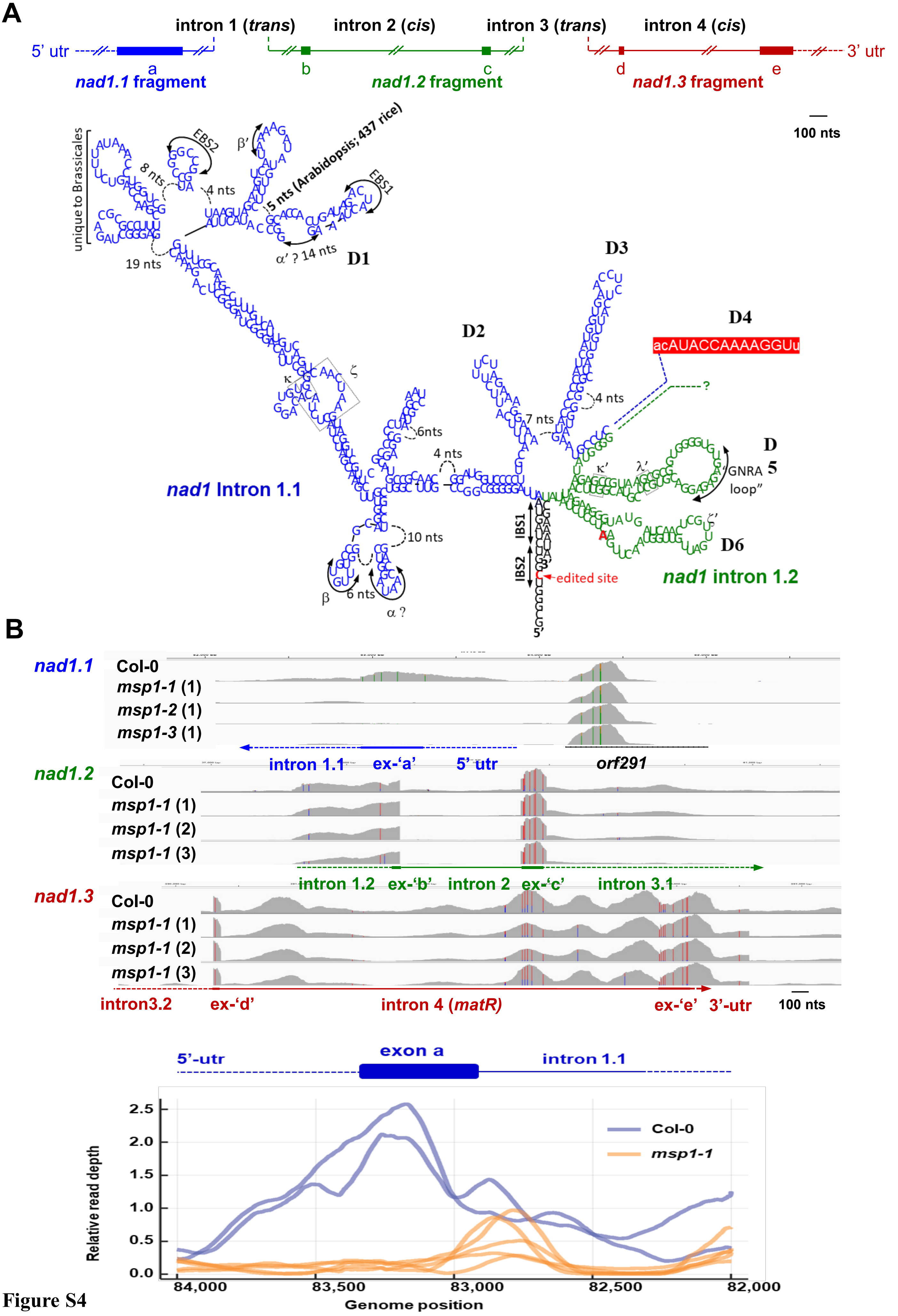
A
intron 1 (trans)
intron 2 (cis)
intron 3 (trans)
intron 4 (cis)
5' utr
3' utr
nad1.1 fragment
nad1.2 fragment
nad1.3 fragment
100 nts
unique to Brassicales
5 nts (Arabidopsis; 437 rice)
D1
D2
D3
D4
D5
D6
acAUACCAAAAGGUu
"GNRA loop"
edited site
IBS1
IBS2
nad1 Intron 1.1
nad1 intron 1.2
B
nad1.1
nad1.2
nad1.3
Col-0
msp1-1 (1)
msp1-2 (1)
msp1-3 (1)
msp1-1 (2)
msp1-1 (3)
intron 1.1
ex-'a'
5' utr
orf291
intron 1.2
ex-'b'
intron 2
ex-'c'
intron 3.1
intron3.2
ex-'d'
intron 4 (matR)
ex-'e'
3'-utr
100 nts
5'-utr
exon a
intron 1.1
Relative read depth
Genome position
Col-0
msp1-1

Figure S4

#### A. predicted MSP1 RNA recognition site

| PPR domain | 5th / last AAs | A | C | G | U | |
|---|---|---|---|---|---|---|
| GLSF**N**LVIKALCKLRFVDRAIEVFRG MPERKCLP**D** | **N / D** | -0.527074216 | 0.661761035 | 0.554154587 | **0.828282956** | **U>C>G** |
| GYTY**C**TLMDGLCKEERIDEAVLLLDE MQSEGCSP**S** | **C / S** | **0.150493483** | -1 | -0.383947494 | -1 | **A** |
| PVIY**N**VLIDGLCKKGDLTRVTKLVDN MFLKGCVP**N** | **N / N** | -0.444301501 | **0.771492317** | -0.069522593 | **0.618918582** | **C>U** |
| EVTY**N**TLIHGLCLKGKLDKAVSLLER MVSSKCIP**N** | **N / N** | -0.444301501 | **0.771492317** | -0.069522593 | **0.618918582** | **C>U** |
| DVTY**G**TLINGLVKQRRATDAVRLLSS MEERGYHL**N** | **G / N** | **0.416872982** | -0.299148456 | -0.195618907 | -0.512394419 | **A** |
| QHIY**S**VLISGLFKEGKAEEAMSLWRK MAEKGCKP**N** | **S / N** | **0.723559179** | -0.231353898 | -0.195618907 | -0.223389259 | **A** |
| IVVY**S**VLVDGLCREGKPNEAKEILNR MIASGCLP**N** | **S / N** | **0.723559179** | -0.231353898 | -0.195618907 | -0.223389259 | **A** |
| AYTY**S**SLMKGFFKTGLCEEAVQVWK EMDKTGCSR**N** | **S / N** | **0.723559179** | -0.231353898 | -0.195618907 | -0.223389259 | **A** |
| KFCY**S**VLIDGLCGVGRVKEAMMVWS KMLTIGIKP**D** | **S / D** | **0.167028452** | -0.490904827 | **0.946060803** | 0.065409631 | **G>>A>U** |
| TVAY**S**SIIKGLCGIGSMDAALKLYHE MLCQEEPKSQP**D** | **S / D** | **0.167028452** | -0.490904827 | **0.946060803** | 0.065409631 | **G>>A>U** |
| VVTY**N**ILLDGLCMQKDISRAVDLLNS MLDRGCDP**D** | **N / D** | -0.527074216 | 0.661761035 | 0.554154587 | **0.828282956** | **U>C>G** |

#### B. Distribution of MSP1-mtRNA target scores

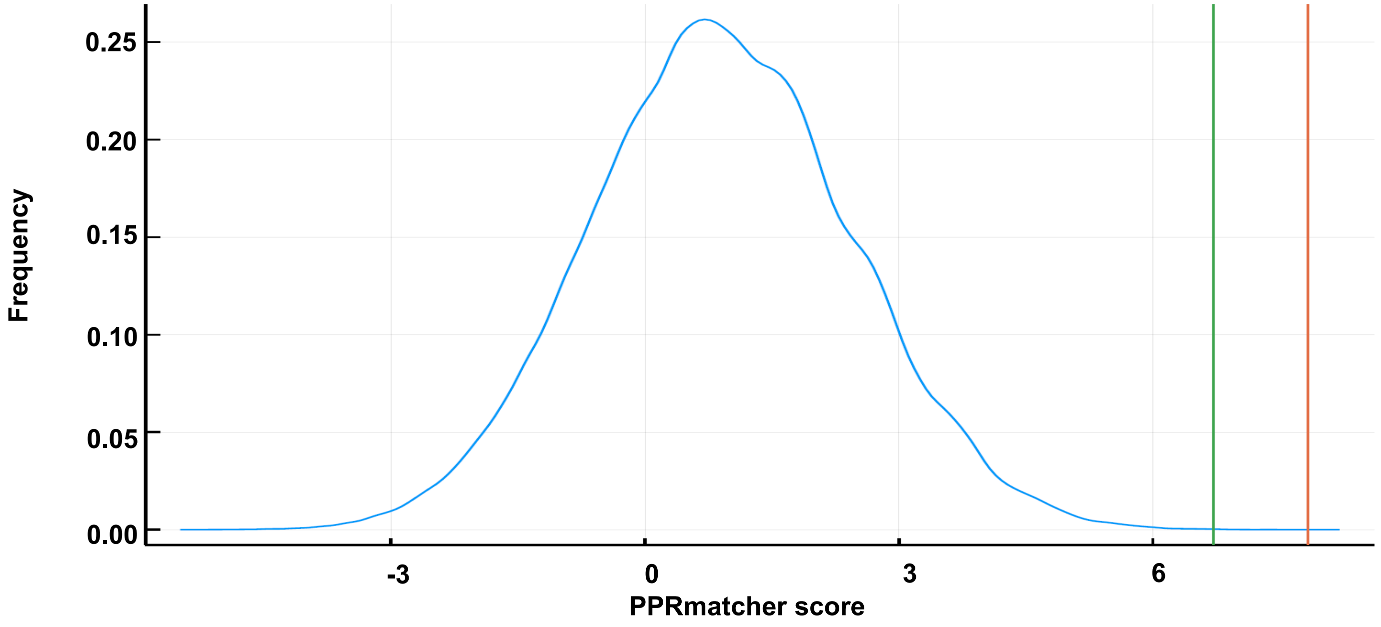


Figure S5

A

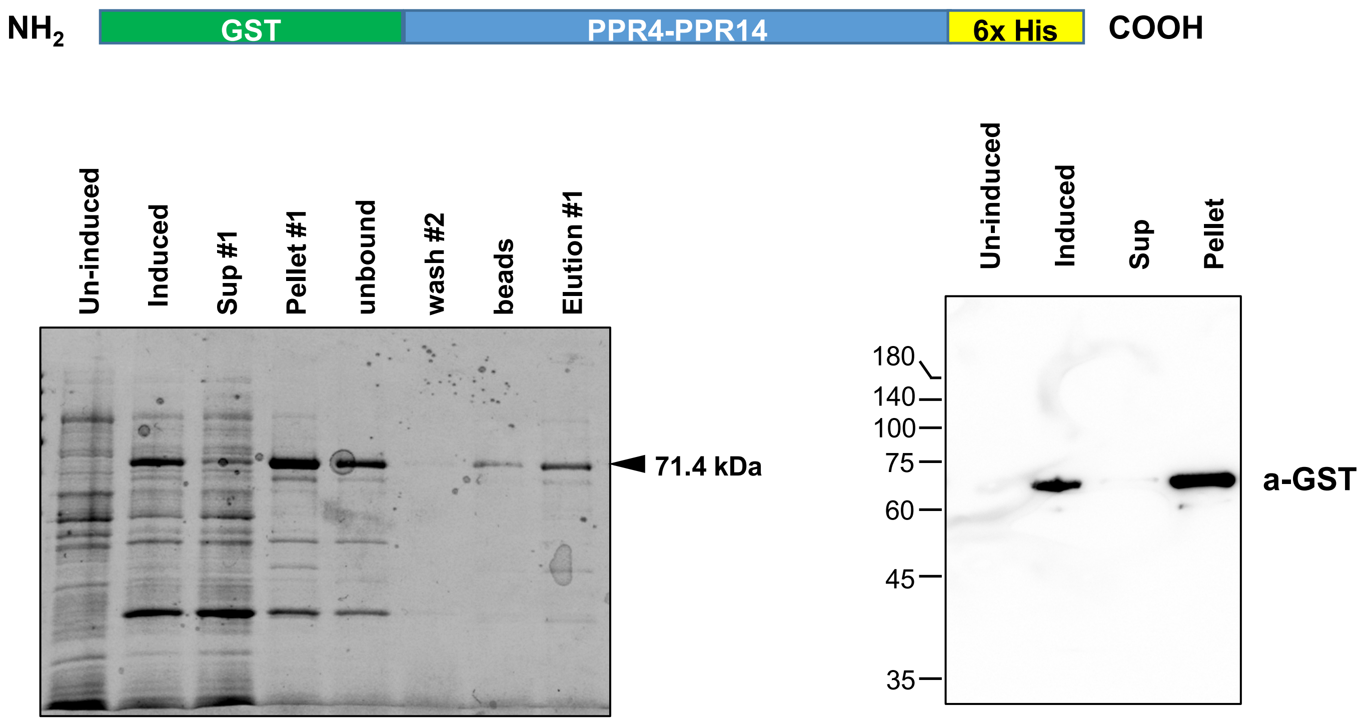
NH2
GST
PPR4-PPR14
6x His
COOH
Un-induced
Induced
Sup #1
Pellet #1
unbound
wash #2
beads
Elution #1
71.4 kDa
Un-induced
Induced
Sup
Pellet
180
140
100
75
60
45
35
a-GST

B

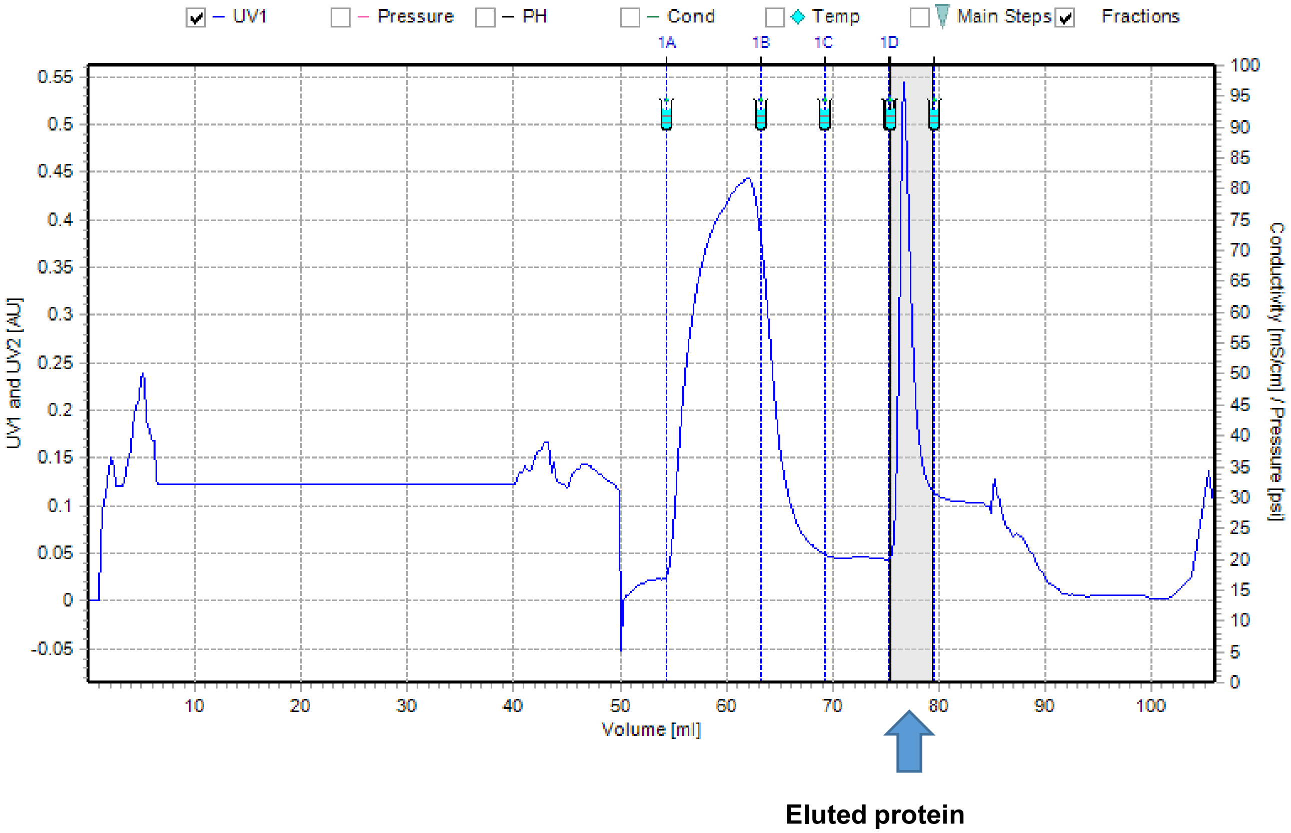
UV1
Pressure
PH
Cond
Temp
Main Steps
Fractions
1A
1B
1C
1D
UV1 and UV2 [AU]
Conductivity [mS/cm] / Pressure [psi]
Volume [ml]
Eluted protein

Figure S6

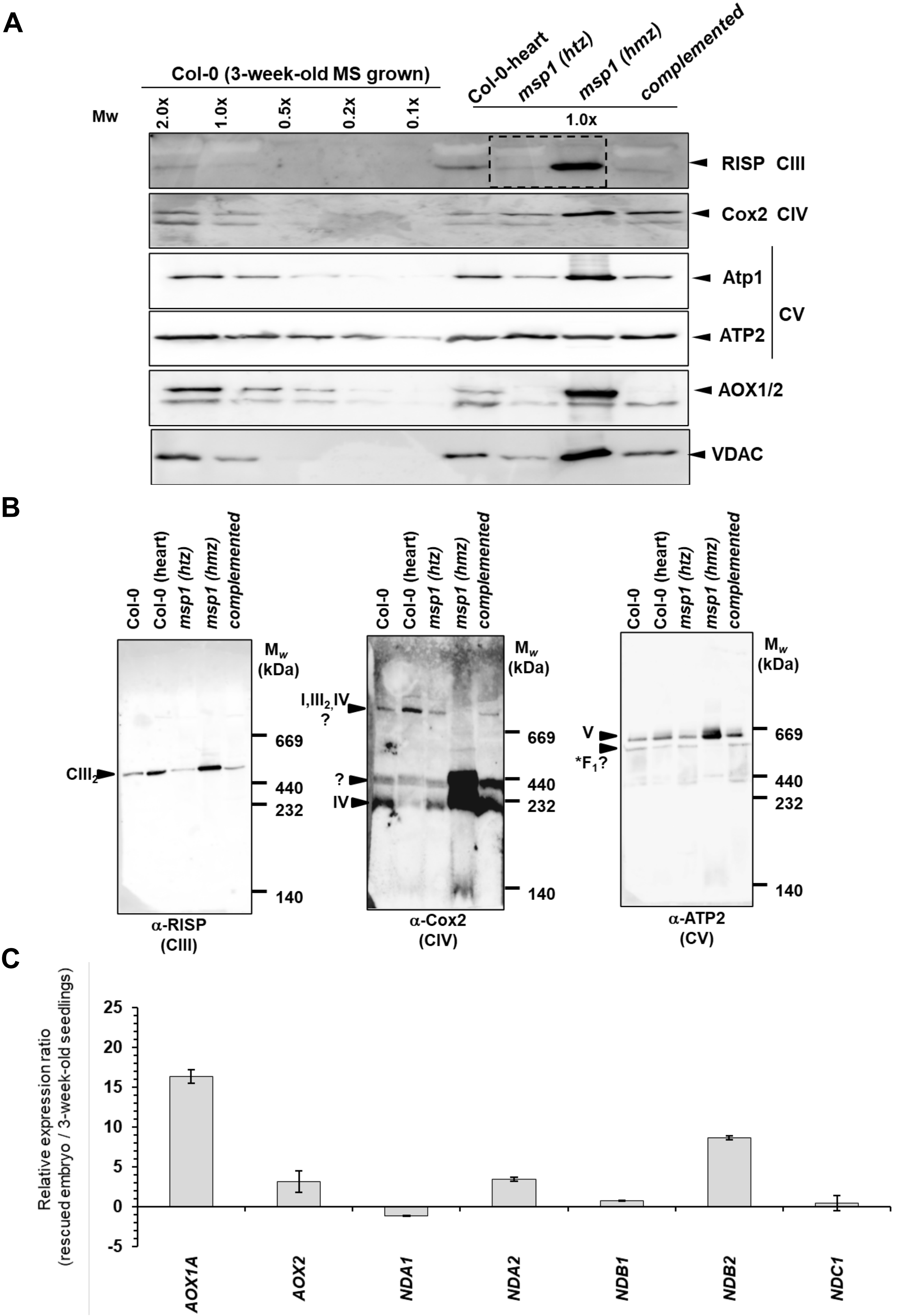

A
Col-0 (3-week-old MS grown)
Col-0-heart
msp1 (htz)
msp1 (hmz)
complemented
Mw
2.0x
1.0x
0.5x
0.2x
0.1x
1.0x
RISP CIII
Cox2 CIV
Atp1
ATP2
CV
AOX1/2
VDAC
B
Col-0
Col-0 (heart)
msp1 (htz)
msp1 (hmz)
complemented
Mw (kDa)
669
440
232
140
CIII2
α-RISP (CIII)
I,III2,IV ?
?
IV
α-Cox2 (CIV)
V
*F1?
α-ATP2 (CV)
C
Relative expression ratio (rescued embryo / 3-week-old seedlings)
25
20
15
10
5
0
-5
AOX1A
AOX2
NDA1
NDA2
NDB1
NDB2
NDC1


**Figure S7**

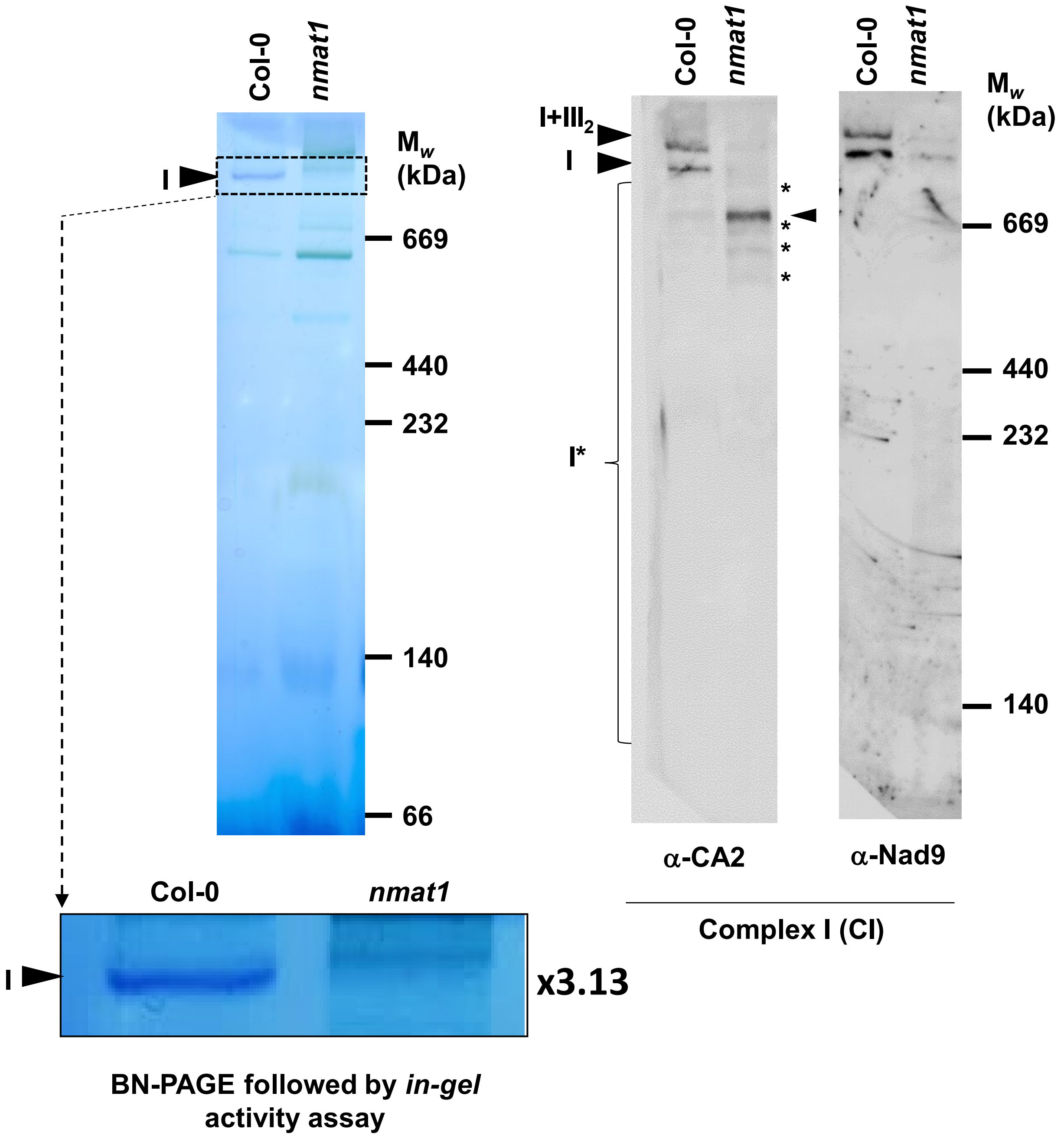

Col-0
nmat1
I
Mw (kDa)
669
440
232
140
66
Col-0
nmat1
I
x3.13
BN-PAGE followed by in-gel activity assay
Col-0
nmat1
Col-0
nmat1
I+III2
I
I*
*
*
*
*
669
440
232
140
α-CA2
α-Nad9
Complex I (CI)


**Figure S8**

```
                *        20         *        40         *        60         *        80         *       100
A.t.:  GCCTAGCTTTGGTAGATTTCCCCCAACGCAATTCACAG----AAATTCCACGAATCCC---T-TGGTGGATAAGTCCGACGACTCAGCAGCAGTGCGGAATTGAGTTT : 100
S.l.:  GCCTAGCTTTGGTAGATTTCCCCCAACGCAATCCACA----GAAATTCCACGAATCCC---TT-GGTGGATAAGTCCGACGACTCAGCAGCAGTGCGGAATTGAGTTT : 100
N.t.:  GCCTAGCTTTGGTAGATTTCCCCCAACGCAATCCACA----GAAATTCCACGAATCCC---TT-GGTGGATAAGTCCGACGACTCAGCAGCAGTGCGGAATTGAGTTT : 100
T.a.:  GCCTAGCTTTGGTAGATTTCCCCCAACGCAATTCAAAAACGGAAATTCCACGAATCCC---TGAGGTGGATAAGTCCGACGACTCAGCAGCAGTGCGGAATTGAGTTT : 105
O.s.:  GCCTAGCTTTGGTAGATTTCCCCCAACGCAATTCAAAAACGGAAATTCCACGAATCCT---TGAGGTGGATAAGTCCGACGACTCAGCAGCAGTGCGGAATTGAGTTT : 105
G.ma.: GCCTAGCTTTGGTAGATTTCCCCCAACGCAATCCATA----TAAATTCCACGAATCCC---TT-GGTGGATAAGTCCGACGACTCAGCAGCAGTGCGGAATGGAGTTT : 100
Z.m.:  GCCTAGCTTTGGTAGATTTCCCCCAACGCAATTCAAAAACGGAAATTCCACGAATCCC---TGAGGTGGATAAGCCCGACGACTCAGCAGCAGTGCGGAATTGAGTTT : 105
G.mo.: GCCTAGCTTTGGTAGATTTCCCCCAACGCAATCCGCA----GAAATTCCACGAATCCC---TT-GGTGGATAAGTCCGACGACTCAGCAGCAGTGCGGAATTGAGTTT : 100
G.b.:  GCCTAGCTTTGGTAGATTTTCCCCGGCGCAATCCACAAACTAGAATTCCACGAATCCCATCCTTGGTGGATAAGTCCGACGACTCAGCAGCAGTGCGGAATGGAGTTC : 108
C.t.:  GCCTAGCTCCGGCAGATCTTCCCCAACGCAATCCACAAACTAGAATTCCGCGAATCCCATCCTTGGTGGATAAGTCCGACGACTCAGCAGCAGTGCGGAATGGAGTTT : 108
C.h.:  GCCTAGCTCCGGCAGATCTTCCCCAACGCAATCCACAAACTAGAATCCCGT----CCCATCCTTGGTGGATAAGTCCGACGACTCAGCAGCAGTGCGGAATGGGGTTT : 104
Z.i.:  GCCTAGCTCCGGCAGATCTTCCCCAACGCAATCCACAAACTAGAATCCCGT----CCCATCCTTGGTGGATAAGTCCGACGACTCAGCAGCAGTGCGGAATGGGGTTT : 104
       GCCTAGCT  GG AGAT T CCCCaaCGCAAT Ca A      AATtCC cgaatCCc      GGTGGATAAGtCCGACGACTCAGCAGCAGTGCGGAAT GaGTTt

        *       120         *       140         *       160         *       180
A.t.:  CTCGTCTGGTGGATCT---GATCTATA--G--TT----------ATGGGGCAATACATACCAAAAGGTTCGA-AGAA-AG : 161
S.l.:  CTTGTCTGGTGGATCT---GATCTATA--G--TT----------AGGGGGCAATACATACCAAAAGGTTCGA-AGA---A : 159
N.t.:  CTTGTCTGGTGGATCT---GATCTATA--G--TT----------AGGGGGCAATACATACCAAAAGGTTCGA-AGA---A : 159
T.a.:  CTGGTCCGGTGGATCT---GATCTATA--G--TT----------AGGGGGCAATACATACCAAAAGGTTCGA-AGA---A : 164
O.s.:  CTGGTCCGGTGGATCT---GATCTATA--G--TT----------AGGGGGCAATACATACCAAAAGGTTCGA-AGA---A : 164
G.ma.: CTCGTCTGGTGGATCT---GATCT----T----T----------AGGGGGCAATACATACCAAAAGGTGCTTTAG----- : 154
Z.m.:  CTGGTCCGGTGGATCT---GATCTATTCTA--TT----------CTGGGGCAATACATACCAAAAGGTTCGA-AAGA--A : 167
G.mo.: CTCGTCCGGTGGATCT---GATCTATA--G--TT----------AGGGGGCAATACATACCAAAAGGTTCGA-AGAAGGA : 162
G.b.:  CTCGTCCGGTGGATCTTCGGATCTATA--GTTCCGACCGAGGAAAGGGGGCGATACATACTAAAAGGTTCGA-AGA---A : 182
C.t.:  CTCGTCCGGTGGATCTTCGGATCCATA--GTTCCGG----GGAAAGGGGGCGATACATACCAAAAGGTTCGA-AGAAGGA : 181
C.h.:  CTCGTCCGGTGGATCTTCGGATCCATA--GTTCCGG----GGAAAGGGGGCGATACATACCAAAAGGTTCGA-AGAAGGA : 177
Z.i.:  CTCGTCCGGTGGATCTTCGGATCCATA--GTTCCGG----GGAAAGGGGGCGATACATACCAAAAGGTTCGA-AGA---A : 174
       CT GTC GGTGGATCT   GATC ata  g              agGGGGC ATACATACcAAAAGGTtCga Aga   a
```

**MSP1 RNA binding site**

### Figure S9

A

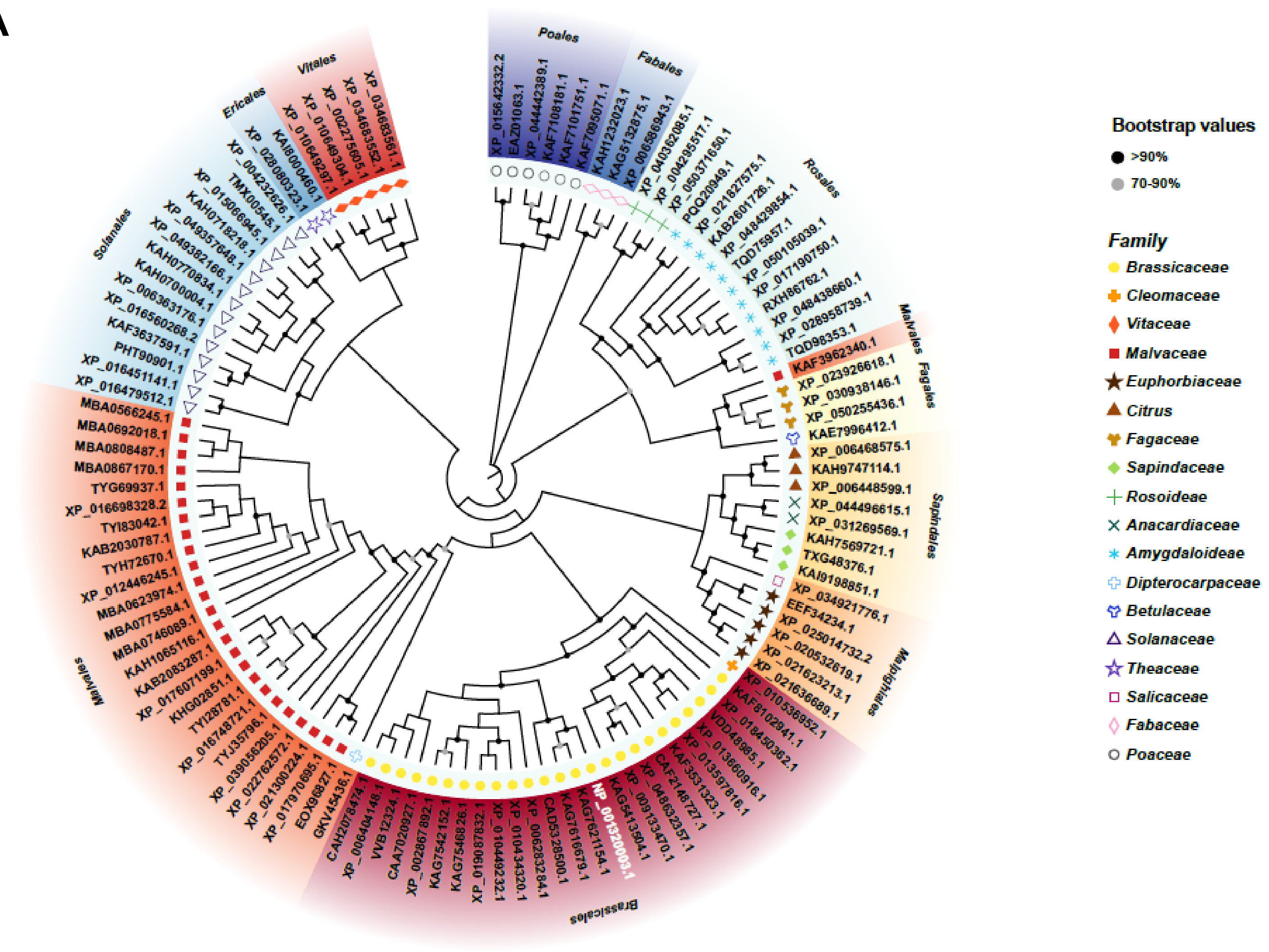


B

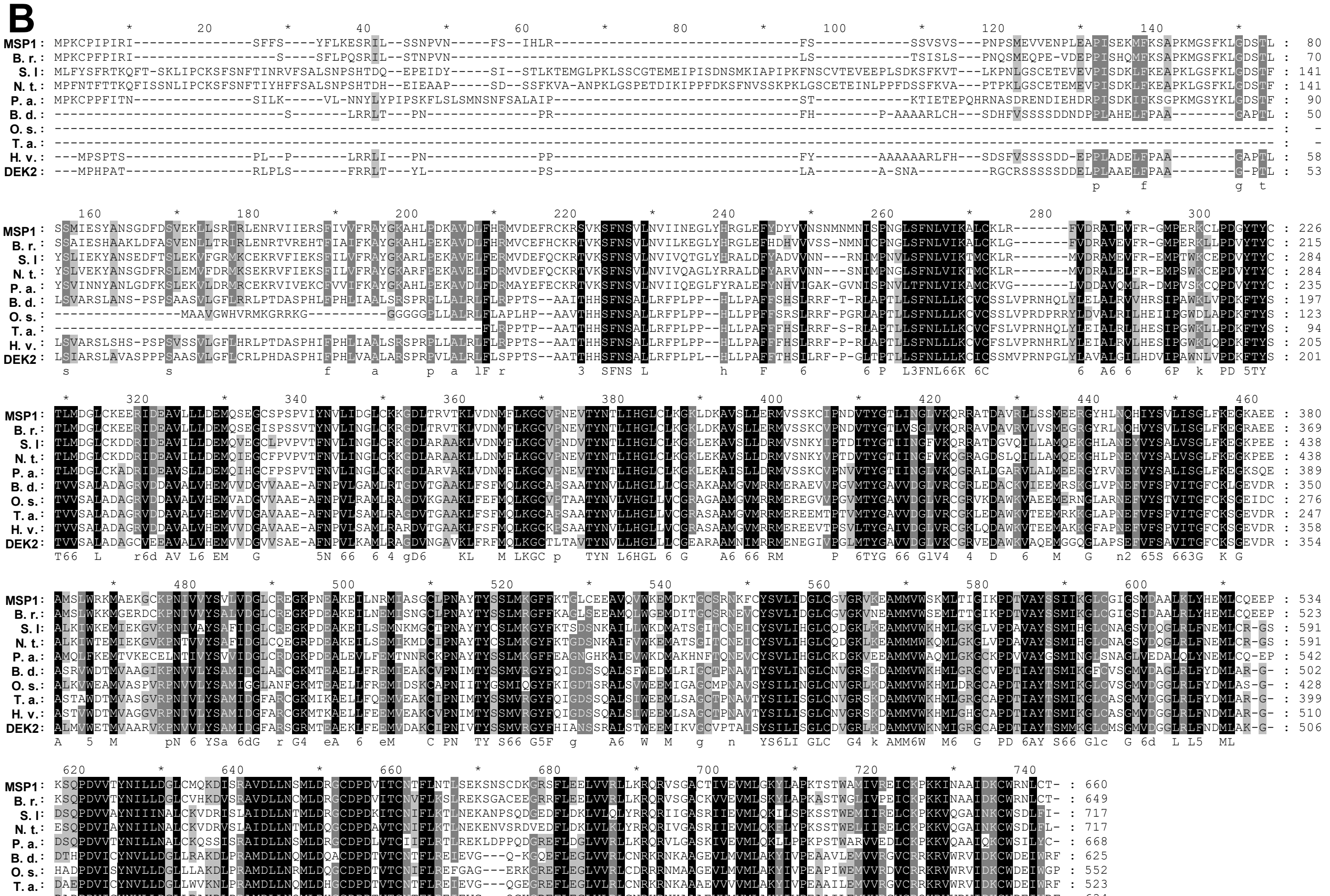


Figure S10

#### Splicing efficiencies in *ca2*

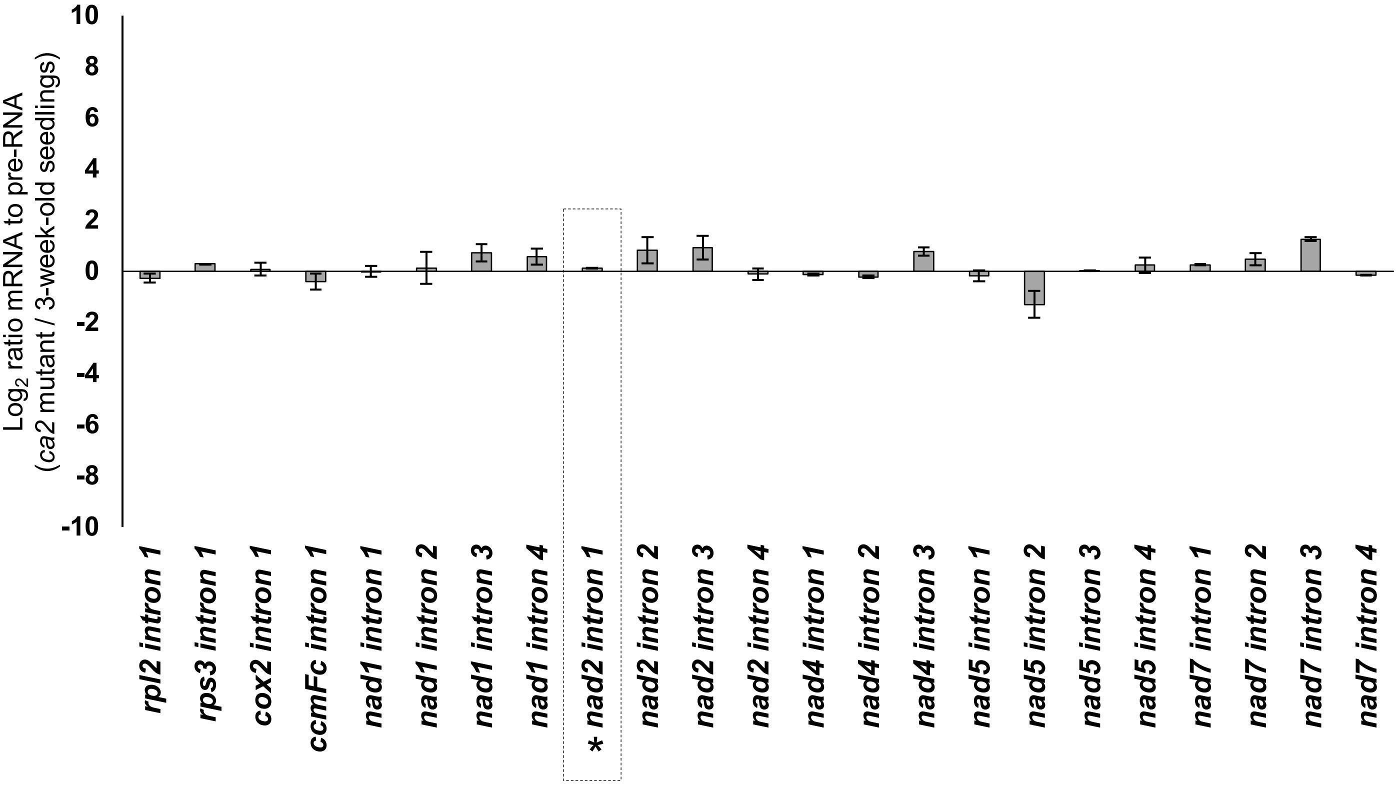


**Figure S11**

**Supplementary Table S1.**

a. Lists of oligonucleotides used for the analysis of the splicing profiles of wild-type and *msp1-1* mutant plants by RT-qPCR experiments.

| Gene | Forward primer | Reverse primer |
|---|---|---|
| *rpl2* | CCGAAGACGGATCAAGGTAA | CGCAATTCATCACCATTTTG |
| *rpl2* intron exon2 | TTAGGAAGAGCCGTACGAGG | CGCAATTCATCACCATTTTG |
| *rps3* | AGCCGAAGGTGAGTCTCGTA | CCGATTTCGGTAAGACTTGG |
| *rps3* intron1 exon2 | AGCCGAAGGTGAGTCTCGTA | TCTACGGCGGGGTCACTAT |
| *cox2* | TGGGGGATTAATTGATTGGA | TGATGCTGTACCTGGTCGTT |
| *cox2* intron1 exon2 | TGGGGGATTAATTGATTGGA | AGCAGTACGAGCTGAAAGGC |
| *ccmFc* | GTGGGTCCATGTAAATGATCG | CACATGGAGGAGTGTGCATC |
| *ccmFc* intron1 exon1 | CCCGGATCGAATCAGAGTT | CACATGGAGGAGTGTGCATC |
| *nad1* exon1-2 | GACCAATAGATACTTCATAAGAGACCA | TTGCCATATCTTCGCTAGGTG |
| *nad1* intron1 exon2 | GACCAATAGATACTTCATAAGAGACCA | CGTGCTCGTACGGTTCATAG |
| *nad1* exon2-3 | ATTCAGCTTCCGCTTCTGG | TCTGCAGCTCAAATGGTCTC |
| *nad1* intron2 exon2 | GGTTGGGTTAGGGGAACATC | TCTGCAGCTCAAATGGTCTC |
| *nad1* exon3-4 | AAAAGAGCAGACCCCATTGA | TCCGTTTGATCTCCCAGAAG |
| *nad1* intron3 exon4 | AAAAGAGCAGACCCCATTGA | GGGAGCTGTATGAGCGGTAA |
| *nad1* exon4-5 | AGCCCGGGATCTTCTTGA | TCTTCAATGGGGTCTGCTC |
| *nad1* intron4 exon5 | AGCCCGGGATCTTCTTGA | ACGGAGCTGCATCCCTACT |
| *nad2* exon1-2 | GCGAGCAGAAGCAAGGTTAT | GGATCCTCCCACACATGTTC |
| *nad2* intron1 exon2 | GCGAGCAGAAGCAAGGTTAT | CCCATTCCTAACCAGTGGAG |
| *nad2* exon2-3 | AAAGGAACTGCAGTGATCTTGA | AATATTTGATCTTAGGTGCATTTTC |
| *nad2* intron2 exon2 | CCCGATCCGATAGTTTACAA | AATATTTGATCTTAGGTGCATTTTC |
| *nad2* exon3-4 | GCGCAATAGAAAGGAATGCT | CTATGGGTCTACTGGAGCTACCC |
| *nad2* intron3 exon4 | GCGCAATAGAAAGGAATGCT | GGCGAATTTCAAACTTGTGG |
| *nad2* exon4-5 | CAAAGGAGAGGGGTATAGCAA | TATTTGTTCTTCGCCGCTTT |
| *nad2* intron4exon4 | CTTATTCGTGGCAACCTTCC | TATTTGTTCTTCGCCGCTTT |
| *nad4* exon1-2 | ATTCTATGTTTTTCCCGAAAGC | GAAAAACTGATATGCTGCCTTG |
| *nad4* intron1 exon2 | CCGTATGATGCGGAAGTCTC | GAAAAACTGATATGCTGCCTTG |
| *nad4* exon2-3 | AATACCCATGTTTCCCGAAG | TGCTACCTCCAATTCCCTGT |
| *nad4* intron2 exon3 | GCGGAACGACCAGAAAAATA | TGCTACCTCCAATTCCCTGT |
| *nad4* exon3-4 | TTCCTCCATAAATTCTCCGATT | TGAAATTTGCCATGTTGCAC |
| *nad4* intron3 exon4 | TCTAGCTTGGTTCGGAGAGC | TGAAATTTGCCATGTTGCAC |
| *nad5* exon1-2 | TGGACCAAGCTACTTATGGATG | CCATGGATCTCATCGGAAAT |
| *nad5* intron1 exon2 | TGGACCAAGCTACTTATGGATG | TTCGCAAATAGGTCCGACT |
| *nad5* exon2-3 | TACCTAAACCAATCATCATATC | CTGGCTCTCGGGAGTCTCTT |
| nad5 intron2-exon2 | GTACGATCGTGTCGGGTGA | CTGGCTCTCGGGAGTCTCTT |
| *nad5* exon3-4 | AACTCGGATTCGGCAAGAA | GATATGATGATTGGTTTAGGTA |
| nad5 intron3-exon4 | AACTCGGATTCGGCAAGAA | GCCGTGTAATAGGCGACCA |
| *nad5* exon4-5 | AACATTGCAAAGGCATAATGA | GTTCCTGCGTTTCGGATATG |
| *nad5* intron4 exon5 | AACATTGCAAAGGCATAATGA | CCTGTAAACCCCCATGATGT |
| *nad7* exon1-2 | ACCTCAACATCCTGCTGCTC | AAGGTAAAGCTTGAAGATAAGTTTTGT |
| *nad7* intron1 exon2 | ACGGTTTTTAGGGGGATCTG | AAGGTAAAGCTTGAAGATAAGTTTTGT |
| *nad7* exon2-3 | GAGGGACTGAGAAATTAATAGAGTACA | TGGTACCTCGCAATTCAAAA |

| | | |
|---|---|---|
| *nad7* intron2 exon3 | AGTGGGAGAGCCGTGTTATG | TGGTACCTCGCAATTCAAAA |
| *nad7* exon3-4 | ACTGTCACTGCACAGCAAGC | CATTGCACAATGATCCGAAG |
| *nad7* intron3 exon4 | TAAAGTGAAGTGGTGGGCCT | CATTGCACAATGATCCGAAG |
| *nad7* exon4-5 | GATCAAAGCCGATGATCGTAA | AGGTGCTTCAACTGCGGTAT |
| *nad7* intron4 exon5 | CGGCCAAATGACTACAGGAT | AGGTGCTTCAACTGCGGTAT |

b. Lists of oligonucleotides used for the analysis of the mRNA profiles of wild-type and *msp1-1* mutant plants by RT-qPCR experiments.

| gene target | oligo name | sequence (5'-to-3') |
|---|---|---|
| *AOX1A* | *aox1aF* | AGCATCATGTTCCAACGACGTTTC |
| | *aox1aR* | GCTCGACATCCATATCTCCTCTGG |
| *AOX1B* | *aox1bF* | GGACCGTGAAATCTCTTCGATGGC |
| | *aox1bR* | TCTAGCATCATTGCTCTGCATCCG |
| *AOX1C* | *aox1cF* | TCTTCCAGAGGAGGTATGGTTGCC |
| | *aox1cR* | AGTGCATAAGCATCCCTCCAACC |
| *AOX1D* | *aox1dF* | TTTGCTCGAAGAGGCTGAGAACG |
| | *aox1dR* | CTCGTTCGTACCATTTGGGTTGTG |
| *AOX2* | *aox2F* | ACGGTGATTCGTGCTGATGAAGC |
| | *aox2R* | TCCTTGATTGCGAATGTCAGAAGC |
| *NDA1* | *nda1F* | GTATCCAACCGGCGATTTCACG |
| | *nda1R* | AGTTACAGTCTCACAATGCACCTC |
| *NDA2* | *nda2F* | TGGTGTTGGTCCTTCTCCTTTCG |
| | *nda2R* | TCCATTCGTCAATGCCAATCCTTC |
| *NDB1* | *ndb1F* | TAACACATTTGGCACTCCTGGTG |
| | *ndb1R* | CTCTGTGCATCCTCTACTTCCTTG |
| *atp1* | *atp1F* | TCACTTCGACACGTCTTTGC |
| | *atp1R* | GGAATGGCCTTGAATCTTGA |
| *atp6* | *atp6-1F* | TCTTTTGCGAGTCAATGCAC |
| | *atp6-1R* | TCTCGCGTATCTCACATTGC |
| *atp8* | *atp8F* | CCGTCGACTTATTGGGAAAA |
| | *atp8R* | TTCCTTGGCCATGTACAACA |
| *atp9* | *atp9F* | CATTCCCTCTGACGTCGAAT |
| | *atp9R* | TCGTCGATTCTTACCCTCGT |
| *atp4* | *atp4F* | GGATCAGCTTGCGAATTTGT |
| | *atp4R* | GCAAATTGCTTCCCCACTAA |
| *ccmb* | *ccmBF* | TCTTGGAATCACATCCAGCA |
| | *ccmBR* | CGAGACCGAAATTGGAAAAA |
| *ccmc* | *ccmCF* | AGCTACGCGCAAATTCTCAT |
| | *ccmCR* | GCCGTGGCGATATAAACAAT |
| *ccmfc* | *ccmFcF* | CACATGGAGGAGTGTGCATC |
| | *ccmFcR* | GTGGGTCCATGTAAATGATCG |
| *ccmfn-1* | *ccmFN1F* | AGCTCTTGGCATTGCTTTGT |
| | *ccmFN1R* | AGTGCCACAATCCCATTCAT |
| *ccmfn-2* | *ccmFN2F* | CGTGTCGTTCGTAATGGAAA |
| | *ccmFN2R* | TGATAAGCCCACCAACTTCC |

| | | |
|---|---|---|
| *cob* | *cobF* | TGCCGGAATGGTATTTCCTA |
| | *cobR* | GCCAAAAGCAACCAAAACAT |
| *cox1* | *cox1F* | GTAGCTGCGGTGAAGTAGGC |
| | *cox1R* | CTGCCTGGATTCGGTATCAT |
| *cox2* | *cox2F* | TGATGCTGTACCTGGTCGTT |
| | *cox2R* | TGGGGGATTAATTGATTGGA |
| *cox3* | *cox3F* | CCGTAACTTGGGCTCATCAT |
| | *cox3R* | AAACCATGAAAGCCTGTTGC |
| *mttb* | *mttBF* | GGGGTCTTTCTTTGGAAACC |
| | *mttBR* | TCTCCCTCATTCCACTCGTC |
| *nad1 exons a-b* | *nad1 1-2F* | GACCAATAGATACTTCATAAGAGACCA |
| | *nad1 1-2R* | TTGCCATATCTTCGCTAGGTG |
| *nad1 exons b-c* | *nad1 2-3F* | ATTCAGCTTCCGCTTCTGG |
| | *nad1 2-3R* | TCTGCAGCTCAAATGGTCTC |
| *nad1 exons c-d* | *nad1 3-4F* | AAAAGAGCAGACCCCATTGA |
| | *nad1 3-4R* | TCCGTTTGATCTCCCAGAAG |
| *nad1 exons d-e* | *nad1 4-5F* | AGCCCGGGATCTTCTTGA |
| | *nad1 4-5R* | TCTTCAATGGGGTCTGCTC |
| *nad2 exons a-b* | *nad2 exons a-bF* | GCGAGCAGAAGCAAGGTTAT |
| | *nad2 exons a-bR* | GGATCCTCCCACACATGTTC |
| *nad2 exons b-c* | *nad2 exons b-cF* | AAAGGAACTGCAGTGATCTTGA |
| | *nad2 exons b-cR* | AATATTTGATCTTAGGTGCATTTTC |
| *nad2 exons c-d* | *nad2 exons c-dF* | GCGCAATAGAAAGGAATGCT |
| | *nad2 exons c-dR* | CTATGGGTCTACTGGAGCTACCC |
| *nad2 exons d-e* | *nad2 exons d-eF* | CAAAGGAGAGGGGTATAGCAA |
| | *nad2 exons d-eR* | TATTTGTTCTTCGCCGCTTT |
| *nad3* | *nad3F* | CGAATGTGGTTTCGATCCTT |
| | *nad3R* | GCACCCCTTTTCCATTCATA |
| *nad4 exons a-b* | *nad4 exons a-bF* | ATTCTATGTTTTTCCCGAAAGC |
| | *nad4 exons a-bR* | GAAAAACTGATATGCTGCCTTG |
| *nad4 exons b-c* | *nad4 exons b-cF* | AATACCCATGTTTCCCGAAG |
| | *nad4 exons b-cR* | TGCTACCTCCAATTCCCTGT |
| *nad4 exons c-d* | *nad4 exons c-dF* | TTCCTCCATAAATTCTCCGATT |
| | *nad4 exons c-dR* | TGAAATTTGCCATGTTGCAC |
| *nad4L* | *nad4L-F* | GGGGAATCCTCCTTAATAGACG |
| | *nad4L-R* | AACGAAAATGGCTAACCCAATA |
| *nad5 exons a-b* | *nad5 exons a-bF* | TGGACCAAGCTACTTATGGATG |
| | *nad5 exons a-bR* | CCATGGATCTCATCGGAAAT |
| *nad5 exons b-c* | *nad5 exons b-cF* | TACCTAAACCAATCATCATATC |
| | *nad5 exons b-cR* | CTGGCTCTCGGGAGTCTCTT |
| *nad5 exons c-d* | *nad5 exons c-dF* | AACTCGGATTCGGCAAGAA |
| | *nad5 exons c-dR* | GATATGATGATTGGTTTAGGTA |
| *nad5 exons d-e* | *nad5 exons d-eF* | AACATTGCAAAGGCATAATGA |
| | *nad5 exons d-eR* | GTTCCTGCGTTTCGGATATG |
| *nad6* | *nad6F* | TATGCCGGAAAGGTACGAAG |
| | *nad6R* | GTGAGTGGGTCAGTCGTCCT |
| *nad7 exons a-b* | *nad7 exons a-bF* | ACCTCAACATCCTGCTGCTC |

| | | |
|---|---|---|
| | *nad7 exons a-bR* | AAGGTAAAGCTTGAAGATAAGTTTTGT |
| *nad7 exons b-c* | *nad7 exons b-cF* | GAGGGACTGAGAAATTAATAGAGTACA |
| | *nad7 exons b-cR* | TGGTACCTCGCAATTCAAAA |
| *nad7 exons c-d* | *nad7 exons c-dF* | ACTGTCACTGCACAGCAAGC |
| | *nad7 exons c-dR* | CATTGCACAATGATCCGAAG |
| *nad7 exons d-e* | *nad7 exons d-eF* | GATCAAAGCCGATGATCGTAA |
| | *nad7 exons d-eR* | AGGTGCTTCAACTGCGGTAT |
| *nad9* | *nad9F* | GGATGACCCTCGAAACCATA |
| | *nad9R* | CACGCATTCGTGTACAAACC |
| *rpl2* | *rpl2F* | CCGAAGACGGATCAAGGTAA |
| | *rpl2R* | CGCAATTCATCACCATTTTG |
| *rpl5* | *rpl5F* | AAGGGGTTCGACAGGAAAGT |
| | *rpl5R* | CGTATTTCGACCGGAAAATC |
| *rpl16* | *rpl16F* | GAGCATTTGCCAAACTCACA |
| | *rpl16R* | CGGACACTTTCATCGTGCTA |
| *rps3* | *rps3F* | CCGATTTCGGTAAGACTTGG |
| | *rps3R* | AGCCGAAGGTGAGTCTCGTA |
| *rps4* | *rps4F* | ACCCATCACAGAGATGCACA |
| | *rps4R* | TCACACAAACCCTTCGATGA |
| *rps7* | *rps7F* | CTCGAACTGAACGCGATGTA |
| | *rps7R* | AAGCTGCTTCAAGGATCCAA |
| *rps12* | *rps12F* | AGCCAAAGTACGGTTGAGCA |
| | *rps12R* | TTTGGGTTTTTCTGCACCAT |
| *matR* | *matR-F* | AATTTTTGCGAGAGCTGGAA |
| | *matR-R* | TTGAACCCCGTCCTGTAGAC |
| *rrn18* | *rrn18F* | CGTCACCTGGGTCAAAAACT |
| | *rrn18R* | GCTTGAAAACCGAAGTGAGC |
| *rrn26* | *rrn26F* | GACGAGACTTTCGCCTTTTG |
| | *rrn26R* | CTTGGAGCGAATTGGATGAT |
| *rrn5* | *rrn5F* | CCGACCTCGATATGTGGAATCGTC |
| | *rrn5R* | TGGACCATGTCTCCCGAACAATC |
| *18S rRNA* (nuclear) | *18S nucl-F* | AAACGGCTACCACATCCAAG |
| | *18S nucl-R* | ACTCGAAAGAGCCCGGTATT |
| *actin2* (At3g18780, nuclear) | *actin2-F* | GGTAACATTGTGCTCAGTGGTGG |
| | *actin2-R* | AACGACCTTAATCTTCATGCTGC |
| *GAPDH* | *GAPDH-F* | TCTCGATCTCAATTTCGCAAAA |
| | *GAPDH-R* | CGAAACCGTTGATTCCGATTC |

c. Lists of oligonucleotides used for the analysis of *nad1* transcript levels between wild-type and *msp1-1* mutant plants by RT-qPCR experiments.

| | | |
|---|---|---|
| *nad1a* | *ex-a_F* | TTGCCATATCTTCGCTAGGTG |
| | *ai1i_R* | AGTCTCATAGGCGACCGAAC |
| | *ai1ii_R* | CCTGCGGCACAAACCAATTT |
| *nad1b* | *bi1_F* | GACCAATAGATACTTCATAAGAGACCA |
| | *ex-b_R* | CGTGCTCGTACGGTTCATAG |

**Supplementary Table S2**. List of splicing factors in *Arabidopsis* and their mtRNA profiles.

| Gene | gene family | confirmed intron target(s)*[1] | mutant affected in *nad2* intron 1 | Ref's |
|---|---|---|---|---|
| *ABO5* | PPR | *nad2* i3 | (+)*[2] | (Liu *et al.*, 2010) |
| *ABO8* | | *nad4* i3 | (+) | (Yang *et al.*, 2014) |
| *BIR6* | | *nad7* i1 | yes | (Koprivova *et al.*, 2010) |
| *MID1* | | *nad2* i1 | - | (Zhao *et al.*, 2020) |
| *MISF26* | | *nad2* i3 | No | (Wang *et al.*, 2018) |
| *MISF68* | | *nad2* i2, *nad4* i1, *nad5* i4 | No | |
| *MISF74* | | *nad1* i4, *nad2* i4 | No | |
| *MTL1* | | *nad7* i2 | Yes | (Haili *et al.*, 2016) |
| *OTP43* | | *nad1* i1 | Yes | (Falcon de Longevialle *et al.*, 2007) |
| *OTP439* | | *nad2* i1 | - | (Colas des Francs-Small *et al.*, 2014) |
| *MSP1* | | *nad1* i1 | Yes | this study |
| *DEK2**[3] | | *nad1* i1 | Yes | (Qi *et al.*, 2017) |
| *TANG2* | | *nad5* i3 (possibly *nad5* i2) *[4] | Yes | (Colas des Francs-Small *et al.*, 2014) |
| *MatR* | Maturase | *nad1* i1 and many others | (+) | (Sultan *et al.*, 2016) |
| *nMAT1* | | *nad1* i1, *nad4* i2 | Yes | (Nakagawa & Sakurai, 2006; Keren *et al.*, 2012) |
| *nMAT2* | | *nad1* i2, *nad7* i2, *cox2* i1, and many other transcripts | Yes | (Keren *et al.*, 2009; Zmudjak *et al.*, 2017) |
| *nMAT3* | | *nad1* i3 (*nad1* i1) | Yes | (Shevtsov-Tal *et al.*, 2021) |
| *nMAT4* | | *nad1* i3 (*nad1* i1, *nad1* i4) | Yes | (Cohen *et al.*, 2014) |
| *ABO6* | Helicase | *nad1* i2, *nad1* i3, *nad5* i1 (*nad2* i2) | Yes | (He *et al.*, 2012) |
| *PMH2* | | Many organellar transcripts | Yes | (Köhler *et al.*, 2010; Zmudjak *et al.*, 2017) |
| *mCSF1* | CRM | Many organellar transcripts | Yes | (Zmudjak *et al.*, 2013) |
| *MTERF15* | mTERF | *nad2* i3 | No | (Hsu *et al.*, 2014) |
| *WTF9* | PORR | *ccmFc* i1, *rpl2* i1 | (+) | (Colas des Francs-Small *et al.*, 2012) |
| *ODB1* | RAD52-like | *nad1* i2 | Yes | (Gualberto *et al.*, 2015) |
| *OZ2* | RanBP2 | *nad1* i1, *nad2* i3, *nad5* i1, *nad5* i2, *nad7* i2, *rps3* i1 | Yes | (Bentolila *et al.*, 2021) |
| *RUG3* | RCC1-like | *nad2* introns 2 and 3 (*cox2* i1) | No | (Kühn *et al.*, 2011; Su *et al.*, 2017) |
| *UL18-L1* | UL18-Like | *nad5* i4 | Yes | (Wang *et al.*, 2020) |

| other mtRNA processing factors affecting *nad2* intron 1 splicing | | | | |
|---|---|---|---|---|
| MTSF1 | PPR | *nad4* stabilization | Yes | (Haili *et al.*, 2013) |
| SLO4 | | *nad4* editing | Yes | (Weissenberger *et al.*, 2017) |

*[1] - i, intron; *[2]- quotes indicate to putative intron targets; *[3]- *GRMZM2G110851 (DEK2)* gene encodes a putative maize ortholog of MSP1; *[4] - (+), indicates that the splicing of *nad2* intron 1 is affected to some degree in the mutant.

**Supplementary Table S3.** List of antibodies used for the analysis of Col-0 and *msp1* mutants.

| Antibody | Protein I.D. | origin | serum | dilution | Reference / source |
|---|---|---|---|---|---|
| Atp1 | Mitochondrial ATP-synthase subunit 1 | *Zea mays* | Mouse (monoclonal) | 1/500 | (Luethy *et al.*, 1993) |
| ATP2 | Mitochondrial ATP-synthase subunit 2 | *Zea mays* | Mouse (monoclonal) | 1/500 | (Luethy *et al.*, 1993) |
| CA2 | γ-carbonic anhydrase-like subunit 2 | *Arabidopsis thaliana* | Rabbit (polyclonal) | 1/1,000 | (Perales *et al.*, 2005; Sunderhaus *et al.*, 2006) |
| Cox2 | Cytochrome oxidase subunit-2 | *Arabidopsis thaliana* | Rabbit (polyclonal) | 1/5,000 | Agrisera antibodies, AS04 053A |
| Nad1 | NADH-dehydrogenase complex subunit-1 | *Arabidopsis thaliana* | Rabbit (polyclonal) | 1/1000 | Gift of Dr. Etienne Meyer, Halle U. |
| Nad9 | NADH-dehydrogenase complex subunit-9 | *Triticum spp.* | Rabbit (polyclonal) | 1/50,000 | (Lamattina *et al.*, 1993) |
| RISP | Rieske iron-sulfur protein | *Arabidopsis thaliana* | Rabbit (polyclonal) | 1/5,000 | Gift of Prof. Ian Small, UWA |
| VDAC | Mitochondrial membrane-associated β–barrel proteins | *Zea mays* | Mouse (monoclonal) | 1/1000 | Thomas Elthon collection, PM035 |
| AOX1/2 | Alternative oxidase | *Sauromatum guttatum* | Rabbit (polyclonal) | 1/500 | Agrisera antibodies, AS04 054 |

**Supplementary Table S4**.

**A.** Relative accumulation of mitochondrial proteins in wild-type and *msp1* plants.

| | ***msp1-1*** **(hmz)** | ***msp1-1*** **(htz)** | **Col-0-heart** | ***msp1*****/35S:MSP1** |
|---|---|---|---|---|
| **Complex I** | | | | |
| CA2 | 3.7 ± 0.5 | 2.3 ± 0.6 | 2.2 ± 1.2 | 1.3 ± 0.5 |
| Nad1 | LoD* | 1.2 ± 0.2 | 0.9 ± 0.7 | 1.4 ± 0.3 |
| Nad9 | 2.1 ± 0.3 | 0.4 ± 0.3 | 1.8 ± 0.6 | 0.7 ± 0.2 |
| **Complex III** | | | | |
| RISP | 6.2 ± 0.7 | 1.1 ± 0.4 | 2.3 ± 0.2 | 1.4 ± 0.5 |
| **Complex IV** | | | | |
| Cox2 | 3.1 ± 0.8 | 1.4 ± 0.3 | N.D.** | 1.3 ± 0.4 |
| **Complex V (ATP synthase)** | | | | |
| Atp1 | 3.6 ± 1.1 | 1.1 ± 0.2 | 1.6 ± 0.5 | 1.3 ± 0.4 |
| ATP2 | 1.3 ± 0.7 | 2.0 ± 0.4 | 1.2 ± 0.3 | 2.2 ± 0.7 |
| AOX1/2 | 3.4 ± 0.9 | 1.0 ± 0.3 | N.D.** | 1.9 ± 0.7 |
| VDAC | 4.3 ± 1.3 | 2.1 ± 0.5 | N.D.** | 1.9 ± 0.7 |

* - LoD - below limit of detection

**B.** Relative accumulation of native organellar respiratory complexes in wild-type and *msp1* plants.

| **Genetic line** | **holo-CI** | | $\mathbf{CIII_2}$ | **CIV** | **CV** |
|---|---|---|---|---|---|
| | CA2 | Nad9 | | | |
| **Col-0** | 1.00 ± 0.13 | 1.00 ± 0.07 | 1.00 ± 0.16 | 1.00 ± 20.4 | 1.00 ± 0.4 |
| **Col-0-heart** | 2.35 ± 0.22 | 2.57 ± 0.15 | 1.24 ± 0.09 | 0.54 ± 0.4 | 1.14 ± 0.8 |
| ***msp1-1 (htz)*** | 2.07 ± 0.14 | 0.91 ± 0.09 | 1.15 ± 0.12 | 0.67 ± 0.12 | 1.02 ± 0.03 |
| ***msp1-1 (hmz)*** | LoD* | LoD* | 3.37 ± 1.04 | 5.37 ± 1.21 | 3.34 ± 0.21 |
| ***msp1*****/35S:MSP1** | 2.21 ± 0.23 | 1.14 ± 0.21 | 2.45 ± 0.16 | 2.85 ± 0.55 | 1.54 ± 0.17 |

* - LoD - below limit of detection

**Supplementary Table S5.** Oligonucleotides used in screening of individual T-DNA insertion lines in Arabidopsis and cloning of the *MSP1-GFP* gene-fusion construct.

| Gene target | Gene I.D. | oligo name | Oligonucleotide sequence (5'-to-3') |
|---|---|---|---|
| ***MSP1*** | At4g20090 | MSP1-1-F1 | CTCCCAAAATGGGTTCTTTC |
| | | MSP1-1-F2 | GAATGACACATGTAAACATCTG |
| | | MSP1-1-F3 | TCAATTAATTGAGAGGAAACAAA |
| | | MSP1-1-R1 | CATCAGGCAAACACTTTCTCT |
| ***ACTIN2*** | At3g18780 | Act2-F | TCTTCCGCTCTTTCTTTCCAAG |
| | | Act2-R | CTGGCGTACAAGGAGAGAAC |
| **SALK T-DNA right border** | pROK2 T-DNA | RB-10064 | CCAGATCCGGTGCAGATTATTTG |
| **SALK T-DNA left border** | pROK2 T-DNA | LB-6443 | CATCGCCCTGATAGACGGTT |
| **SALK T-DNA left border** | pROK2 T-DNA | LBb1.3 | ATTTTGCCGATTTCGGAAC |
| ***MSP1-GFP-1*** | pSAT6-eGFP-N1 cloning | MSP1-1-GFP-F | ACATAGCCATGGACATGCCCAAATGCCCAATCCC |
| | | MSP1-1-GFP-R | CCATCAGGATCCCTCTAAAGACAACAATGAAACTA |
| ***MSP1-GFP-2*** | pCAMBIA Cloning | PF_SAT6-EcoRI | ACATAGGAATTCTTCCCAGTCACGACGTTGTA |
| | | PR_SAT6-SmaI | ACATAGCCCGGGCAGCTATGACCATGATTACTG |
| ***35S-MSP1-1*** | pSAT6-35S-Promoter cloning | MSP1-1-COMP-F | ACATAGCCATGGACATGCCCAAATGCCCAATCCC |
| | | MSP1-1-COMP-R | CCATCAACTAGTTCAAGTACACAGGTTCCTCC |
| ***35S-MSP1-2*** | pCAMBIA Cloning | PF_SAT6-SmaI | ACATAGCCCGGGTTCCCAGTCACGACGTTGTA |
| | | PR_SAT6-SalI | ACATAGGTCGACCAGCTATGACCATGATTACTG |
