## Supplemental Materials and Methods for "*MSP1* encodes an essential RNA-binding PPR factor required for *nad1* maturation and complex I biogenesis in *Arabidopsis* mitochondria"

### Supplemental Data S2. Detailed Materials and Methods

#### Plant material and growth conditions

*Arabidopsis thaliana* (ecotype Columbia-0) was used in all experiments. Wild-type (Col-0) and *mps1* mutant line SALK-142675 (*msp1-1*), SALK-018927 and SALK-070654 were obtained from ABRC, at Ohio State University. The growth conditions used for the analysis of wild-type plants and mutant lines were identical to those described previously (Shevtsov-Tal *et al.*, 2021). PCR was used to screen the plant collection and check the insertion integrity of each individual line. Sequencing of PCR products was used to analyze the precise insertion site in the T-DNA lines.

#### GFP localization assays

*In-vivo* protein localization by GFP fusion were performed via root transformation in tissue culture, and as indicated previously (Weigel & Glazebrook, 2006). The N-terminal region (300 bps) of *MSP1* (*AT4G20090*) gene was fused in-frame to GFP coding sequence and cloned into a modified pCAMBIA binary vector (gift of Dr. Yoram Eyal, ARO). The vector was introduced into *A. tumefaciens* (strain GV3101), and then used to infect *Arabidopsis* root culture cells. After 48 hours, the infected cells were analyzed by confocal microscopy at the Bio-Imaging unit (Institute of Life Sciences, The Hebrew University of Jerusalem, Jerusalem Israel). Mitochondria were visualized by MitoTracker, as described previously (Nguyen *et al.*, 2022).

#### Embryo rescue and establishment of homozygous *MSP1* mutants

The development of *msp1* mutant-line embryos was performed essentially as described previously (Shevtsov-Tal *et al.*, 2021), using white seeds obtained from green mature siliques of wild-type or heterozygous *msp1-1* (SALK-line 142675) plants.

#### Plant transformation and functional complementation of the *MSP1-1* mutation

A 1,989 bp fragment containing the coding region of *MSP1* was generated by PCR, cloned under the control of the 35S promoter region and cloned into a modified pCAMBIA vector (gift of Dr. Yoram Eyal, ARO). The shuttle vector containing the transgene was electroporated into *A. tumefaciens* (strain GV3101), that was used to transform *msp1-1* plants by the floral dip method (Clough & Bent, 1998).

**Microscopic and macroscopic analyses of *Arabidopsis* wild-type and mutant plants**

Analysis of whole plant morphology, roots, leaves, siliques and seeds of wild-type and mutant lines were examined under a stereoscopic (dissecting) microscope or light microscope at the Bio-Imaging unit of the Institute of Life Sciences (The Hebrew University of Jerusalem, Israel).

**RNA extraction and analysis**

RNA extraction and analysis were performed essentially as described previously (Sultan *et al.*, 2016; Shevtsov-Tal *et al.*, 2021). qPCR reactions were run on a LightCycler 480 (Roche), using the LightCycler 480 SYBR Green I Master mix in a final volume of 5 µL. *GAPDH* (*AT1G13440*), *ACTIN2* (*AT3G1878*), 18S rRNA gene (*AT3G41768*), *rrn26* (*i.e.*, mitochondrial 26S rRNA gene, *ATMG00020*), *rrn5* (*ATMG01380*) and *rrn18* (*ATMG01390*), were used as reference genes for normalization in the qPCR analyses.

**Transcriptome mapping by high-throughput (RNA-seq) analysis**

Total plant RNA was extracted from *Arabidopsis* wild-type and mutant lines. Sequencing was carried out by IDT (Synthezza, Jerusalem, Israel), with Illumina NovaSeq 6000 platform (Novogene Corporation, China), used for a paired-end 150 bp sequencing strategy. Initial read-quality filtering was performed with FastQC Version 0.11.9 (Babraham Bioinformatics). Reads were filtered for low quality and primer or adapter contamination using BBduk from the BBtools package (Joint Genome Institute). Filtered reads were mapped to the updated *Arabidopsis* mitogenome (BK010421) (Sloan *et al.*, 2018), using STAR version 2.7.9a (version 2.2.9) (Dobin *et al.*, 2013) and BBMAP (https://sourceforge.net/projects/bbmap; v38.92). Sequences are available at the Sequence Read Archive (BioProjects PRJNA704631 and PRJNA768306).

**Preparation of recombinant MSP1 protein**

A g-Block fragment (Synthezza, Jerusalem, Israel) containing the MSP1s' PPR domains 4 to 14 was cloned into pGEX-4T1, in-frame to GST in its N-terminus and a C-terminal 6xHis tag. The recombinant protein (rMSP1) was expressed in *E. coli* strain BL21 cells, grown at 37°C to $OD_{600}$ of 0.6, and induced with 1.0 mM IPTG for 2 h at 37°C. The cells were pelleted, resuspended with His purification buffer (Bio Rad) supplemented with 6M guanidine hydrochloride and lysed by a high-pressure chamber manifold (Constant systems, Northants, UK). The recombinant GSTrMSP1-His protein was purified by affinity chromatography (IMAC) and dialyzed against 50% glycerol buffer (10mM Tris-HCl pH7, 500 mM NaCl, 1% Triton X-100, and 5mM βME).

#### ***In-vitro* RNA binding assays**

*In-vitro* transcription of RNA was carried out with T7-RNA polymerase in the presence of an RNase-inhibitor. Following DNase digestion, phenol extraction and ethanol precipitation, 5' labeling of RNAs was done with [γ-$^{32}$P]-ATP and T4 PNK. Non-radioactive, body-labeled RNAs were obtained with N6-biotinlated C/UTP ribonucleotides. Filter-binding assays were performed with a slot-blot manifold (Ostersetzer *et al.*, 2005; Barkan *et al.*, 2007; Keren *et al.*, 2008). RNAs were cross-linked to the membranes by UV-illumination, and detection was carried out by immunoblotting (biotinylated RNAs) or laser-scanner platform (, Typhoon, $^{32}$P-labeled RNAs). The fraction of bound RNA was calculated as the ratio between the signal of the RNA captured by the protein (upper-nitrocellulose membrane) and total RNA signals (upper and lower charged-membrane).

#### **Exonuclease and endonuclease protection assays**

The PNPase protection assay was perform essentially as described by Prikryl et al. (2011). The radiolabeled RNA was heated for 2 min at 95 °C and snap cooled on ice, before was used in the binding reactions (10 µL). In each reaction, 1 µL of recombinant PNPase, RNase-A or Benzonase, and samples were incubated at 25°C for 15 min. The samples were denatured by heating in 60% formamide and resolved on 10% AA/7M urea gels, 1×TBE buffer.

#### **Enriched mitochondrial membrane preparation**

Crude mitochondrial membranes were prepared from 200 mg of liquid MS-grown plantlets, essentially as described previously (Pineau *et al.*, 2008). Protein concentration was determined by the Bradford method (BioRad) according to the manufacturer's protocol, with bovine serum albumin (BSA) used as a calibrator.

#### **Plant protein extraction and analysis**

Immunoblotting was performed essentially as described previously (Nguyen *et al.*, 2022), with aliquot equivalent to 10 mg crude mitochondria extract. Detection was carried out by chemiluminescence assays after incubation with an appropriate secondary antibody. The relative accumulation of proteins was estimated from their relative intensities by densitometry measurements and quantifications using the ImageJ software.

**Blue native (BN) electrophoresis for isolation of native organellar complexes**

Blue native (BN)-PAGE of crude organellar membrane fractions was performed generally according to the methods described by Pineau *et al.* (2008) and Nguyen *et al.* (2022). An aliquot equivalent to 40 mg of crude *Arabidopsis* mitochondrial extracts, obtained from wild-type and *MSP1* plants was solubilized with 5% (w/v) digitonin in BN solubilization buffer (Shevtsov-Tal *et al.*, 2021), and loaded onto a native 4-16% linear gradient gel. For immunoblotting, the gel was transferred to a PVDF membrane. CI activity assays were performed essentially as described previously (Eubel *et al.*, 2005).
